# Stress-induced DNA methylome plasticity and transcriptional re-programming in *Staphylococcus aureus*

**DOI:** 10.64898/2026.05.22.727101

**Authors:** Luke B. Jones, Maisem Laabei, Stefan Bagby

**Author notes:** Corresponding author:Email addresses: LBJ -; ML -; SB.

## Abstract

*Staphylococcus aureus*, a major human and livestock pathogen, is the second biggest cause of antimicrobial resistance-associated mortality. Although *S. aureus* transcriptional regulation has been extensively characterised, the potential role of DNA methylation in *S. aureus* transcriptional regulation and stress response remains largely undefined. We tackled this gap by combining genome-wide nanopore sequencing-derived DNA methylation data and transcriptomic data, acquired before and during exposure of methicillin-resistant *Staphylococcus aureus* strain USA300 to clinically relevant oxidative, antibiotic and nitrosative stresses. Stress-induced significant DNA methylation changes were rare, with <0.1% of cytosines/adenines undergoing ≥20% change in methylation; these changes were enriched within genomic features (protein-coding genes, predicted promoter regions, ncRNAs). Transcription changes reflected metabolic, regulatory and stress-specific pathway adjustments. Many of the stress-induced DNA methylation changes occurred alongside transcription changes, although there was no obvious overarching relationship between the directions of changes in DNA methylation and transcription. Pre-stress treatment methylation entropy tended to be elevated at sites containing a base that underwent stress-induced change in methylation level, identifying focal sites of pre-existing methylome heterogeneity relative to both local and genome-wide backgrounds; the magnitude of the site-specific methylation entropy peak, moreover, correlated with the magnitude of subsequent methylation change. Genome-wide entropy levels were consistent with a finite number of methylation patterns, distinguished by key bases, that could correspond to clinically important *S. aureus* subpopulations exhibiting persistence, dormancy and immune evasion. These findings support the principle that DNA methylation is an important component of the regulatory machinery underpinning *S. aureus* adaptability and persistence.

**Importance:** *Staphylococcus aureus* survives antibiotic treatment and host immune attack partly through phenotypic diversity, yet the regulatory processes that support this adaptability remain incompletely understood. Here, we integrate genome-wide DNA methylation and transcriptomic profiling to examine how epidemic methicillin-resistant *S. aureus* USA300 responds to oxidative, antibiotic, and nitrosative stresses. Stress-induced methylation changes were rare but non-random, concentrated in coding and regulatory regions linked to stress defence, metabolism, persistence, virulence, biofilm formation, and host interaction. Sites that underwent stress-induced methylation changes tended to have unusually high pre-stress methylation pattern diversity relative to local and genome-wide pattern diversity. These findings support a model in which an *S. aureus* population contains a pre-stress repertoire of epigenetic states that stress selectively redistributes via bases that differentiate stable methylation patterns. By revealing a layer of heterogeneity that may support pathogen resilience, this work provides a framework for investigating whether bacterial methylation dynamics can be exploited clinically.

## Introduction

*Staphylococcus aureus* is a challenging opportunistic bacterial pathogen worldwide: it is one of the six ESKAPE multidrug-resistant pathogens and is the primary cause of bloodstream, skin, and device-associated infections [1]. A 2019 survey of global antimicrobial resistance (AMR) burden showed that *S. aureus* is the second-leading pathogen in AMR-associated mortality, and extensions to this work showed that methicillin-resistant *S. aureus* (MRSA) was the fastest increasing contributor to AMR-associated mortality between 1990 and 2020 [2,3]. As such, *S. aureus* is a substantial clinical and economic burden.

Phenotypic plasticity is a key *S. aureus* survival characteristic, enabling formation of specialised subpopulations such as persister cells and small colony variants (SCVs) that can withstand hostile environments. These subpopulations often survive potent antibiotic and immunological assaults, subsequently causing relapses weeks or months later [4]. Kracalik *et al*., for example, observed recurrence rates of 16% in US patients with MRSA, up to 61% within 180 days [5]. This adaptive capability is believed to contribute to persistent infections like osteomyelitis, endocarditis, and prosthetic joint infections.

The USA300 lineage exemplifies these *S. aureus* characteristics: it can persist long-term within hosts, survive inside cells, and recur following treatment, and is therefore an ideal model for studying stress survival, phenotypic variability, and chronic infection. Identified in San Francisco in 2000, USA300 quickly became the dominant community-associated MRSA clone across USA and parts of Europe. Its genome contains numerous mobile genetic elements, including SCCmec IV, ACME, and PVL, which together enhance skin colonisation, virulence, immune evasion, and transmission [6].

Transcriptional regulation in *S. aureus* has been extensively characterised, including the roles of transcription factors, sigma factors and transcriptional regulatory networks [7,8], two-component signal transduction systems such as the *agr* quorum sensing locus [9], and RNA regulators [10]. The role of DNA methylation in *S. aureus* transcriptional regulation, however, merits further investigation. All three forms of DNA methylation, N6-methyladenine (6mA), N4-methylcytosine (4mC) and C5-methylcytosine (5mC), are found in bacteria. Bacterial DNA methylation affects gene expression through various mechanisms such as altering DNA topology, and DNA-transcription factor/RNA polymerase interactions [11–14]. In several bacterial pathogens, such as *Escherichia coli*, *Salmonella enterica*, *Pseudomonas syringae*, and *Helicobacter pylori*, DNA methylation regulates gene expression in response to environmental changes [14–16]. Moreover, DNA methylome heterogeneity can promote phenotypic diversification within clonal populations, resulting in subpopulations that can display enhanced antibiotic resistance or altered virulence expression linked to recurrent infections [14,17]. This underscores the importance of decoding the DNA methylome to understanding of bacterial adaptation, but only a few studies have examined these effects in *S*.

*aureus*. One recent study connected changes in methylation with persistence phenotypes in clinical MRSA strains from neonatal intensive care outbreaks, implying a role for reversible, heritable epigenetic changes [18]. Korotetskiy *et al*. found that hospital-associated *S. aureus* isolates exhibit “clonal gene expression stability”, in which core gene expression patterns remain stable across sublethal antimicrobial stress conditions [19]. They further demonstrated that stable gene expression patterns align with strain-specific DNA methylation signatures, suggesting an epigenetic basis for transcriptional regulation. Notably, while global methylation patterns remained consistent, differences in single base modification frequencies, particularly near transcription start sites, were statistically associated with strain-specific gene expression patterns. Thus, *S. aureus* has inheritable, methylation-associated gene expression programmes that are both stable and epigenetically distinct across lineages.

To improve understanding of the contribution of methylation to *S. aureus* adaptation to clinically relevant stress conditions, we have characterised the DNA methylome and the transcriptome of *S. aureus* USA300 following exposure to oxidative (hydrogen peroxide; H2O2), antibiotic (methicillin) and nitrosative (S-nitroso-N-acetylpenicillamine; SNAP/NO) treatments. *S. aureus* is subjected to oxidative and nitrosative stresses in numerous host contexts. Neutrophils, for example, are rapidly recruited to infection sites such as skin abscesses, bloodstream and wounds where neutrophils can unleash a potent oxidative burst comprising superoxide, hydrogen peroxide, hypochlorous acid and nitric oxide [20]. During phagocytosis of *S. aureus* cells, moreover, macrophages generate reactive oxygen and reactive nitrogen species that exert a substantial impact on the *S. aureus* transcriptome [21,22]. Even during asymptomatic colonisation of aerobic niches such as skin and nasal passages, *S. aureus* cells are exposed to oxidative stress due to aerobic respiration and from antimicrobial oxidants in sweat and sebaceous secretions [23].

Using Oxford Nanopore Technologies and Illumina sequencing platforms, we have generated base resolution maps of DNA methylation and combined them with differential gene expression data to examine whether methylation changes are feature-specific, and whether methylation changes correlate with oxidative, antibiotic and nitrosative stress-induced transcriptional reprogramming. Our findings reveal non-random, stress-specific methylation changes concentrated in upstream and gene coding regions of differentially expressed genes, supporting a model of targeted epigenetic regulation. This is consistent with the possibility that methylation provides a heritable mechanism contributing to the formation of stress-adapted subpopulations. As such, DNA methylation could be a key part of the apparatus that enables *S. aureus* USA300 to maintain phenotypic diversity, persist under hostile conditions, and remain poised for recurrence without requiring genetic change.

## Materials and methods

### Stress induction and sampling

To identify optimal stress conditions for transcriptomic analysis, *Staphylococcus aureus* subsp. aureus strain USA300-FPR3757 (LAC) [6] was grown to mid-exponential phase (OD600 = 0.7) in tryptic soy broth (TSB; Merck), and subcultured (1:100) into stressor cultures. Survival assays lasted three hours (37 °C, 200 rpm), with sampling at 20-minute intervals for colony forming unit (CFU) enumeration on agar (Thermo Scientific). Conditions were considered optimal when there was a 30–90% decrease in CFUs compared to untreated controls between 30 and 90 minutes, followed by recovery to below 50% of the original CFU count by 180 minutes. These parameters were selected to maximise physiological stress, ensuring a robust transcriptional response without triggering large scale, irreversible cell death. Based on these results, all samples for DNA and RNA extraction were collected at the 60-minute time point, corresponding to maximal perturbation. At this time point, CFU levels across treatments typically showed a 20–30% decrease, consistent with the onset of significant stress while remaining within the predefined optimal window. Final working concentrations were: 2.25 mM hydrogen peroxide (H O; Thermo Scientific) for oxidative stress, 150 μg/mL methicillin (Merck) for antibiotic stress, and 400 μM S-nitroso-N-acetylpenicillamine (SNAP; Merck) for nitrosative stress, applied in 1 L cultures. DNA and RNA were extracted according to the manufacturer’s protocol [24], with bead-beating settings optimised for long sequencing reads (4 × 30 s cycles at 8,000 rpm).

### DNA sequencing and genome assembly

*Staphylococcus aureus* subsp. aureus strain USA300-FPR3757 (LAC) genomic DNA was extracted from three replicates of pre-treatment and post-treatment for each of H□O□, methicillin, and SNAP, making a total of eighteen samples (nine pre-treatment and nine post-treatment). Each of these was sequenced on a MinION (Oxford Nanopore Technologies, ONT), following the Native Barcoding Kit 24 V14 protocol (ONT). Basecalling was performed with Dorado (v0.9.0; dna_r10.4.1_e8.2_400bps_sup@v5.0.0).

The reads from the nine pre-treatment samples were merged for genome sequence assembly. Assembly was carried out using Flye (v2.9.2) [25] in a two-step approach. First, reads longer than 1 kb with quality scores above Q38 were used to generate high confidence contigs at approximately 2477× coverage. These were scaffolded using reads above 5 kb with Q20 or better, with 15% base quality lenience to enrich repetitive genomic regions, leading to 6000× coverage. This approach produced three circular contigs. Medaka (v2.0.1) [26] was used for consensus polishing. Read quality filtering was done with fastp (v0.24.0) [27]. Assembly quality was verified using CheckM [28] and QUAST [29], benchmarked against the USA300 ISMMS1 reference genome (GenBank: GCA_000568455.1).

### RNA extraction, library preparation, and sequencing

*Staphylococcus aureus* subsp. aureus strain USA300-FPR3757 (LAC) total RNA was extracted from three replicates of pre-treatment and post-treatment for each of H□O□, methicillin, and SNAP, making a total of eighteen samples (nine pre-treatment and nine post-treatment). Each sample was assessed for integrity using Agilent TapeStation high sensitivity analysis. Only samples with RNA Integrity Number above nine were pursued further. Ribosomal RNA was depleted using the Ribo-off rRNA Depletion Kit V2 for bacteria (N417, Vazyme, China). Library prep was performed using ALFA-SEQ Directional RNA Lib Prep Kit (Findrop). Libraries were sequenced on an Illumina NovaSeq X Plus platform in paired-end 150 bp mode (PE150; BMKGENE, UK).

### Transcriptome analysis

RNA-seq reads were first trimmed using cutadapt (v4.4) [30], aligned to the assembled genome with HISAT2 (v2.2.1) [31], and sorted using SAMtools (v1.12) [32]. Gene expression was quantified at the gene level using featureCounts (v2.0.8) [33], and differential expression analysis was performed with DESeq2 (v1.42.1) [34]. Functional enrichment analysis of differentially expressed genes was carried out using KEGG pathway analysis through the clusterProfiler package (v4.15.1) [35].

### Genomic feature prediction

Open reading frames (ORFs) were predicted using PGAP [36]. Genes which were not annotated by PGAP (including *katA* and *rot*) were identified using BLAST+ (v2.16.0) [37]. Operon structures were predicted using Operon-mapper [38]. Upstream regulatory regions were defined as sequences spanning 20–700 bp upstream of each operon, truncated if overlapping with same-strand genes or within 50 bp of another upstream promoter. Terminators were defined as 5–200 bp regions located downstream of operons, extending for either 200 bp or until the start of a downstream gene on the same strand, with a maximum allowed overlap of 10 bp. Non-coding RNAs (ncRNA) were identified using cmsearch (v1.1.5) [39].

Throughout this study, “genomic features” refers collectively to annotated protein-coding genes, predicted promoter regions, and ncRNAs, i.e., discrete, functionally annotated elements of the genome. Horizontally acquired and mobile genetic elements were identified using a combination of AlienHunter (v1.7.7) [40], ICEberg3 (v3.0) [41], MobileOG-db (v1.1.3) [42], PHASTEST (v3.0) [43], and Phigaro (v2.4) [44], providing comprehensive coverage of integrated elements such as integrative and conjugative elements (ICEs), transposons, prophages, and genomic islands.

### Methylation detection and analysis

Methylation data generated from Nanopore sequencing were processed using Dorado, SAMtools, and Modkit (v0.4.2) [45] to produce BAM and BedMethyl files representing genome-wide methylation calls *per* pre-and post-treatment repeat. Calling of base modifications was performed using super-accuracy models in Dorado. These models have been benchmarked on synthetic ground-truth datasets and are widely used for comparative methylation analyses [46]; independent studies comparing Nanopore-derived methylation calls to previous gold standard approaches support the utility and reliability of Nanopore-derived methylation calls [47–49]. These files were then converted to base-resolution BED format using SeqKit (v2.4.0) [50] and Modkit. The associated output reports the percentage of reads in which a particular base is methylated, which we term the population level (PL) of that base; ΔPL is the pre-and post-treatment difference in PL. A Python script was used to intersect methylation events across the genome with annotated genomic features, allowing identification of regulatory hotspots (Supplementary Table 1). Methylation changes were assessed for each set of three stressor replicates compared to the corresponding pre-treatment replicates. In addition, BedMethyl files were used with Modkit to calculate methylation entropy (ME) for each annotated gene-coding region and each defined upstream region, with default settings.

## Statistical analysis

Enrichment of significant (≥ 20% |ΔPL|; threshold justified in Results section “Genome-Level Trends in Methylation”) stress-induced base methylation changes within annotated genomic features was assessed using a one-tailed hypergeometric test. For each sample, all adenine/cytosine genomic positions, including those that underwent significant methylation changes, were mapped to annotated and unannotated regions in a strand-aware (strand in question) manner and separately in a strand-blind (both strands) manner. Counts of overlapping bases were used to compute enrichment relative to a defined genomic background. Enrichment *p*-values were adjusted for multiple comparisons using the Benjamini–Hochberg false discovery rate (FDR; *q*-value) method. This analysis was implemented in Python using genomic interval matching and standard statistical libraries.

To assess the relationship between stress responsive sites and pre-existing methylation heterogeneity, pretreatment Modkit 50bp methylation entropy windows supported by ≥10 reads were analysed. Site-overlapping ME was defined as the support-weighted median entropy of windows overlapping each callable base (sites here being bases which later reach |ΔPL| ≥0.20). Inverse sampling-weighted linear models tested whether site-overlapping ME predicted subsequent |ΔPL| across the callable genomic risk set, adjusting for baseline PL, sequencing depth, ME support, local GC content, target base density, number of overlapping windows, treatment callability and treatment identity. For sites with |ΔPL| ≥0.20, focal ME enrichment was assessed relative to supported non-overlapping windows within ±500 bp and to eligible callable positions matched by modification, contig and strand across the genome. Confidence intervals and *p*-values were calculated using 5-kb genomic-block CR1 robust standard errors, with Holm correction within each pre-specified test family.

To examine whether changes in methylation entropy were associated with differential gene expression, we binned genes into percentile groups based on the absolute z-score of methylation entropy change (|(ΔME)|) and assessed their relationship with absolute log fold change (|log FC|) values from the transcriptome analysis. Ordinary least squares (OLS) regression was performed separately for each percentile bin (50th, 75th, and 95th), using |(ΔME)| as the independent variable and |log FC| as the dependent variable. Analyses were stratified by strand orientation (same vs. complementary) based on PGAP gene annotations (39). For each bin, we reported the OLS slope (β), R², and two-tailed *p*-value, along with non-parametric 95% confidence intervals for β and R² estimated by bootstrap resampling (B = 500). We also assessed null distributions of β under permutation by shuffling |(ΔME)| values (1,000 iterations) and plotted the resulting histograms to visualise deviation from the null. Pseudo-replicates, induced by multigene operons sharing promoter or terminator regions, were dereplicated using an operon-wide mean |log FC|. Scatter plots were generated with bin-specific colouring and point transparency, with shaded regions indicating bootstrap confidence intervals of OLS fits. All analyses and visualisations were implemented in Python.

## Results

### Genome sequence of S. *aureus* USA300

Nanopore sequencing of DNA from the nine pre-treatment and nine post-treatment samples yielded a combined total of 11,807,091 reads. Of these, 59.8% had a Phred quality score above Q20, with read lengths ranging from 1 kb to 1.06 Mb. The genome sequence assembly using the combined reads from the nine pretreatment samples comprised a 2,874,399 bp chromosome and two plasmids (27,066 bp and 3,125 bp); all contigs were circular and NCBI BLASTn analysis confirmed that both of the plasmids matched previously described *S. aureus* plasmids. The genome sequence assembly showed 97.99% completeness and 0.24% contamination according to CheckM [28] analysis, indicative of an accurate and nearly complete assembly. Annotation with PGAP identified 2,843 ORFs, including 2,757 protein-coding genes [36]. QUAST [29] analysis showed that all sequencing reads (pre-treatment and post-treatment) aligned back to the assembly. The assembly covered 98.8% of the USA300 ISMMS1 reference genome, had comparable GC content (32.8%), and differed structurally at four regions ≥500 bp in size. Assembly coverage was consistent across contigs, ranging between 250x and 750x *per* replicate, with few suspicious drops in coverage (Supplementary Figure 1). These results establish that the assembled genome sequence is highly complete with very deep coverage, biologically representative, and suitable for downstream methylation and expression analysis.

### Transcriptomic validation of hydrogen peroxide, methicillin and S-nitroso-N-acetylpenicillamine stress-specific responses

Illumina RNA sequencing of eighteen samples (three replicates of pre-treatment and post-treatment for each of H□O□, methicillin, and SNAP) generated 342,899,554 paired-end reads, averaging ∼19 million *per* sample; this constitutes ultra-deep transcriptomic resolution [51]. Global transcriptional profiling revealed statistically significant, condition-specific responses at the individual gene level (Figure 1), whereas gene set enrichment analysis (GSEA) assessed coordinated expression shifts across complete ranked gene lists, including genes that were not individually significant (Figure 2). Differences in pathway membership and gene set composition can therefore produce apparent discrepancies between the gene-level and pathway-level profiles; these pathway-specific expression distributions are shown in Supplementary Figure 2.

**Figure 1.**
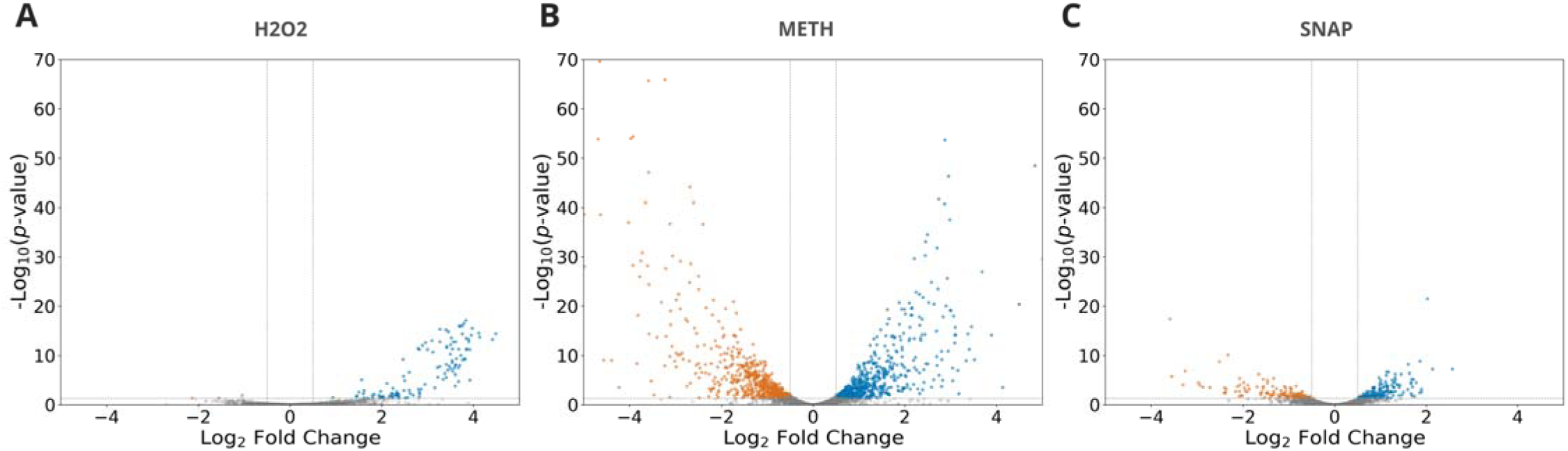
Volcano plots of differential gene expression. DESeq2 results for stress treatments: (A) H₂O₂, (B) methicillin, (C) SNAP. Genes shown in grey have insignificant adjusted p-values and/or showed | log_2_ fold change| < 0.5, while genes shown in blue and orange exhibited significant expression increase or decrease, respectively. In order to accommodate a common y-axis that enables direct comparison between panels, 10 highly significant methicillin-responsive genes with –log₁₀(p) > 70 fell outside the plotting range and are not shown. These include: blaZ (ACN-SPI_RS00105; log₂FC = +6.33), rlmD (ACNSPI_RS06510; log₂FC = +5.25), vraX (ACNSPI_RS13585; log₂FC = +5.54), DUF6583 family protein (ACNSPI_RS00725; log₂FC = −6.39), hypothetical protein (ACNSPI_RS03695; log₂FC = +5.01), pyrP (ACNSPI_RS10670; log₂FC = −4.13), hypothetical protein (ACNSPI_RS11520; log₂FC = −5.04), tRNA-Asn (ACN-SPI_RS11585; log₂FC = +5.24), and hypothetical protein (ACNSPI_RS13590; log₂FC = +5.05). All data points for H₂O₂ and SNAP are displayed.

**Figure 2.**
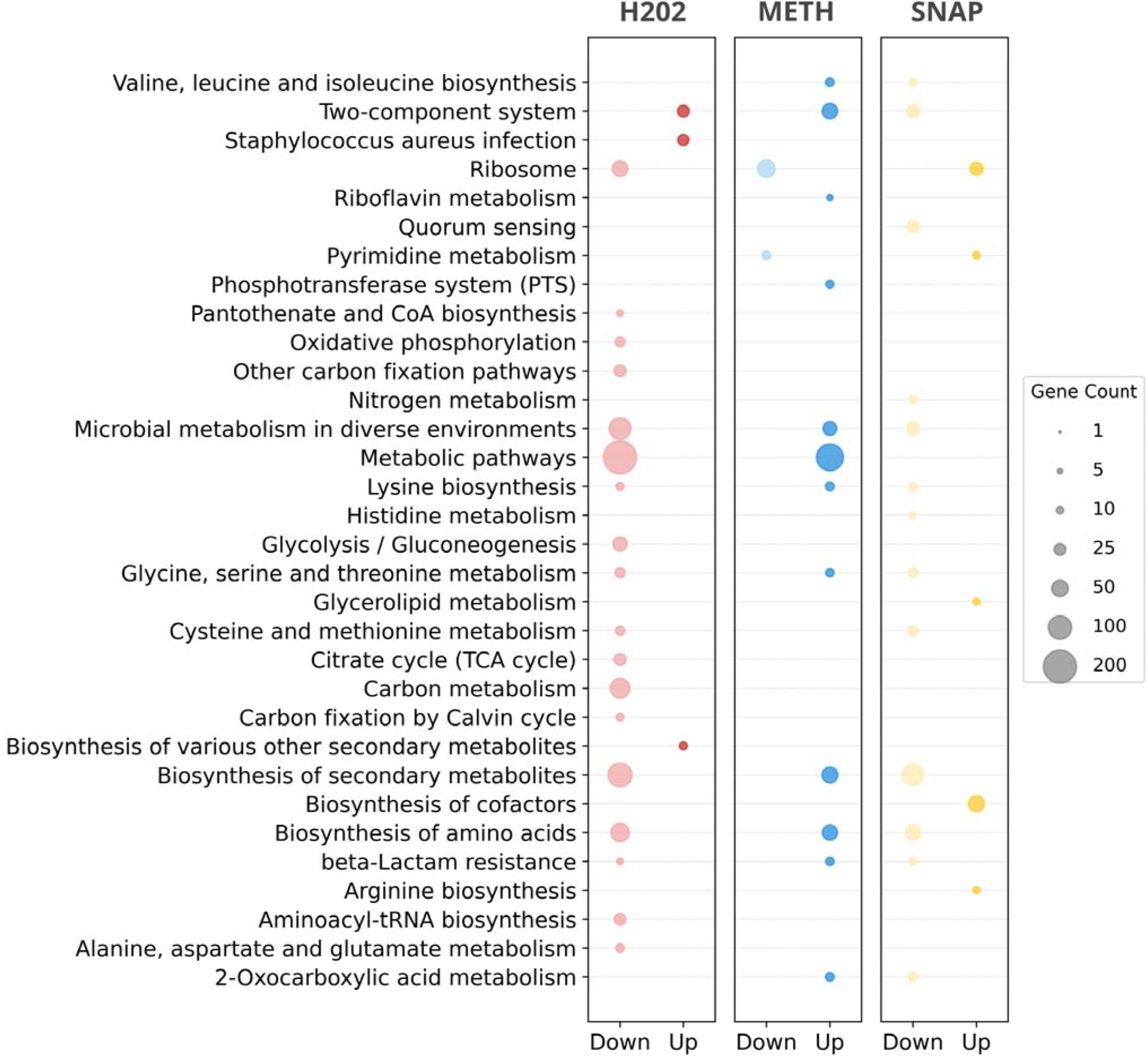
KEGG pathway enrichment and KEGG orthology-assigned gene expression distributions. Gene set enrichment analysis (GSEA) was performed using all genes ranked by log₂FC, without applying a significance threshold. Pathway enrichment is based on the normalised enrichment score, indicating directionali ty, with circle size reflecting gene set size. Plots for each treatment (H₂O₂, methicillin, SNAP) are separated into downregulated and upregulated axes, and coloured by treatment. GSEA was performed using complete ranked gene li sts; pathway enrichment direction and bubble number should therefore not be interpreted as direct measures of the numbers of significantly upregulated or downregulated genes.

121 genes were significantly differentially expressed upon H□O□ treatment, comprising 118 upregulated genes and 3 downregulated genes (Figure 1). H□O□ triggered a broad oxidative and genotoxic stress response. Strongly induced genes included a prophage-associated cluster containing phage replication, recombination, structural and lysis-related functions, consistent with prophage activation following DNA damage and providing a sensitive transcriptional indicator of peroxide-associated stress in lysogenic *S. aureus.* H□O□ also induced the high affinity phosphate-uptake genes *pstC*, *pstA* and *pstB* (log□FC = +1.45 to +1.98). At the pathway level, H□O□ treatment produced negative enrichment of “Carbon metabolism,” “Glycolysis / Gluconeogenesis,” “Citrate cycle (TCA cycle),” “Oxidative phosphorylation,” “Biosynthesis of amino acids,” “Aminoacyl-tRNA biosynthesis,” and “Ribosome,” whereas the “Two-component system” pathway showed positive enrichment (Figure 2). Together, these changes indicate coordinated regulatory activation accompanied by reduced investment in respiration, translation and biosynthesis. The canonical peroxide detoxification genes *katA* and *ahpCF* were highly expressed but were not significantly induced at the sampled time point, consistent with the high basal antioxidant capacity reported for aerobically grown *S. aureus*, particularly the constitutively high activity of KatA [52,53].

Methicillin treatment induced the strongest transcriptional response, involving upregulation of 598 genes and downregulation of 608 genes (Figure 1). Treatment provoked a focused β-lactam-induced cell wall stress response, with strong upregulation of the β-lactamase gene *blaZ* and its sensor-regulator *blaR1* (log FC = +6.33 and +2.88, respectively), together with the SCCmec-associated resistance determinant *mecA* (log FC = +2.40). The *vraDE* envelope-defence transporter genes and an adjacent *vraH*-annotated gene were also strongly upregulated (log FC = +2.42 to +3.11), reflecting a broader cell envelope adaptation profile. Consistent with these gene-level changes, the KEGG “β-Lactam resistance” and “Two-component system” pathways showed positive enrichment, whereas ribosomal genes showed broad negative enrichment (Figure 2). Methicillin additionally repressed *spa* and its positive regulator *sarS* (log□FC = −4.23 and −2.03, respectively), indicating wider remodelling of virulence-associated expression. The simultaneous induction of *blaZ*, *mecA* and envelope-defence functions therefore provides direct evidence of a β-lactam-specific cell wall stress response, consistent with the established roles of PBP2a and cell wall stress signalling in MRSA adaptation to β-lactam exposure [54].

Exposure to SNAP resulted in upregulation of 154 genes and downregulation of 123 genes (Figure 1). SNAP treatment initiated a nitrosative stress programme anchored in NO detoxification and accompanied by metabolic and respiratory adjustment (Figure 2). The NO-detoxifying flavohemoglobin *hmp* was significantly induced (log FC = +1.21), providing direct evidence that SNAP generated biologically active nitrosative stress. *hmp* encodes the principal NO-detoxification protein in *S. aureus* and is responsible for most measurable cellular NO consumption. SNAP also induced the staphyloferrin B receptor *sirA* (log□FC = +1.15), consistent with altered iron homeostasis. In contrast, the nitrate respiration-associated *narHJI* genes and regulator *nreA* were repressed (log FC = −1.91 to −2.16), as were the sulfur metabolism genes *metC* and *metE* (log□FC = −1.32 and −1.57, respectively). KEGG analysis showed positive enrichment of “Biosynthesis of cofactors”, “Pyrimidine metabolism” and “Ribosome”, together with negative enrichment of “Nitrogen metabolism”, “Cysteine and methionine metabolism”, “Biosynthesis of amino acids”, “Quorum sensing” and the aggregate “Two-component system” pathway (Figure 2). These results indicate selective metabolic and regulatory reprogramming rather than global biosynthetic arrest. Together, significant *hmp* induction and extensive remodelling of iron-, cofactor-, nitrogen-and sulfur-associated processes are consistent with the established effects of NO on respiratory electron flow, haem proteins and iron–sulfur-dependent metabolism [55,56].

Expression of the *agr* quorum sensing operon showed strong suppression with SNAP treatment (log FC = – 1.17 to –1.37 across *agrABCD*), small changes with H□O□ (log FC = –0.14 to +0.19), and moderate upregulation with methicillin (log FC = +0.53 to +1.01). These trends suggest stressor-specific modulation of quorum sensing, with nitric oxide stress in particular promoting repression of *agr*-controlled virulence pathways.

Taken together, these transcriptomic profiles demonstrate that *S. aureus* USA300 responded to diverse stressors in our experiments with tailored, robust, system-level gene expression programmes. This engagement of distinct, biologically appropriate transcriptional responses confirms that our experimental system provided a valid reference framework for relating molecular changes, in this case DNA methylation, to clinically relevant stress conditions.

### Genome-level trends in methylation

We evaluated stress-induced DNA methylation changes based on the prevalence of methylation at each base, which we termed the population level (PL), where PL is the percentage of reads in which a particular base is methylated. For example, a 60% PL indicates that the base under consideration is methylated in 60% of sequencing reads. Stress treatment effects were measured as ΔPL, the difference between pre-treatment PL and post-treatment PL. Previous research has shown that modest changes in methylation can have functional impact in bacterial systems [57,58]. Furthermore, our experimental data indicated that sublethal stressors induced ≥ 20% reductions in CFU compared to controls by the time of sampling (60 minutes), suggesting that changes of this magnitude reflect biologically meaningful perturbations. The Modkit (Oxford Nanopore Technologies) differential methylation module consistently assigned positive confidence scores (score > 0) to sites at which PL changed by ≥ 20%, whereas sites with smaller magnitude changes clustered closer to the noise threshold. Taken together, these observations support the use of |ΔPL| ≥ 20%, in combination with ≥10x coverage, as an appropriate threshold for identifying methylation changes that are both statistically reliable and biologically meaningful.

Genome-wide profiling revealed distinct pre-treatment methylation patterns for adenine (A, 6mA) and cytosine (C, 4mC and 5mC). Among sites with non-absolute (PL ≠ 0% or 100%) pre-treatment PL values, methylation levels generally clustered around 30% PL. 0.1042% of adenines, 0.0107% (4mC) and 0.0001% 5mC of cytosines had pre-treatment PL ≥90%. Conversely, 96.7049% of adenines, 98.4625% (4mC) and 99.9463% (5mC) of cytosines had PL ≤10%. Stress treatments consistently induced |ΔPL| ≥ 20% in at least 10-fold more cytosines than adenines, with < 0.0233% of cytosines and < 0.0008% of adenines showing |ΔPL| ≥ 20% with each treatment (Figure 3). A one-tailed hypergeometric test with Benjamini–Hochberg correction showed that changes in 4mC methylation were significantly enriched in gene coding regions (adjusted *p* = 0.023) compared to unannotated regions in strand-aware and/or strand-unaware analyses. Due to limited detection events, 6mA and 5mC methylation changes lacked sufficient statistical power to be tested, but we note that 100% of 6mA, and 90–100% of 5mC |ΔPL| ≥ 20% methylation changes occurred within genomic features (strand-unaware); 97.8–99.1% of |ΔPL| ≥ 20% 4mC events occurred within genomic features across the three stress treatments, also strand-unaware. A similar non-random distribution was observed for 10–20% |ΔPL| events, with 97.4–99.2% for adenine and 95.1–95.8% for cytosine occurring within genomic features (includes upstream, coding, and terminator regions, as defined in Methods ‘Genomic Feature Prediction’). These observations indicate a non-random distribution such that treatment-induced differential methylation is targeted to genomic features, consistent with the notion that DNA methylation has a functional role in *S. aureus* USA300.

**Figure 3.**
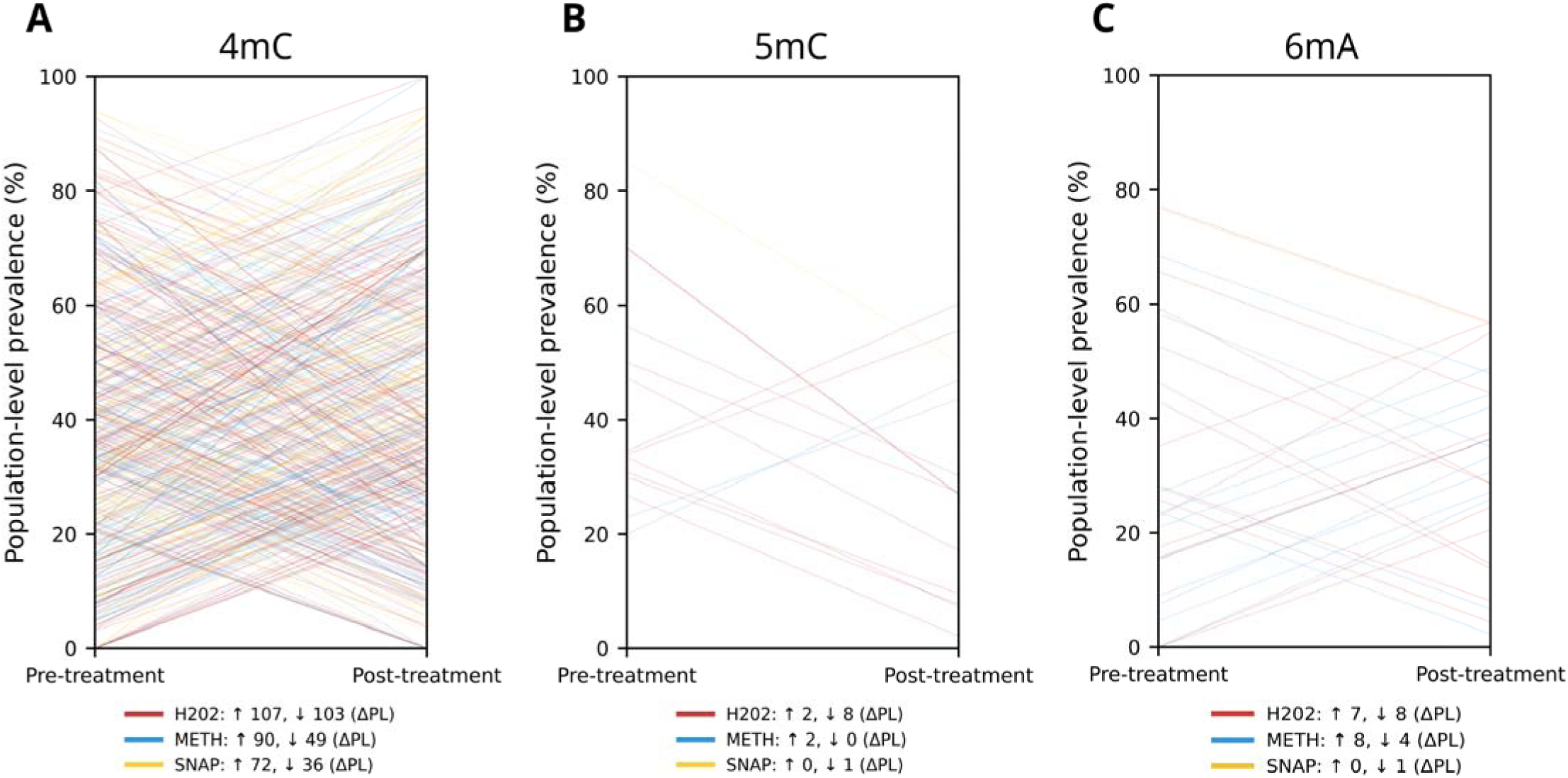
Genome-wide single base-resolution differential methylation upon stress treatment at the population level. Each line represents a base that underwent a significant change in methylation (≥ 20% ΔPL) between pre-treatment and post-treatment, for cytosine (left, 4mC; centre, 5mC) and adenine (right, 6mA). The y-axis shows population level (PL) methylation, defined as the percentage of sequencing reads with the base in methylated form; within each plot, pre-treatment PL is on the left, and post-treatment PL is on the right. Line opacity increases according to the magnitude of the methylation change (ΔPL). Line colour indicates treatment type: red (H_2_O_2_), blue (methicillin), and yellow (SNAP). The number of bases with ≥ 20% |ΔPL|, including whether methylation increased or decreased (indicating an increase or decrease in PL, respectively), is stated for each treatment. The number (n) of bases with ≥ 20% |ΔPL| is shown below each plot, grouped by treatment and direction of change: ↑ = increased methylation (higher PL), ↓ = decreased methylation (lower PL).

### Stress-responsive sites and pre-treatment methylome heterogeneity

Methylation entropy (ME), a measure of methylation pattern diversity across sequencing reads in a genomic region, was low in genomic features with mean ME = 0.146 ± 0.070 across all samples, but ME of individual genomic features spanned a wide range (0.000–0.551). As ME ranges between 0 and 1, with 0 indicating uniformity (a single methylation pattern), the mean of 0.146 ± 0.070 suggests that, in most instances, a finite number of methylation patterns coexists within genomic features. Stress treatments, moreover, induced minimal changes to ME across the genome (ΔME = 0.001 ± 0.331). Together with our knowledge of global methylation changes, such as the fact that very few |ΔPL| ≥ 20% methylation events occurred at sites with 0% PL pre-treatment (Figure 3), this supports the inference that stress treatments induce enrichment or depletion of pre-treatment methylation patterns, and that emergence of new methylation patterns is atypical. Such fine scale modulation may reflect tight, efficient regulation, and suggests that methylome heterogeneity is largely established before stress treatment [45,59].

Sites with stress-induced |ΔPL| ≥0.20 coincided with focal peaks of pre-treatment methylation entropy (ME). Among sites with coverage >10, the mean ME of 50-bp windows overlapping the response site exceeded that of supported, non-overlapping 50-bp windows within the surrounding ±500-bp region by 0.234 for 4mC (95% CI, 0.217–0.251; Holm-adjusted *p* = 1.56 × 10), 0.270 for 5mC (95% CI, 0.222–0.318; Holm-adjusted *p* = 1.86 × 10), and 0.218 for 6mA (95% CI, 0.173–0.264; Holm-adjusted *p* = 4.60 × 10). These within-location contrasts were positive at 93.2% of 4mC sites, 100% of 5mC sites, and 86.4% of 6mA sites with |ΔPL| ≥20%. Consistently, 57.3% of 4mC sites, 100% of 5mC sites, and 54.5% of 6mA sites with |ΔPL| ≥20% overlapped windows occupying the highest decile of ME within their respective local contexts.

Stress-responsive sites also showed unusually high pre-treatment ME relative to the callable genomic background. Site-overlapping ME exceeded the corresponding contig-wide mean, matched for modification and strand, by 0.262 for 4mC (95% CI, 0.246–0.278; Holm-adjusted *p* = 1.02 × 10), 0.243 for 5mC (95% CI, 0.194–0.293; Holm-adjusted *p* = 3.33 × 10), and 0.138 for 6mA (95% CI, 0.0705–0.206; Holm-adjusted *p* = 3.97 × 10). The median 4mC response site lay at the 97.8th percentile of genome-wide ME, with 67.7% of sites in the highest decile. All 5mC sites were in the genome-wide highest decile of genome-wide ME, whereas 45.5% of 6mA sites were in the highest decile. The 5mC results were therefore directionally consistent with those for 4mC and 6mA but should be interpreted cautiously because only ten response sites passed the quality filtering defined in the Methods section ‘Statistical analysis’, and the local 5mC background was uniformly low.

To extend this, we assessed whether local peaks in pre-treatment ME were predictive of |ΔPL| following treatment. After adjustment for pre-treatment PL, sequencing depth, ME support, GC content, callability and treatment, higher pre-treatment site-overlapping ME remained positively associated with greater subsequent |ΔPL| for 4mC (Holm-adjusted *p* = 0.00153; partial *R*² = 2.22%), 5mC (*p* = 0.0112; partial *R*² = 1.53%), and 6mA (*p* = 2.25 × 10 ¹³; partial *R*² = 9.40%). Together, these findings support a model in which stress-responsive methylation sites represent focal nodes of pre-treatment methylome heterogeneity that are more likely to undergo methylation change.

### Stress-induced shifts in methylome heterogeneity and transcriptional change

Changes in ME within coding and upstream regions showed potential correlation with transcriptional output (Figure 4). To mitigate statistical bias and uncover otherwise obscured trends, genomic features were grouped into percentile bins (50th, 75th, and 95th) based on the magnitude of methylation entropy change (|(ΔME)|), enabling stratified regression analysis across increasing effect sizes. For 6mA, same-strand loci showed no association at low ΔME (β ≈ 0, *p* ≫ 0.05, where β is the gradient of the log_2_FC vs |(ΔME)| trend lines in Figure 4), whereas a strong positive correlation emerged once |(ΔME)| exceeded the 75th percentile in coding regions (β ≈ 4–10, *p* < 0.001) and persisted at the 95th percentile (β ≈ 6–10, *p* < 10); upstream regions showed a similar but less significant trend. 4mC exhibited a positive association at the 75th percentile in coding regions (β ≈ 10.9–16.7, *p* < 10 ³) which was reduced at the 95th percentile; this pattern was also observed in opposing strand upstream regions (β ≈ 16.8, *p* = 2.2 × 10 ^8^), while same-strand upstream regions increased in significance by the 95th percentile (β ≈ 12.7, *p* = 3.7 × 10). 5mC displayed a similar trend to 4mC, but with weaker signal. Bootstrap 95% confidence intervals excluded zero for most significant slopes, and permutation histograms placed these β values near the extremes of their null distributions, reinforcing their non-random nature. Nevertheless, R² was often < 0.05, implying that changes in methylation entropy accounted for only a fraction of observed expression variance. Terminator region tests yielded opposing β values across percentiles, wide confidence intervals that frequently overlapped zero, and *p* ≫ 0.05 across nearly all percentile bins, limiting the interpretive value of those tests. As we defined terminators and upstream regions by windows relative to their corresponding operon, they encompassed non-regulatory intergenic sequences flanking operons; genuine signals were consequently probably diluted by noise. The weak trends are therefore presented as suggestive rather than conclusive, and more strongly suggestive in coding and upstream regions than in terminators.

**Figure 4.**
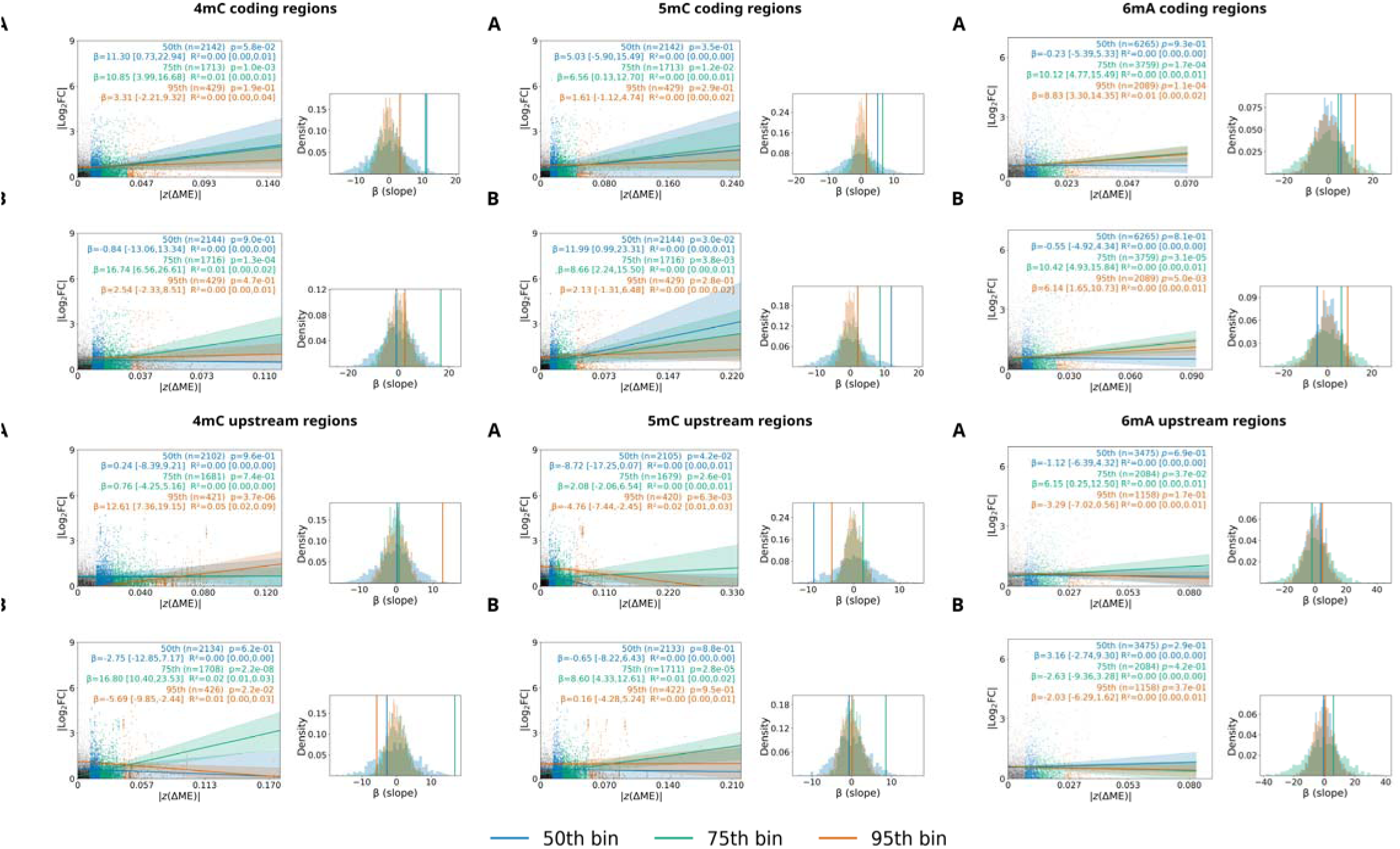
Coupling between methylation entropy change and transcriptional response in coding and upstream regions, resolved by percentile, modification type, and strand. Scatter plots show | log₂FC| as a function of | (ΔME)|. For columns 4mC, 5mC and 6mA, results are divided into (A) Same strand and (B) opposing strand coding region. Points are coloured by percentile of | (ΔME)|, with grey 0–50%, blue 50–75%, green 75–95%, orange 95–100%. An OLS fit is shown for each coloured bin, with a shaded 95% bootstrap band. Scatterplots are annotated with n (number of regions in a percentile bin), β ± 95% confidence interval (β = gradient of trendline, p = p-value, and R² = explained variance). The histogram in each panel gives bin-specific permutation nulls (1,000 shuffles of | (ΔME)| within the bin); the observed β for that bin is the vertical line of the same colour.

Collectively, the data indicate that large, strand-aware shifts in coding region methylation entropy are predictors of transcriptional change, whereas signals in upstream and terminator regions are inconclusive. In any case, while methylation entropy change appears broadly predictive, its influence on transcription is probably modulated or surpassed by other factors.

### Methylation signatures of stress response: general

Base-level resolution inspection of loci containing methylation changes in upstream and gene coding regions revealed that changes occurred mainly in genes relevant to the stress treatment, with putative methylation changes (ΔPL) corresponding qualitatively with transcriptional change (log FC) in numerous instances.

Across treatments and modification types, |ΔPL| ≥20% methylation changes affected <6.13% of protein-coding genes; for calculation of this proportion, a protein-coding gene was classified as methylation-affected when its coding sequence or assigned upstream region contained at least one |ΔPL| ≥20% site, and each affected gene contributed only once irrespective of the number of |ΔPL| ≥20% sites. The affected genes included classical stress response genes involved in detoxification and DNA repair and, more prominently, loci underpinning broader stress adaptation, including metabolic rerouting, transporter activity, immune evasion, and virulence (based on RefSeq and PGAP orthology annotations). The stressor-specific nature of these methylation profiles is consistent with targeted and adaptive epigenetic regulation. The base-level methylation changes, genomic feature overlaps, functional annotations, and associated gene expression fold changes across treatments are detailed in Supplementary Table 1. The following paragraphs highlight representative upstream and coding regions in which coincident methylation and transcription changes indicate functional relevance of DNA modification (Figure 5).

**Figure 5.**
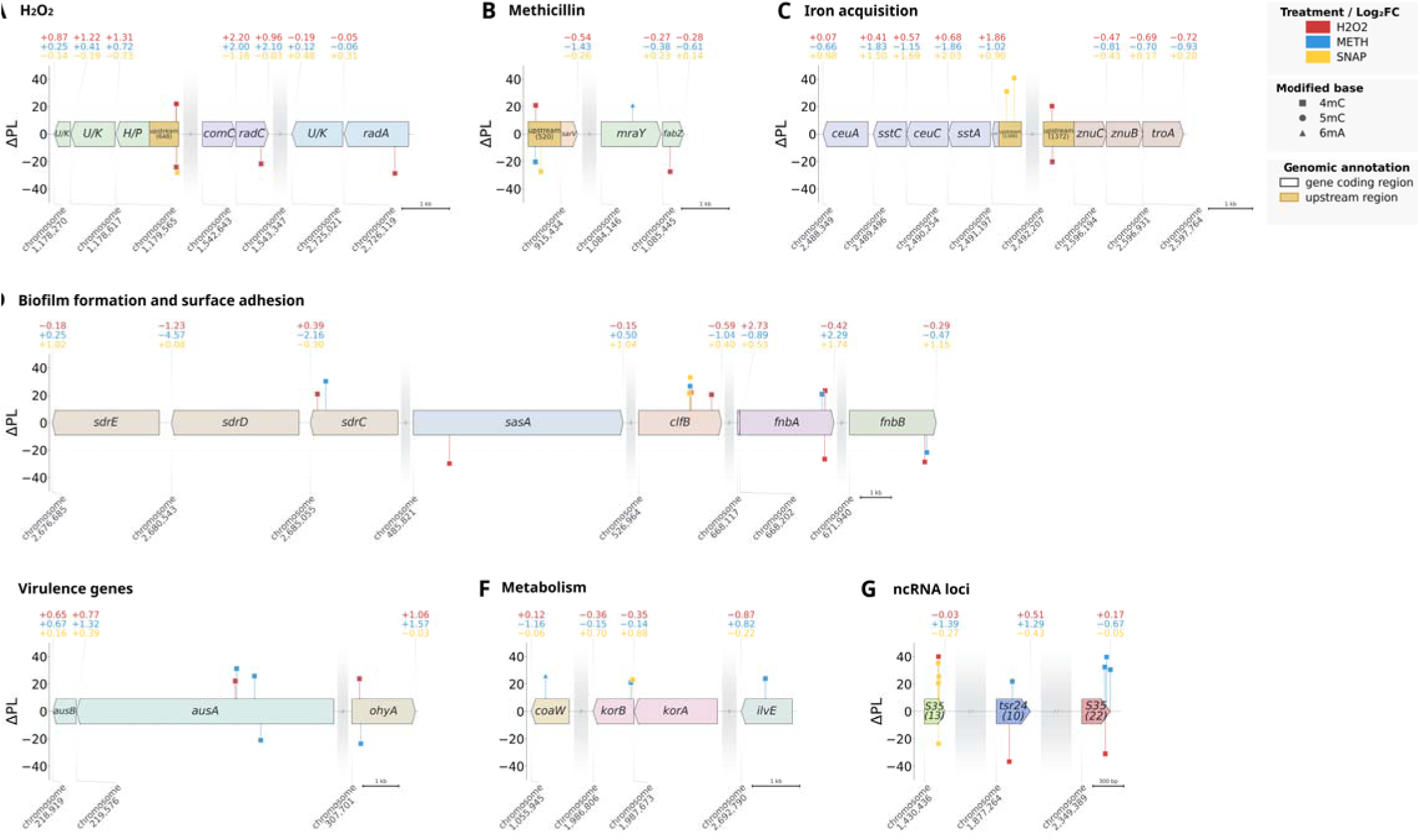
Genomic context of treatment-associated methylation changes. Genomic regions discussed in “Methylation signatures of stress response” are shown for (A) H₂O₂ response, (B) methicillin response, (C) iron acquisition, (D) biofilm formation and surface adhesion, (E) virulence, (F) metabolism, and (G) non-coding RNA loci. SNAP is excluded due to the absence of directly applicable gene hits. Genes belonging to the same operon share a colour, and up-stream regions are included when modified. Gene-specific log₂ fold-changes are shown above each feature in treatment colours and linked to the corresponding transcriptional end. Methylation changes are plotted as lollipops, with height indicating ΔPL, colour indicating treatment, and marker shape indicating 4mC, 5mC, or 6mA. Coordinate labels indicate the strand-aware transcriptional start of each feature. Grey breaks denote omitted genomic sequence; displayed regions are otherwise plotted to scale, with a scale bar provided for each panel. Hypothetical proteins are labelled H/T and unnamed proteins are labelled U/K.

### Methylation signatures of stress response: H_2_O_2_

Under H□O□ exposure, a limited number of genes directly involved in oxidative stress mitigation underwent methylation changes and transcriptional upregulation (Figure 5A). An *exonuclease domaincontaining protein* (full entry in Supplementary Table 1), for example, which may contribute to DNA repair or replication stress buffering [60], was markedly upregulated. We note the possibility that because it is encoded within a predicted operon containing a cI-like XRE-family repressor, induction of this *exonuclease domain-containing protein* may reflect prophage DNA processing following stress-induced transition from lysogeny to lytic replication, supporting phage induction in response to DNA damage [61]. This gene shared a 4mC methylation change, albeit in opposite directions, with SNAP treatment (H□O□ ΔPL = +22%, SNAP ΔPL = –28%), with contrasting transcriptional outcomes (H□O□ log□FC = +1.22, SNAP log FC = –0.19). The lack of methylation change and mild expression increase (log FC = +0.41) for this gene with methicillin treatment, therefore, alongside the opposing methylation and expression outcomes under H□O□ and SNAP treatments, suggests treatment-specific epigenetic regulation occurred at this location. Additionally, among the treatments studied here, H□O□ uniquely imposed methylation changes in coding regions of DNA repair genes *radA* (4mC, ΔPL = –29%) and *radC* (4mC, ΔPL = –22%), two genes linked to ROS survival and to host persistence *via* DNA repair [62]. These H□O□-specific methylation changes coincided with upregulation in *radC* (log FC = +0.96), which was also seen with methicillin (log□FC = +2.10), and negligible change with SNAP (log FC = +0.03), while *radA* was negligibly changed with H□O□ (log□FC = –0.05) and methicillin (log□FC = –0.06), but upregulated with SNAP (log□FC = +0.31). Collectively, although the relationship between methylation and expression was non-uniform across these contexts, H□O□ was distinctive in showing methylation changes at ROS-linked DNA repair loci.

### Methylation signatures of stress response: methicillin

In response to methicillin, multiple genes associated with cell wall remodelling and β-lactam resistance exhibited methylation changes and transcriptional repression (Figure 5B). Two such genes, isoforms of *murA* and *mraY*, within an operon involved in initiating peptidoglycan biosynthesis, exhibited upstream methylation changes exclusively under methicillin exposure (6mA, ΔPL = +21%), and displayed divergent expression across treatments (H□O□ log□FC = –0.27, methicillin –0.38, SNAP +0.23) [63]. Regulator *sarV*, which promotes autolysin expression and cell wall turnover, was consistently downregulated across treatments (H□O□ log□FC = –0.54, methicillin –1.42, SNAP –0.26), an expected adaptation for stress resilience [64]. This downregulation was paired with an upstream cytosine showing 4mC methylation change with each treatment (H□O□ ΔPL = +21%, methicillin –20%, SNAP –27%). The stronger downregulation of *sarV* with methicillin treatment is consistent with the heightened envelope stress specifically imposed by β-lactam treatment; in this context, continued autolysin activity becomes detrimental.

### Methylation signatures of stress response: SNAP

Under SNAP exposure, no methylation changes were observed in upstream or coding regions of genes directly involved in nitric oxide detoxification or survival, in contrast to the targeted stress response modifications observed with H□O□ and methicillin treatments. Beyond stress-specific modules, however, a broader layer of epigenetic modulation was observed across all treatments. These changes affected genes linked to general physiological adaptation, suggesting downstream repercussions for persistence, pathogenesis, metabolic flexibility, and stress tolerance under diverse environmental pressures.

### Methylation signatures of stress response: iron acquisition

Methylation and expression changes in iron acquisition systems were observed upon H□O□ and SNAP treatment (Figure 5C). Upstream of an operon which RefSeq highlighted as *ceuABCD*-like (containing *siderophore ABC transporter substrate-binding protein, iron chelate uptake ABC transporter family permease subunit*, and *ABC-transport proteins*), two cytosines showed 4mC methylation changes exclusive to SNAP (ΔPL = +31% and +41%). Expression change with H□O□ was small (log□FC = +0.07 to +0.68), whereas upregulation was observed with SNAP (log FC = +0.98 to +2.03), and repression with methicillin (log FC = –0.66 to –1.86). Additionally, upstream of an ABC metal uptake operon (containing *znuC*, *znuB*, and *troA-*like), two 4mC methylation change sites were identified, both unique to H□O□ treatment (ΔPL = +20% and –20%), accompanied by downregulation with H□O□ (log□FC = –0.47 to –0.72) and methicillin (log FC = –0.70 to –0.93), and mixed effects with SNAP (log FC = –0.43 to +0.20). Methylation changes may therefore contribute to regulation of metal homeostasis systems, supporting specific adaptation to oxidative and nitrosative stresses, where iron, and zinc to a lesser degree, are tightly sequestered by the host to restrict pathogen function. With SNAP, this may also help to replenish damaged haem-containing proteins. Modulation of these systems could facilitate iron and zinc scavenging and retention, supporting immune evasion and metabolic resilience during infection [65–67].

### Methylation signatures of stress response: biofilm formation and surface adhesion

Several genes associated with biofilm formation and surface adhesion exhibited methylation changes suggestive of stress-induced regulatory tuning (Figure 5D). The genes *fnbA* and *fnbB* that encode fibronectin-binding proteins, essential for surface adhesion and biofilm establishment [68], showed differential coding region 4mC methylation and expression responses. *fnbB* exhibited two methylation changes, one with H□O□ (ΔPL = –29%), and another with methicillin (ΔPL = –22%), with expression changes following a mixed pattern, including repression with H□O□ (log□FC = –0.29) and methicillin (log FC = –0.47), and strong upregulation with SNAP (log FC = +1.15). *fnbA* underwent two 4mC methylation changes in response to H□O□ (ΔPL = –26% and +23%), with methicillin showing one change (ΔPL = +21%). Corresponding expression changes were mild repression with H□O□ (log□FC = –0.42), and strong upregulation with both methicillin (log FC = +2.29) and SNAP (log FC = +1.74). The stronger expression response in *fnbA* indicates a more sensitive regulatory link to environmental stress and adaptive expression control than *fnbB*. Clumping factor *clfB*, which contributes to nasal colonisation, clumping behaviour, and immune evasion through biofilm formation [69], underwent 4mC methylation change across all conditions, with no shared sites between treatments. SNAP triggered the strongest methylation shift (ΔPL

= +22% and +33%), methicillin affected a single site (ΔPL = +26%), and H□O□ two sites (ΔPL = +21% and +22%). These shifts were paired with varying transcriptional responses (H□O□ log FC = –0.59, methicillin –1.04, SNAP +0.40). Serine-rich repeat adhesin *sasA* underwent one methylation change upon H□O□ treatment (ΔPL = –30%); this was associated with mild transcriptional repression (log FC = –0.15), whereas neither methicillin nor SNAP induced any methylation changes but both produced expression uplift of log FC = +0.50 and +1.04, respectively. *sasA* promotes surface adhesion and cell-cell aggregation, and is implicated in chronic infection settings [70]. Given these functions, stress-responsive upregulation of *sasA* may contribute to persistent colonisation or immune evasion under challenging host conditions. Similarly, the *sdrCDE* locus encodes MSCRAMM adhesins involved in aggregation, surface attachment and host interaction; SdrC specifically promotes host cell adhesion and biofilm formation through β-neurexin binding [71,72]. A single 4mC methylation change occurred in *sdrC* with methicillin treatment (ΔPL = +30%), coinciding with strong transcriptional repression (log FC = −2.16). Neither H□O□ nor SNAP produced methylation changes in this locus, with only modest *sdrC* expression changes (log FC = +0.39 and −0.30, respectively).

### Methylation signatures of stress response: virulence genes

Methylation changes in multiple virulence genes showed strong correlation with transcriptional outcomes (Figure 5E). The *non-ribosomal peptide synthetase* gene cluster *ausA* exhibited 4mC methylation changes under H□O□ and methicillin treatments, one being a shared site (H□O□ ΔPL = +22%, methicillin ΔPL =

+31%), plus three unique to methicillin (ΔPL = +31%, +26%, –21%) These changes were accompanied by greater transcriptional upregulation with H□O□ and methicillin (log FC H□O□ = +0.77, methicillin

+1.32, SNAP +0.39). Non-ribosomal peptide synthetase systems have been shown to promote phagosomal escape, particularly under oxidative stress, enhancing survival within host cells [73]. Notably, the conservation of methylation change at a specific site with both H□O□ and methicillin treatments, alongside upregulation, may reflect a pre-existing regulatory circuit. This may represent preparation for phagosomal conditions rather than a direct response to oxidative stress. *Oleate hydratase* coding region, *ohyA*, showed 4mC methylation change at a single base with both H□O□ (ΔPL = +21%) and methicillin (ΔPL = –23%), with upregulation under both treatments (H□O□ log FC = +1.06, methicillin +1.57), and downregulation with SNAP (log FC = –0.03). Oleate hydratase is vital for colonisation of mammalian skin (through the neutralisation of antimicrobial fatty acids, such as palmitoleate), but also produces hydroxy-fatty acids that suppress pro-inflammatory cytokines, thereby promoting immune evasion and facilitating infection [74,75].

### Methylation signatures of stress response: metabolism

Metabolic rerouting and respiratory adjustment emerged as consistent features of stress adaptation, with methylation changes observed in genes involved in anaerobic respiration (e.g., *2-oxoacid:ferredoxin oxidoreductase subunit beta*), redox-regulated amino acid metabolism (e.g., *branched-chain amino acid aminotransferase, ilvE*), and coenzyme A biosynthesis (e.g., *type II pantothenate kinase, coaW*; Figure 5F). Methylation change in *korB* (encoding *2-oxoacid:ferredoxin oxidoreductase subunit beta*) was exclusive to one cytosine, observed with SNAP (4mC, ΔPL = +23%) and methicillin (4mC, ΔPL = +21%), associated with transcriptional shift across treatments, particularly SNAP (H□O□ log□FC = –0.36, methicillin –0.15, SNAP +0.70). This suggests that SNAP-specific epigenetic activation promotes ferredoxin-linked redox metabolism, potentially facilitating anaerobic energy production and supplying reduced ferredoxin for downstream nitric oxide detoxification [76]. *ilvE* and *coaW* exhibited methylation changes solely under methicillin treatment (4mC, ΔPL = +23% and 6mA, ΔPL = +26%), accompanied by treatment-specific transcriptional responses: *ilvE* log FC = –0.87, +0.82, –0.22 and *coaW* log FC = +0.12, –1.16, –0.06, in both cases for H O_2_, methicillin and SNAP, respectively. This may support adaptive metabolic flexibility under methicillin stress by providing carbon skeletons and metabolic intermediates that contribute to lipid biosynthesis, energy generation, and nitrogen management, processes essential for cell envelope remodelling and β-lactam survival [77,78].

### Methylation signatures of stress response: phage-associated genes

Several phage-associated genes, including genes encoding terminase subunits, capsid proteins, a pathogenicity island protein and endonucleases, exhibited some of the largest methylation changes in the dataset, particularly with H□O□ and SNAP treatment. Shared methylation sites between these two conditions were infrequent, and they showed a mix of ΔPL directionality. Epigenetic changes coincided with strong transcriptional upregulation with H□O□, more modest upregulation and occasional repression with SNAP, and varied responses with methicillin. Oxidative stress-induced phage activation has previously been linked to virulence factor modulation and may also promote biofilm formation through sublethal prophage induction, controlled lysis, and extracellular DNA release [79,80]. These observations indicate that prophage modules can act as methylation-sensitive regulators, contributing to stress adaptation, immune modulation and biofilm maturation in hostile conditions (Figure 6).

**Figure 6.**
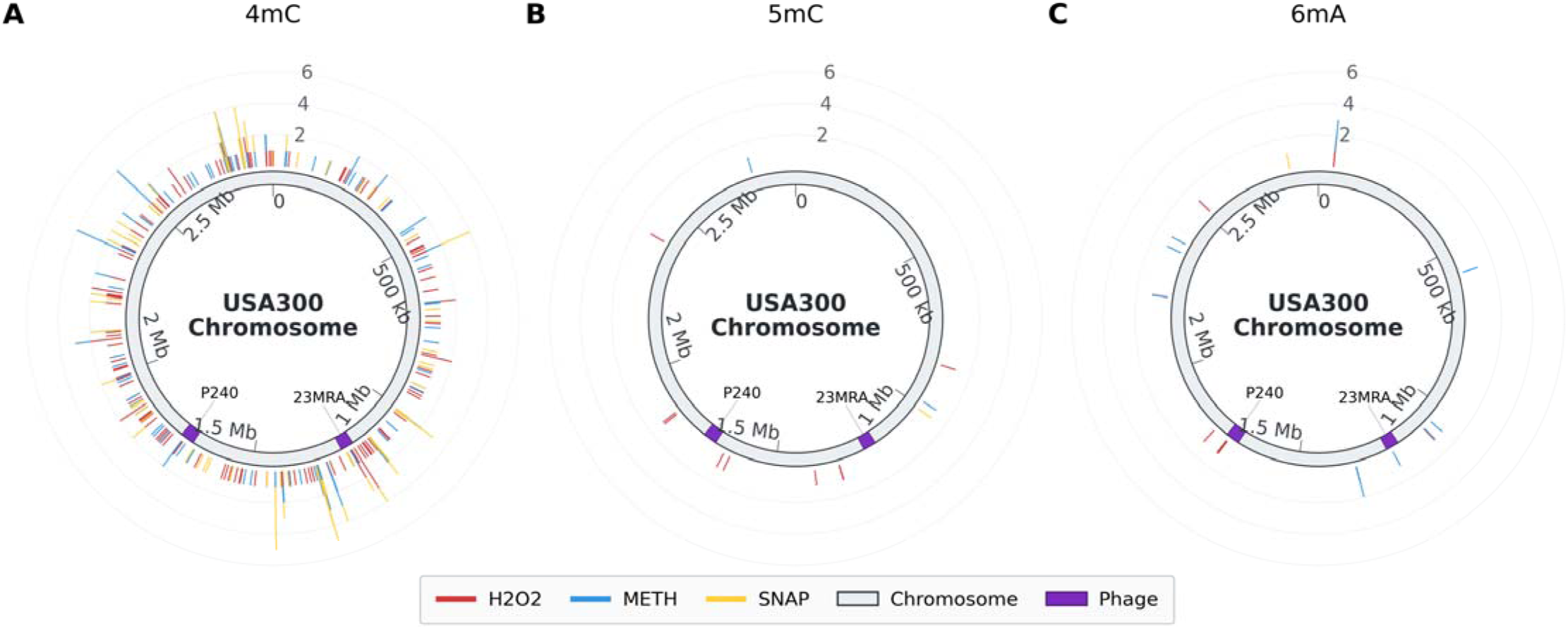
Circular maps showing the chromosomal distribution of significant (A) 4mC, (B) 5mC, and (C) 6mA changes in 100-bp bins plotted relative to phage insertions. Radial lines indicate the number of significant bases per bin and are coloured by treatment. Grey segments represent the chromosome, with prophage regions highlighted in purple and labelled internally. Genomic coordinates are shown around the inner circumference.

### Methylation of non-coding RNA loci

Genomic loci encoding non-coding RNAs (ncRNAs), including their upstream regions, also exhibited distinct and condition-specific methylation changes (Figure 5G). In contrast to methylation changes in coding and upstream regions of protein-coding genes, ncRNA methylation changes were typically confined to narrow genomic windows (∼5 bp), but the specific cytosines undergoing methylation change varied across treatments, rarely overlapping (Supplementary Table 1). Methylation shifts were predominantly increases in cytosine PL, with associated expression outcomes including neutral, downregulation and upregulation, depending on context.

S35 family members were particularly affected by stress treatments, with 12 of 30 loci representing coding regions containing at least one cytosine showing ≥ 20% |ΔPL| under at least one treatment. Members of the S35 family, plus the tsr24 family, provide illustrative examples. *S35_22* (Supplementary Table 1), for instance, underwent 4mC methylation change at a shared site in both H_2_O_2_ (ΔPL = –31%) and methicillin (ΔPL = +40%) stress conditions, along with others unique to methicillin (ΔPL +31% and +32%), coinciding with upregulation with H_2_O_2_ (log FC = +0.17) and downregulation with other treatments (methicillin log FC = –0.67, SNAP –0.05). *tsr24_10* showed two methylation changes, one unique to H_2_O_2_ (ΔPL = – 37%) and another to methicillin (ΔPL = +22%), both with upregulation (H_2_O_2_ log FC = +0.51, methicillin +1.29). Downregulation was observed with SNAP (log FC = –0.43). S35_13 showed a single 4mC methylation change with H_2_O_2_ (ΔPL = +40%), SNAP showed four 4mC change sites (ΔPL = +35%, +26%, +21%, and –23%). These coincided with downregulation with H_2_O_2_ (log FC = –0.03) and methicillin (log FC = –0.27), upregulation with SNAP (log FC = +1.39). Collectively, these examples show that opposing methylation changes within the same ncRNA often coincided with different transcriptional responses, whereas similar methylation change profiles across treatments frequently coincided with similar expression profiles. While the functions of such ncRNAs have not been well studied, some, including tsr24, have been functionally linked to virulence and stress [10,81].

A substantial proportion of methylation changes (34.1% to 48.6% of all ≥ 20% |ΔPL|) occurred within structural RNA loci, tRNA and rRNA, or related coding regions. These events were distributed across annotated tRNA and rRNA regions, with minimal overlap in the cytosines modified between treatments, suggesting targeted, stress-specific methylation. The dominant modification was increased cytosine methylation, followed by a decrease in cytosine methylation and occasional changes to adenine methylation. More detailed interpretation is hindered by the rRNA depletion used during sequencing library preparation.

## Discussion

In order to investigate the relationship between DNA methylation and transcription in *S. aureus*, we subjected USA300 to sub-lethal concentrations of three clinically relevant stressors, hydrogen peroxide (oxidative), methicillin (antibiotic), and S-nitroso-N-acetylpenicillamine (nitrosative). Below we examine the stress-induced nanopore sequencing-derived DNA methylation profiles, including whether these profiles are stressor-specific and their association with transcriptional changes in pathways including those involved in stress response, metabolism, virulence, and host interaction.

We considered stress-induced |ΔPL| ≥ 20% methylation changes to be both statistically reliable and biologically meaningful, based on our experimental observations and data analysis, and on previous reports that modest changes in methylation can have functional impact in bacteria [58,82]. Stress-induced |ΔPL| ≥ 20% methylation changes were statistically significantly enriched in genomic features: 100% of adenine methylation changes and > 97.1% of 4mC cytosine methylation changes occurred in upstream regulatory regions, coding regions, and ncRNA loci. Differential methylation events were more frequent as 4mC changes than 5mC or 6mA, suggesting that 4mC methylation makes a greater contribution to dynamic regulation in *S. aureus*. This is consistent with the finding that a persistent *S. aureus* lineage with altered *agr* regulation had significant shifts in cytosine methylation profiles, particularly in regulatory genes [18]. Cytosine methylation can act as a global epigenetic regulator, modulating transcriptional programs, virulence genes, and stress-response pathways, as demonstrated in *Helicobacter pylori* where loss of an m C MTase altered expression of more than 100 genes [83]. Moreover, cytosine methyltransferases can modulate oxidative stress survival, as shown for the mycobacterial m C MTase Rv3204, which is required for tolerance to ROS and for proper activation of antioxidant gene networks [84]. Our observations highlight cytosine methylation as a potentially important but underexplored component of the staphylococcal stress response network.

We observed potential correlations between changes in DNA methylation and gene expression across treatments, including a relationship between shifts in methylation entropy (a measure of methylation heterogeneity) and transcriptional outcomes (Figure 4). There was no apparent simple, overarching ΔPL–log_2_FCrelationship; rather, the relationship was base-specific or genomic feature-specific, with a few examples of cross-treatment consistency in the relative directions of methylation and expression changes at a particular location. Methylation and expression changes (Supplementary Table 1) occurred mainly in treatment re-sponse-specific genes including genes directly involved in stress responses, such as detoxification and DNA repair, and genes supporting broader physiological adaptations, including metabolic rerouting, transporter activity, host interaction and biofilm formation, as discussed below. One alternative explanation is that some methylation changes are secondary to transcription, occurring preferentially at loci whose DNA becomes more accessible during transcriptional remodelling. Although this explanation cannot be excluded in all cases, the overlap between methylation-affected loci (being a count of genes with ≥1 |ΔPL| ≥ 20% methylation changes within the gene coding region or associated upstream region) and significantly differentially expressed genes did not offer general support, accounting for 53.7% of H□O□-responsive genes but only 5.3% and 8.7% of methicillin-and SNAP-responsive genes, respectively.

Genome-wide methylation entropy changed only slightly across stress treatments; together with the fact that <0.01% of cytosines and adenines underwent |ΔPL| ≥ 20% methylation changes, and that <3.30% of cytosines and adenines had ΔPL >10%, this denotes retention of specific methylation patterns, indicative of tight regulation of DNA methylation. The low value of genome-wide methylation entropy, moreover, indicates that there is a finite number of methylation patterns – we conjecture that these could correspond to USA300 subpopulations present both before and after stress treatment. Importantly, sites that subsequently underwent |ΔPL| ≥20% coincided with focal peaks of pre-treatment methylation entropy relative to both their immediate genomic context and the genome-wide background. Higher pre-treatment ME at the genomic positions that changed under treatment, moreover, was correlated with larger treatment-induced |ΔPL|. Taken together, these observations support the interpretation that at least some stress-responsive sites represent nodes that may contribute disproportionately to redistribution of levels of methylation configurations from a finite repertoire of pre-treatment methylation configurations within a clonal population. Under this model, a stressor leads to preferential enrichment of methylation configurations associated with advantageous phenotypes while depleting less favourable states. This interpretation is consistent with recent evidence that methylation-mediated epigenetic phase variation can generate phenotypic heterogeneity and promote bacterial adaptation under fluctuating environmental conditions, including antibiotic exposure [14,85].

The resulting inference that DNA methylation change contributes to the characteristic phenotypic heterogeneity, adaptability and persistence of *S. aureus* [86,87] is supported by recent findings including the observation by Yamazaki *et al.* that altered cytosine methylation contributed to flexible regulation of *agr* in hospital outbreak-derived MRSA subclones that consequently acquired resistance to antibiotics and environmental stress. In contrast to the strain investigated by Yamazaki *et al.*, we did not observe methylation changes in *agr*, *pcrA* and *rpsD*, perhaps reflecting isolate-or context-specific variation in epigenetic regulation and expression [18]. Potentially in line with this interpretation, Korotetskiy *et al*. observed strain-dependent expression patterns associated with distinct methylation profiles across four antibiotic-resistant hospital isolates of *S. aureus* [19]. Together, these observations underscore the role of DNA methylation as a dynamic and selective regulatory layer that contributes to stress adaptation, possibly in a strain-and condition-specific manner.

Under hydrogen peroxide exposure, methylation changes were concentrated in genes associated with DNA repair, metal acquisition, and oxidative stress adaptation. This targeted pattern aligns with the elevated DNA damage potential of reactive oxygen species and supports the model of a methylation-linked subpopulation preconditioned for intracellular persistence, as previously proposed in oxidative stress contexts [86,89]. In contrast, methicillin treatment prompted methylation shifts in loci essential for cell envelope remodelling, including peptidoglycan biosynthesis and associated metabolic pathways. These changes likely contribute to β-lactam resilience by tuning structural maintenance and resource allocation, supporting cell wall stability under antibiotic pressure. SNAP treatment did not affect canonical nitric oxide detoxification pathways at the methylation level, but we observed condition-specific methylation in redox-regulated metabolic genes. This points to an indirect route of adaptation, potentially supporting anaerobic respiration and redox balance under nitrosative stress, in line with established roles for ferredoxin-linked pathways in nitric oxide survival [76].

Across all stress conditions, certain genes involved in virulence, colonisation and biofilm formation exhibited coincident changes in methylation and expression. This included genes that are central to immunoevasion and surface adaptation; their stress-responsive methylation patterns suggest that epigenetic modulation tunes host interaction potential in a context-specific manner [66,68]. Under oxidative stress we observed widespread methylation and expression shifts in prophage and pathogenicity island genes. This reflects a known phage response to oxidative stress, in which phage reactivation promotes biofilm formation through the release of extracellular DNA and other cellular debris [79,80]. These observations support the view that methylation changes contribute to structural and phenotypic plasticity under environmental stress, including the activation of communal defence strategies. Other virulence-associated loci such as *ausA* and *ohyA* displayed conserved methylation changes across stress treatments at shared methylation sites, and also showed shifts in transcriptional output. These patterns may reflect a stress-responsive virulence circuit epigenetically primed for host interaction, particularly under intracellular conditions such as phagosomal confinement [73,74]. Such adaptations are relevant beyond immediate stress tolerance, suggesting that methylation may act more broadly to prime virulence-associated phenotypes within clonal populations, facilitating individual and communal persistence, immune evasion, or host colonisation, depending on context.

## Limitations and future directions

While this study provides a population-level view of methylation dynamics in *S. aureus* under clinically relevant stresses, several limitations constrain mechanistic interpretation. Methylation data here rely on Oxford Nanopore calls such that DNA modifications other than methylation cannot be ruled out, although there is little evidence that other modifications, such as phosphorothioation and hypermodifications of guanine, are common features of *S. aureus* genomes. Nanopore sequencing can, moreover, distinguish methylation (5mC) from hydroxymethylation (5hmC) [49]. Future work could include further validation of methylation changes using orthogonal approaches such as LC-MS/MS of digested DNA.

The use of bulk sequencing precluded resolution of co-occurring methylation and transcriptional events at the single cell level, potentially masking rare subpopulation behaviours and preventing definitive identification of distinct epigenetic subpopulations and their phenotypic outcomes. Similarly, our bulk transcriptomic data reflected population-averaged expression, limiting detection of less abundant transcriptional states. Single cell and/or subpopulation-specific methylome and transcriptome profiling in an *in vitro* context and in *in vivo* infection models is desirable to link epigenetic variation directly to phenotypic heterogeneity. The high levels of methylation change observed in tRNA and rRNA loci suggest roles in regulation of translation which merit investigation.

## Conclusions

Our findings suggest that *Staphylococcus aureus* USA300 undergoes targeted, base-specific DNA methylation changes in response to clinically relevant stressors. No obvious overarching trend was observed linking the relative directions of changes in methylation and transcription, suggesting transcription regulation *via* methylation is base-and context-specific. Bases undergoing significant changes in methylation level appear to be involved in differentiation of stable epigenetic patterns which mostly existed pre-treatment in the population, consistent with epigenetic heterogeneity as a pre-configured state permitting rapid adaptation to environmental challenge. This supports a model in which a stress-naive population is already heterogeneously primed for environmental challenge, with stress exposure selectively fine tuning a mostly pre-treatment repertoire of DNA methylome configurations. DNA methylation may thereby enable dynamic subpopulation partitioning, adjusting and reinforcing transcriptional profiles suited to both stress tolerance and host interaction. Affected loci here included both classical stress response genes and, more prominently, genes underpinning persistence, immune evasion, metabolic rerouting, and biofilm formation - traits central to chronic infection and treatment resilience. These observations point toward an epigenetic architecture that favours condition-specific adaptation but is also geared toward pathogenesis and host survival niches.

## Data availability

All sequence data generated are available on the Sequence Read Archive (NCBI SRA), associated with the genomic assembly under BioProject ID: PRJNA1228599.

## Author Contributions Statement

The project was conceptualised and supervised by Stefan Bagby and Maisem Laabei. Funding was acquired by Stefan Bagby. Planning of experimental work was conducted by Luke Jones and Stefan Bagby. Experimental work and data analysis was conducted by Luke Jones. Writing (original draft) was done by Luke Jones. Writing (review & editing) was done by Stefan Bagby, Luke Jones and Maisem Laabei.

