## Supplemental Table 1 for "Stress-induced DNA methylome plasticity and transcriptional re-programming in *Staphylococcus aureus*"

<table>
<tr><th>region</th><th>function</th><th>coordinates</th><th>H2O2_4mC</th><th>H2O2_5mC</th><th>H2O2_6mA</th><th>METH_4mC</th><th>METH_5mC</th><th>METH_6mA</th><th>SNAP_4mC</th><th>SNAP_5mC</th><th>SNAP_6mA</th><th>H2O2-Log2FC</th><th>METH-Log2FC</th><th>SNAP-Log2FC</th></tr>
<tr><td>gene:S35_1</td><td>S35_1</td><td>chromosome:35307-35537(+)</td><td></td><td></td><td></td><td>chromosome:35409-35410(+):+0.303571[0.525,0.82857144]</td><td></td><td></td><td>chromosome:35464-35465(+):+0.216247[0.1521739,0.36842105]</td><td></td><td></td><td></td><td></td><td></td></tr>
<tr><td>gene:S35_10</td><td>S35_10</td><td>chromosome:1272272-1272605(-)</td><td>|||chromosome:1272586-1272587(+):+0.28499[0.3529412,0.63793105]</td><td></td><td></td><td>chromosome:1272541-1272542(-):+0.224868[0.071428575,0.2962963]|||chromosome:1272430-1272431(+):+0.223529[0.6,0.8235294]</td><td></td><td></td><td></td><td></td><td></td><td>0.199975</td><td>-0.125117</td><td>-0.281566</td></tr>
<tr><td>gene:S35_11</td><td>S35_11</td><td>chromosome:1273732-1274054(-)</td><td>chromosome:1273883-1273884(-):+0.373016[0.071428575,0.44444445]</td><td></td><td></td><td>chromosome:1273882-1273883(-):+0.204545[0.54545456,0.75];chromosome:1273883-1273884(-):-0.270588[0.47058824,0.2]</td><td></td><td></td><td>chromosome:1273882-1273883(-):-0.236264[0.8076923,0.5714286]</td><td></td><td></td><td>0.502325</td><td>0.237809</td><td>-0.620971</td></tr>
<tr><td>gene:S35_12</td><td>S35_12</td><td>chromosome:1430093-1430257(+)</td><td></td><td></td><td></td><td>|||chromosome:1430157-1430158(-):+0.266667[0.6,0.8666667]</td><td></td><td></td><td></td><td></td><td></td><td>-0.138484</td><td>1.61876</td><td>-0.217279</td></tr>
<tr><td>gene:S35_13</td><td>S35_13</td><td>chromosome:1430435-1430636(+)</td><td>|||chromosome:1430572-1430573(-):+0.4[0.3,0.7]</td><td></td><td></td><td></td><td></td><td></td><td>chromosome:1430569-1430570(+):+0.352564[0.23076923,0.5833333];chromosome:1430577-1430578(+):+0.254945[0.51428574,0.7692308]|||chromosome:1430571-1430572(-):+0.206767[0.57894737,0.78571427];chromosome:1430573-1430574(-):-0.234756[0.6097561,0.375]</td><td></td><td></td><td>-0.0278807</td><td>1.38932</td><td>-0.274363</td></tr>
</table>

| | | | | | | | | | | | | | | |
|---|---|---|---|---|---|---|---|---|---|---|---|---|---|---|
| gene:S35_16 | S35_16 | chromosome:1834048-1834374(-) | \|\|\|chromosome:1834204-1834205(+):+0.315942[0.2173913,0.53333336] | | | | | | \|\|\|chromosome:1834206-1834207(+):-0.204503[0.5121951,0.30769232] | | | -0.0583047 | -0.8521 | 0.438552 |
| gene:S35_18 | S35_18 | chromosome:1923360-1923689(+) | chromosome:1923543-1923544(+):-0.278571[0.31428573,0.035714287] | | | chromosome:1923421-1923422(+):+0.237879[0.05,0.28787878] | | | | | | 0.248459 | -0.133303 | -0.119825 |
| gene:S35_22 | S35_22 | chromosome:2349388-2349667(+) | chromosome:2349622-2349623(+):-0.309524[0.64285713,0.33333334] | | | chromosome:2349622-2349623(+):+0.397059[0.25,0.64705884];chromosome:2349665-2349666(+):+0.305556[0.44444445,0.75]\|\|\|chromosome:2349611-2349612(-):+0.323077[0.07692308,0.4] | | | | | | 0.170038 | -0.670593 | -0.049468 |
| gene:S35_26 | S35_26 | chromosome:2460952-2461138(-) | \|\|\|chromosome:2461007-2461008(+):-0.319672[0.5,0.18032786] | | | | | | | | | 0.662128 | -0.0160274 | -0.432267 |
| gene:S35_27 | S35_27 | chromosome:2627831-2628052(+) | | | | chromosome:2627951-2627952(+):+0.331731[0.4375,0.7692308] | | | | | | 0.219565 | 0.557856 | -0.307076 |
| gene:S35_3 | S35_3 | chromosome:260277-260547(+) | \|\|\|chromosome:260291-260292(-):+0.212202[0.115384616,0.3275862] | | | \|\|\|chromosome:260390-260391(-):+0.200717[0.3548387,0.5555556] | | | | | | -0.00809045 | -1.05998 | -0.197858 |
| gene:S35_9 | S35_9 | chromosome:1262867-1263190(-) | \|\|\|chromosome:1263081-1263082(+):-0.251196[0.84210527,0.59090906] | | | | | | | | | 0.302895 | -0.151921 | -0.555915 |
| gene:rli23_1 | rli23_1 | chromosome:362604-362703(+) | | | | chromosome:362697-362698(+):-0.268251[0.7419355,0.47368422] | | | | | | 1.6379 | -1.14169 | -0.837343 |
| gene:tsr24_10 | tsr24_10 | chromosome:1877263-1877597(+) | \|\|\|chromosome:1877389-1877390(-):-0.366071[0.9285714,0.5625] | | | chromosome:1877418-1877419(+):+0.219231[0.05,0.26923078] | | | | | | 0.511949 | 1.28938 | -0.432236 |

| | | | | | | | | | | | | | | |
|---|---|---|---|---|---|---|---|---|---|---|---|---|---|---|
| gene:tsr24_11 | tsr24_11 | chromosome:2098787-2099021(-) | chromosome:2098834-2098835(-):-0.396104[0.71428573,0.3181818]\|\|\|chromosome:2098826-2098827(+):-0.217949[0.8333333,0.61538464] | | | chromosome:2098878-2098879(-):+0.20597[0.6,0.80597013] | | | | | | 0.112096 | -0.664311 | 0.286161 |
| gene:tsr24_2 | tsr24_2 | chromosome:825508-825861(+) | chromosome:825742-825743(+):-0.208333[0.45833334,0.25];chromosome:825749-825750(+):+0.201754[0.6315789,0.8333333] | | | | | | | | | 0.863627 | 1.67356 | -0.367814 |
| gene:tsr24_6 | tsr24_6 | chromosome:1393224-1393446(+) | | | | | | | \|\|\|chromosome:1393378-1393379(-):-0.256667[0.41666666,0.16] | | | 1.4842 | -0.792498 | -0.8198 |
| gene:tsr24_7 | tsr24_7 | chromosome:1410738-1410977(-) | chromosome:1410835-1410836(-):-0.209091[0.8,0.59090906] | | | \|\|\|chromosome:1410830-1410831(+):+0.249417[0.20512821,0.45454547] | | | \|\|\|chromosome:1410808-1410809(+):-0.212121[0.3939394,0.18181819] | | | -0.103587 | 0.905643 | -0.880702 |
| gene:tsr24_8 | tsr24_8 | chromosome:1499164-1499403(-) | | | | \|\|\|chromosome:1499241-1499242(+):+0.309955[0.38235295,0.6923077] | | | | | | 1.28996 | 0.529559 | 0.495681 |
| gene:tsr24_9 | tsr24_9 | chromosome:1860449-1860688(-) | \|\|\|chromosome:1860519-1860520(+):-0.22043[0.38709676,0.16666667] | | | | | | | | | -0.0743698 | 1.05634 | -0.50493 |
| gene;ACNSPI_00145 | aminoglycoside O-phosphotransferase APH(3')-IIIa | pUSA300-1:22523-23318(-) | \|\|\|pUSA300-1:22744-22745(+):+0.233333[0.3,0.53333336] | | | pUSA300-1:23030-23031(-):+0.269231[0.1923077,0.46153846] | | | | | | 0.0199552 | 0.5428 | 0.0504578 |
| gene;ACNSPI_00180 | alkyl hydroperoxide reductase subunit F | chromosome:1070-2594(+) | | | | | | | chromosome:2136-2137(+):-0.27451[0.33333334,0.05882353] | | | -0.24055 | -1.42397 | 0.524968 |

<table>
<tr><th></th><th></th><th></th><th></th><th></th><th></th><th></th><th></th><th></th><th></th><th></th><th></th><th></th><th></th><th></th></tr>
<tr><td>gene;ACNSPI_00375</td><td>hypothetical protein</td><td>chromosome:35334-35631(-)</td><td></td><td></td><td></td><td>|||chromosome:35409-35410(+):+0.303571[0.525,0.82857144]</td><td></td><td></td><td>|||chromosome:35464-35465(+):+0.216247[0.1521739,0.36842105]</td><td></td><td></td><td>0.64641</td><td>-0.948721</td><td>-1.66085</td></tr>
<tr><td>gene;ACNSPI_00390</td><td>LLM class flavin-dependent oxidoreductase</td><td>chromosome:37363-38425(-)</td><td></td><td></td><td></td><td>chromosome:37847-37848(-):+0.302083[0.16666667,0.46875]</td><td></td><td></td><td></td><td></td><td></td><td>-0.523412</td><td>-0.0336714</td><td>0.45681</td></tr>
<tr><td>gene;ACNSPI_00420</td><td>transcription antiterminator</td><td>chromosome:43750-45706(+)</td><td>|||chromosome:45595-45596(-):-0.22[0.3,0.08]</td><td></td><td>chromosome:45603-45604(+):-0.29064[0.42857143,0.13793103]</td><td></td><td></td><td>|||chromosome:45654-45655(-):+0.21657[0.08955224,0.30612245]</td><td></td><td></td><td></td><td>-0.150827</td><td>0.393641</td><td>0.9073</td></tr>
<tr><td>gene;ACNSPI_00425</td><td>PTS sugar transporter subunit IIA</td><td>chromosome:45710-46154(+)</td><td></td><td></td><td>chromosome:46139-46140(+):-0.212698[0.6571429,0.44444445];chromosome:46146-46147(+):-0.240602[0.5263158,0.2857143]</td><td></td><td></td><td></td><td></td><td></td><td></td><td>0.532869</td><td>1.07124</td><td>1.05881</td></tr>
<tr><td>gene;ACNSPI_00430</td><td>PTS sugar transporter subunit IIB</td><td>chromosome:46155-46440(+)</td><td></td><td></td><td></td><td></td><td></td><td>chromosome:46187-46188(+):+0.20979[0.15384616,0.36363637]</td><td></td><td></td><td></td><td>0.972412</td><td>1.14763</td><td>0.814662</td></tr>
<tr><td>gene;ACNSPI_00535</td><td>PTS sugar transporter subunit IIC</td><td>chromosome:68960-69950(-)</td><td></td><td></td><td></td><td></td><td></td><td></td><td>|||chromosome:69775-69776(+):-0.218254[0.8055556,0.5873016]</td><td></td><td></td><td>-0.564456</td><td>-0.798991</td><td>-0.275979</td></tr>
<tr><td>gene;ACNSPI_00805</td><td>6-phospho-beta-glucosidase</td><td>chromosome:117413-118850(-)</td><td></td><td></td><td></td><td>chromosome:118566-118567(-):+0.252174[0.4,0.65217394]</td><td></td><td></td><td></td><td></td><td></td><td>0.0103718</td><td>0.439825</td><td>0.0417711</td></tr>
<tr><td>gene;ACNSPI_00990</td><td>3-hydroxyacyl-CoA dehydrogenase/enoyl-CoA hydratase family protein</td><td>chromosome:158155-160417(+)</td><td></td><td></td><td></td><td></td><td></td><td></td><td>|||chromosome:159266-159267(-):+0.288462[0.25,0.53846157]</td><td></td><td></td><td>0.615895</td><td>0.0560969</td><td>-0.556366</td></tr>
<tr><td>gene;ACNSPI_00995</td><td>acetyl-CoA C-acyltransferase</td><td>chromosome:160446-161631(+)</td><td></td><td></td><td></td><td>chromosome:161591-161592(+):+0.324357[0.21052632,0.53488374]</td><td></td><td></td><td></td><td></td><td></td><td>0.0206173</td><td>-0.260428</td><td>-0.0977758</td></tr>
<tr><td>gene;ACNSPI_01160</td><td>PTS transporter subunit EIIC</td><td>chromosome:203084-204539(-)</td><td>chromosome:203915-203916(-):-0.200669[0.46153846,0.26086956]</td><td></td><td></td><td></td><td></td><td></td><td></td><td></td><td></td><td>-0.0753777</td><td>-0.592446</td><td>0.296003</td></tr>
</table>

| | | | | | | | | | | | | | | |
|---|---|---|---|---|---|---|---|---|---|---|---|---|---|---|
| gene;ACNSPI_01175 | glucose-specific PTS transporter subunit IIBC | chromosome:207087-209133(+) | \|\|\|chromosome:207513-207514(-):-0.246956[0.5869565,0.34] | | | | | | | | | -0.627382 | -2.48642 | 0.710647 |
| gene;ACNSPI_01220 | YagU family protein | chromosome:218111-218591(+) | chromosome:218495-218496(+):-0.210526[0.21052632,0] | | | | | | | | | -0.469805 | 0.853675 | -0.586348 |
| gene;ACNSPI_01230 | non-ribosomal peptide synthetase | chromosome:219575-226751(-) | chromosome:224018-224019(-):+0.221429[0.17857143,0.4] | | | chromosome:224018-224019(-):+0.3125[0.25,0.5625]\|\|\|chromosome:224535-224536(+):+0.257143[0.1,0.35714287];chromosome:224708-224709(+):-0.210237[0.34210527,0.13186814] | | | | | | 0.772194 | 1.32236 | 0.390159 |
| gene;ACNSPI_01280 | cation diffusion facilitator family transporter | chromosome:235435-236395(-) | | | | chromosome:235807-235808(-):+0.201243[0.08080808,0.2820513] | | | | | | 0.168162 | -0.626488 | -0.0623164 |
| gene;ACNSPI_01415 | phosphate/phosphite/phosphonate ABC transporter substrate-binding protein | chromosome:265685-266642(+) | \|\|\|chromosome:265972-265973(-):-0.245645[0.31707317,0.071428575] | | | | | | | | | 1.29349 | -0.497782 | -0.088464 |
| gene;ACNSPI_01425 | phosphonate ABC transporter%2C permease protein PhnE | chromosome:267630-268431(+) | | | | | | | chromosome:268286-268287(+):-0.26186[0.44186047,0.18] | | | 1.60104 | 0.493955 | -0.53983 |
| gene;ACNSPI_01475 | oligosaccharide flippase family protein | chromosome:276531-277962(-) | chromosome:277851-277852(-):+0.318681[0.14285715,0.46153846] | | | | | | | | | 0.278343 | -0.707559 | -0.0576255 |
| gene;ACNSPI_01590 | L-lactate permease | chromosome:302627-304220(-) | | | | | | | chromosome:303720-303721(-):-0.25[0.55,0.3] | | | 0.219422 | -0.0746335 | -0.6902 |
| gene;ACNSPI_01610 | oleate hydratase | chromosome:307700-309476(+) | chromosome:307931-307932(+):+0.238095[0.42857143,0.6666667] | | | chromosome:307931-307932(+):-0.235556[0.68,0.44444445] | | | | | | 1.06271 | 1.56722 | -0.0269741 |
| gene;ACNSPI_01805 | ABC transporter permease | chromosome:348381-349338(-) | | | | | | | chromosome:349048-349049(-):+0.43089[0.41121495,0.84210527] | | | 1.04063 | -0.120899 | -0.0447478 |

| | | | | | | | | | | | | | | |
|---|---|---|---|---|---|---|---|---|---|---|---|---|---|---|
| gene;ACNSPI_01850 | arginine-ornithine antiporter | chromosome:358260-359682(+) | chromosome:359221-359222(+):-0.204872[0.49333334,0.28846154] | | | | | | | | | 0.348404 | -0.53576 | -0.665826 |
| gene;ACNSPI_01865 | carbamate kinase | chromosome:361468-362398(+) | | | | chromosome:361936-361937(+):+0.247412[0.071428575,0.3188406] | | | | | | 0.172072 | 0.167653 | 0.0533807 |
| gene;ACNSPI_01875 | IS3 family transposase | chromosome:362497-363498(-) | | | | \|\|\|chromosome:362697-362698(+):-0.268251[0.7419355,0.47368422] | | | | | | | | |
| gene;ACNSPI_02305 | anion permease | chromosome:449846-451265(-) | | | | \|\|\|chromosome:450064-450065(+):-0.268092[0.53125,0.2631579];chromosome:450070-450071(+):+0.253232[0.1521739,0.4054054] | | | | | | -0.283639 | -1.72314 | -0.488766 |
| gene;ACNSPI_02345 | S-adenosyl-l-methionine hydroxide adenosyltransferase family protein | chromosome:457352-458201(+) | | | | chromosome:457566-457567(+):-0.204678[0.31578946,0.11111111] | | | | | | -1.93963 | 1.56096 | -1.19067 |
| gene;ACNSPI_02435 | intracellular adhesion protein IcaD | chromosome:475884-476190(-) | | | | | | | \|\|\|chromosome:476169-476170(+):-0.308271[0.7368421,0.42857143] | | | -0.508855 | 0.387072 | 0.152437 |
| gene;ACNSPI_02440 | poly-beta-1%2C6 N-acetyl-D-glucosamine synthase IcaA | chromosome:476153-477392(-) | | | | | | | \|\|\|chromosome:476169-476170(+):-0.308271[0.7368421,0.42857143] | | | 0.89835 | -0.195611 | 0.486224 |
| gene;ACNSPI_02500 | serine-rich repeat glycoprotein adhesin SasA | chromosome:485820-492636(+) | chromosome:486991-486992(+):-0.297203[0.61538464,0.3181818] | | | | | | | | | -0.151388 | 0.499412 | 1.03656 |
| gene;ACNSPI_02555 | YhgE/Pip family protein | chromosome:508506-511488(+) | chromosome:509629-509630(+):+0.254386[0.078947365,0.33333334] | | | | | | | | | 0.270579 | -0.271865 | -0.0659837 |

<table>
<tr><th></th><th></th><th></th><th></th><th></th><th></th><th></th><th></th><th></th><th></th><th></th><th></th><th></th><th></th><th></th></tr>
<tr><td>gene;ACNSPI_02565</td><td>fructose-specific PTS transporter subunit EIIC</td><td>chromosome:512551-514504(-)</td><td>chromosome:513224-513225(-):-0.202381[0.2857143,0.083333336]</td><td></td><td></td><td></td><td></td><td></td><td></td><td></td><td></td><td>-0.389623</td><td>-0.417737</td><td>0.431894</td></tr>
<tr><td>gene;ACNSPI_02585</td><td>zinc metalloproteinase aureolysin</td><td>chromosome:518361-519891(+)</td><td>|||chromosome:519561-519562(-):+0.216667[0.33333334,0.55]</td><td></td><td></td><td></td><td></td><td></td><td></td><td></td><td></td><td>0.358435</td><td>-2.22223</td><td>-1.09297</td></tr>
<tr><td>gene;ACNSPI_02625</td><td>MSCRAMM family adhesin clumping factor ClfB</td><td>chromosome:526963-529663(+)</td><td>chromosome:528665-528666(+):+0.221839[0.43333334,0.6551724]|||chromosome:529332-529333(-):+0.205128[0.7948718,1]</td><td></td><td></td><td>|||chromosome:528637-528638(-):+0.266667[0.33333334,0.6]</td><td></td><td></td><td>|||chromosome:528615-528616(-):+0.215311[0.42105263,0.6363636];chromosome:528643-528644(-):+0.331933[0.38235295,0.71428573]</td><td></td><td></td><td>-0.59428</td><td>-1.04366</td><td>0.396314</td></tr>
<tr><td>gene;ACNSPI_02700</td><td>NAD(P)-binding protein</td><td>chromosome:542052-542658(+)</td><td></td><td></td><td></td><td>chromosome:542114-542115(+):+0.228889[0.16,0.3888889]</td><td></td><td></td><td></td><td></td><td></td><td>0.210059</td><td>1.8791</td><td>0.184641</td></tr>
<tr><td>gene;ACNSPI_02820</td><td>oxidoreductase</td><td>chromosome:567352-568213(-)</td><td></td><td></td><td></td><td>chromosome:568187-568188(-):+0.202797[0.18181819,0.3846154]</td><td></td><td></td><td></td><td></td><td></td><td>-0.193633</td><td>0.352174</td><td>-0.556543</td></tr>
<tr><td>gene;ACNSPI_02875</td><td>fructosamine kinase family protein</td><td>chromosome:576367-577234(+)</td><td>|||chromosome:576647-576648(-):+0.301314[0.046511628,0.3478261]</td><td></td><td></td><td></td><td></td><td>chromosome:576580-576581(+):-0.212403[0.27906978,0.06666667]</td><td></td><td></td><td></td><td>0.158709</td><td>0.269303</td><td>-0.13211</td></tr>
<tr><td>gene;ACNSPI_03015</td><td>aminotransferase class I/II-fold pyridoxal phosphate-dependent enzyme</td><td>chromosome:602290-603445(+)</td><td>|||chromosome:602404-602405(-):+0.315556[0.24,0.5555556]</td><td></td><td></td><td></td><td></td><td></td><td></td><td></td><td></td><td>-0.382514</td><td>0.839202</td><td>-0.674224</td></tr>
<tr><td>gene;ACNSPI_03185</td><td>L-serine ammonia-lyase%2C iron-sulfur-dependent%2C subunit alpha</td><td>chromosome:637697-638597(+)</td><td>chromosome:638233-638234(+):-0.300926[0.7083333,0.4074074]</td><td></td><td></td><td></td><td></td><td></td><td></td><td></td><td></td><td>0.000363998</td><td>-0.821197</td><td>-0.110736</td></tr>
<tr><td>gene;ACNSPI_03345</td><td>fibronectin-binding protein FnbA</td><td>chromosome:668201-671258(+)</td><td>|||chromosome:670955-670956(-):-0.264444[0.32,0.055555556];chromosome:670969-670970(-):+0.233333[0.5,0.73333335]</td><td></td><td></td><td>|||chromosome:670870-670871(-):+0.20903[0.26923078,0.47826087]</td><td></td><td></td><td></td><td></td><td></td><td>-0.415707</td><td>2.28516</td><td>1.73926</td></tr>
</table>

| | | | | | | | | | | | | | | |
|---|---|---|---|---|---|---|---|---|---|---|---|---|---|---|
| gene;ACNSPI_03355 | fibronectin-binding protein FnbB | chromosome:671939-674762(+) | chromosome:674388-674389(+):-0.285362[0.390625,0.10526316] | | | |||chromosome:674453-674454(-):-0.216783[0.4090909,0.1923077] | | | | | | -0.288636 | -0.467591 | 1.14657 |
| gene;ACNSPI_03430 | hypothetical protein | chromosome:691149-692474(+) | | | | | | | |||chromosome:691362-691363(-):-0.27381[0.41666666,0.14285715] | | | | | |
| gene;ACNSPI_03540 | ABC transporter ATP-binding protein | chromosome:710866-711616(+) | |||chromosome:711222-711223(-):-0.22686[0.37579617,0.14893617] | | | | | | | | | 1.02099 | -0.692138 | 0.0980786 |
| gene;ACNSPI_03615 | hypothetical protein | chromosome:726425-726545(+) | | | | | | | chromosome:726430-726431(+):+0.246753[0.18181819,0.42857143] | | | 0.751325 | 0.932991 | -0.354686 |
| gene;ACNSPI_03640 | MFS transporter | chromosome:730676-732077(+) | chromosome:731679-731680(+):+0.241597[0.29411766,0.53571427];chromosome:731856-731857(+):+0.214973[0.09090909,0.30588236] | | | | | | | | | -0.575974 | 0.776895 | -0.529676 |
| gene;ACNSPI_03680 | C39 family peptidase | chromosome:742979-743636(-) | | | | chromosome:743481-743482(-):-0.216346[0.5625,0.34615386] | | | | | | -0.547985 | -0.393463 | 0.742507 |
| gene;ACNSPI_03900 | nitrate reductase subunit alpha | chromosome:783519-787209(+) | | | | chromosome:785195-785196(+):+0.485714[0.21428572,0.7] | | | | | | -0.452456 | 0.514277 | -1.60674 |
| gene;ACNSPI_04020 | NAD(P)/FAD-dependent oxidoreductase | chromosome:810176-811211(+) | | | | | | | chromosome:810210-810211(+):+0.209549[0.4827586,0.6923077] | | | -0.255202 | -0.0718402 | 0.858998 |
| gene;ACNSPI_04055 | malate dehydrogenase (quinone) | chromosome:818468-819947(+) | chromosome:819357-819358(+):-0.210084[0.3529412,0.14285715] | | | | | | | | | -0.411586 | -0.83311 | -0.231215 |

| | | | | | | | | | | | | | | |
|---|---|---|---|---|---|---|---|---|---|---|---|---|---|---|
| gene;ACNSPI_04210 | ribose 5-phosphate isomerase A | chromosome:847656-848343(+) | | | | | | | chromosome:847965-847966(+):-0.219048[0.26666668,0.04761905] | | | -0.20824 | -0.994181 | 0.181885 |
| gene;ACNSPI_04285 | bile acid:sodium symporter family protein | chromosome:864909-865827(+) | chromosome:865227-865228(+):+0.212821[0.15384616,0.36666667] | | | | | | | | | -0.691914 | 2.63665 | -0.99713 |
| gene;ACNSPI_04320 | hypothetical protein | chromosome:870735-871065(-) | | | | \|\|\|chromosome:870826-870827(+):-0.329114[0.32911393,0] | | | | | | 0.531642 | 0.757656 | -0.669755 |
| gene;ACNSPI_04370 | FAD-dependent monooxygenase | chromosome:881050-882175(+) | \|\|\|chromosome:881920-881921(-):-0.266129[0.75,0.48387095] | | | | | | | | | -0.366545 | -0.916376 | 0.183616 |
| gene;ACNSPI_04620 | GRP family sugar transporter | chromosome:926009-926873(-) | | | | \|\|\|chromosome:926788-926789(+):+0.224532[0.15384616,0.3783784] | | | | | | -0.311534 | -1.28413 | 0.965745 |
| gene;ACNSPI_04635 | DNA topoisomerase III | chromosome:928127-930263(+) | chromosome:928697-928698(+):+0.260526[0.039473683,0.3] | | | | | | | | | -0.446791 | -0.486848 | -0.0548332 |
| gene;ACNSPI_04705 | 50S ribosomal protein L14 | chromosome:937795-938164(+) | chromosome:938072-938073(+):-0.247863[0.6923077,0.44444445] | | | | | | | | | -0.619905 | -1.48583 | 0.2635 |
| gene;ACNSPI_04790 | 50S ribosomal protein L17 | chromosome:946194-946563(+) | chromosome:946523-946524(+):+0.262599[0.46153846,0.7241379] | | | | | | | | | -0.340682 | -1.29911 | 0.082106 |
| gene;ACNSPI_04980 | BCCT family transporter | chromosome:981590-983153(+) | | | | | chromosome:982137-982138(+):+0.207072[0.22857143,0.43564355] | | | | | -0.175042 | 2.52132 | -0.266315 |

| | | | | | | | | | | | | | | |
|---|---|---|---|---|---|---|---|---|---|---|---|---|---|---|
| gene;ACNSPI_05100 | 16S ribosomal RNA | chromosome:1006426-1007978(+) | |||chromosome:1006820-1006821(-):-0.256757[0.7567568,0.5];chromosome:1007467-1007468(-):+0.267692[0.04,0.30769232];chromosome:1007839-1007840(-):+0.326316[0.47368422,0.8] | | | |||chromosome:1007619-1007620(-):+0.270175[0.06315789,0.33333334] | | | chromosome:1007052-1007053(+):-0.211174[0.24242425,0.03125]|||chromosome:1007320-1007321(-):+0.207143[0.25,0.45714286] | chromosome:1007369-1007370(+):-0.35[0.85,0.5] | | -0.142479 | -0.181064 | 0.176056 |
| gene;ACNSPI_05105 | 23S ribosomal RNA | chromosome:1008282-1011205(+) | chromosome:1009653-1009654(+):+0.32266[0.03448276,0.35714287];chromosome:1010471-1010472(+):-0.2125[0.3125,0.1];chromosome:1010829-1010830(+):-0.270909[0.59090906,0.32];chromosome:1010925-1010926(+):+0.205742[0.6363636,0.84210527]|||chromosome:1008726-1008727(-):-0.295455[0.6818182,0.38636363];chromosome:1009205-1009206(-):+0.201139[0.4117647,0.61290324] | | | |||chromosome:1009206-1009207(-):-0.222222[0.33333334,0.11111111];chromosome:1010147-1010148(-):+0.310247[0.20588236,0.516129] | | | chromosome:1010146-1010147(+):+0.211765[0.2,0.4117647]|||chromosome:1008820-1008821(-):-0.20303[0.36666667,0.16363636];chromosome:1010147-1010148(-):-0.24[0.74,0.5];chromosome:1010365-1010366(-):+0.329412[0.47058824,0.8];chromosome:1010465-1010466(-):+0.219048[0.06666667,0.2857143] | | | -1.28537 | 1.03177 | -0.666824 |
| gene;ACNSPI_05130 | tRNA-Tyr | chromosome:1011662-1011746(+) | | | | | | | chromosome:1011736-1011737(+):+0.25317[0.23958333,0.49275362] | | | | | |
| gene;ACNSPI_05175 | LPXTG-anchored DUF1542 repeat protein FmtB | chromosome:1018063-1025500(+) | |||chromosome:1018449-1018450(-):-0.342857[0.64285713,0.3] | | | | | | | | | | | |

| | | | | | | | | | | | | | | |
|---|---|---|---|---|---|---|---|---|---|---|---|---|---|---|
| gene;ACNSPI_05305 | ATP-grasp domain-containing protein | chromosome:1053551-1054745(+) | | | | chromosome:1054086-1054087(+):+0.26455[0.14285715,0.4074074] | | | | | | -1.01463 | 1.04577 | -0.581932 |
| gene;ACNSPI_05315 | type II pantothenate kinase | chromosome:1055944-1056748(-) | | | | | | \|\|\|chromosome:1056247-1056248(+):+0.259259[0.074074075,0.33333334] | | | | 0.123692 | -1.16029 | -0.0629448 |
| gene;ACNSPI_05470 | UDP-N-acetylglucosamine 1-carboxyvinyltransferase | chromosome:1084145-1085411(+) | | | | | | chromosome:1084810-1084811(+):+0.210053[0.20930232,0.41935483] | | | | -0.268536 | -0.379973 | 0.233642 |
| gene;ACNSPI_05475 | 3-hydroxyacyl-ACP dehydratase FabZ | chromosome:1085444-1085885(+) | chromosome:1085601-1085602(+):-0.274155[0.3611111,0.08695652] | | | | | | | | | -0.280448 | -0.608396 | 0.14105 |
| gene;ACNSPI_05490 | single-stranded DNA-binding protein | chromosome:1086577-1086972(+) | | | chromosome:1086790-1086791(+):+0.206774[0.15686275,0.36363637] | | | | | | | | | |
| gene;ACNSPI_05510 | thiamine phosphate synthase | chromosome:1090731-1091373(+) | chromosome:1090778-1090779(+):+0.25[0.33333334,0.5833333] | | | \|\|\|chromosome:1091153-1091154(-):+0.35101[0.19444445,0.54545456] | | | | | | 0.577698 | 0.722886 | 0.748467 |
| gene;ACNSPI_05665 | tRNA-Leu | chromosome:1122439-1122522(+) | | | | | | | chromosome:1122493-1122494(+):+0.202857[0.15714286,0.36]\|\|\|chromosome:1122450-1122451(-):+0.261905[0.6666667,0.9285714] | | | 3.9244 | -1.22969 | -1.89902 |
| gene;ACNSPI_05675 | 16S ribosomal RNA | chromosome:1122736-1124288(+) | chromosome:1124281-1124282(+):-0.243961[0.5217391,0.2777778] | | | \|\|\|chromosome:1123629-1123630(-):+0.3[0.5,0.8];chromosome:1124012-1124013(-):+0.211348[0.25531915,0.46666667] | | | chromosome:1123675-1123676(+):-0.341176[0.9,0.5588235];chromosome:1124211-1124212(+):-0.280702[0.28070176,0] | | | -0.930647 | 1.3211 | -0.0290437 |

| | | | | | | | | | | | | | | |
|---|---|---|---|---|---|---|---|---|---|---|---|---|---|---|
| gene;ACNSPI_05680 | 23S ribosomal RNA | chromosome:1124592-1127515(+) | chromosome:1126455-1126456(+):+0.234848[0.5833333,0.8181818];chromosome:1127204-1127205(+):+0.214286[0.2857143,0.5]\|\|\|chromosome:1125516-1125517(-):+0.233333[0.1,0.33333334];chromosome:1125986-1125987(-):-0.29[0.65,0.36];chromosome:1126457-1126458(-):-0.232143[0.60714287,0.375] | | | chromosome:1125994-1125995(+):+0.26046[0.0862069,0.34666666];chromosome:1127117-1127118(+):-0.274131[0.7027027,0.42857143];chromosome:1127138-1127139(+):+0.235993[0.29032257,0.5263158]\|\|\|chromosome:1126675-1126676(-):-0.305556[0.5555556,0.25];chromosome:1126869-1126870(-):-0.256944[0.6944444,0.4375] | | | \|\|\|chromosome:1125986-1125987(-):-0.274945[0.36585367,0.09090909];chromosome:1126775-1126776(-):+0.246454[0.17021276,0.41666666] | | | -0.949407 | 0.309904 | -2.56639 |
| gene;ACNSPI_05695 | 3-isopropylmalate dehydratase large subunit | chromosome:1129767-1131138(-) | chromosome:1129915-1129916(-):+0.279245[0.3207547,0.6] | | | | | | | | | -0.0232309 | 1.49904 | -2.11906 |
| gene;ACNSPI_05750 | tRNA (adenosine(37)-N6)-threonylcarbamoyltransferase complex transferase subunit TsaD | chromosome:1140684-1141710(+) | \|\|\|chromosome:1140885-1140886(-):-0.233333[0.5,0.26666668] | | | | | | | | | -0.0490433 | -1.17944 | 0.580981 |
| gene;ACNSPI_05775 | YeeE/YedE family protein | chromosome:1146902-1147982(+) | chromosome:1147794-1147795(+):-0.335968[0.6086956,0.27272728] | | | | | | | | | -1.23569 | -0.785827 | -0.305249 |
| gene;ACNSPI_05790 | LacI family DNA-binding transcriptional regulator | chromosome:1149906-1150857(+) | chromosome:1150714-1150715(+):-0.209302[0.20930232,0] | | | | | | | | | -0.359056 | -0.461882 | 0.202651 |
| gene;ACNSPI_05845 | SdrH family protein | chromosome:1159507-1160761(+) | chromosome:1160090-1160091(+):+0.227575[0.21428572,0.44186047] | | | | | | | | | 0.12742 | 0.670063 | -0.672592 |

<table>
<tr><td></td><td></td><td></td><td></td><td></td><td></td><td></td><td></td><td></td><td></td><td></td><td></td><td></td><td></td><td></td></tr>
<tr><td>gene;ACNSPI_05890</td><td>TrkH family potassium uptake protein</td><td>chromosome:1167336-1168644(-)</td><td>chromosome:1167757-1167758(-):-0.363821[0.5833333,0.2195122]</td><td></td><td></td><td></td><td></td><td></td><td></td><td></td><td></td><td>0.0660154</td><td>-1.07504</td><td>-0.594268</td></tr>
<tr><td>gene;ACNSPI_05965</td><td>MW1434 family type I TA system toxin</td><td>chromosome:1180690-1180906(+)</td><td>chromosome:1180842-1180843(+):-0.240909[0.75,0.5090909];chromosome:1180845-1180846(+):+0.219958[0.13636364,0.35632184]|||chromosome:1180853-1180854(-):+0.221154[0.15384616,0.375]</td><td></td><td></td><td></td><td></td><td></td><td>|||chromosome:1180853-1180854(-):-0.281513[0.3529412,0.071428575]</td><td></td><td></td><td>4.41839</td><td>-0.740678</td><td>-0.369223</td></tr>
<tr><td>gene;ACNSPI_06045</td><td>RusA family crossover junction endodeoxyribonuclease</td><td>chromosome:1189109-1189514(+)</td><td>chromosome:1189167-1189168(+):-0.232278[0.4117647,0.17948718]</td><td></td><td></td><td></td><td></td><td></td><td></td><td></td><td></td><td>3.41967</td><td>-0.17211</td><td>0.129376</td></tr>
<tr><td>gene;ACNSPI_06095</td><td>transcriptional regulator</td><td>chromosome:1192372-1192522(+)</td><td>chromosome:1192484-1192485(+):+0.291667[0.375,0.6666667]</td><td></td><td></td><td></td><td></td><td></td><td></td><td></td><td></td><td>-0.212817</td><td></td><td></td></tr>
<tr><td>gene;ACNSPI_06105</td><td>transcriptional regulator</td><td>chromosome:1192749-1193166(+)</td><td></td><td></td><td></td><td>chromosome:1192906-1192907(+):-0.248054[0.6097561,0.3617021]</td><td></td><td></td><td></td><td></td><td></td><td>3.40255</td><td>-0.171917</td><td>0.0234911</td></tr>
<tr><td>gene;ACNSPI_06185</td><td>phage tail tape measure protein</td><td>chromosome:1202206-1206736(+)</td><td></td><td></td><td></td><td>chromosome:1205061-1205062(+):+0.237143[0.12,0.35714287]</td><td></td><td>chromosome:1203016-1203017(+):-0.213636[0.23636363,0.022727273]</td><td>chromosome:1205681-1205682(+):+0.218402[0.13043478,0.3488372]</td><td></td><td></td><td>3.40926</td><td>-0.425477</td><td>-0.770683</td></tr>
<tr><td>gene;ACNSPI_06190</td><td>phage tail domain-containing protein</td><td>chromosome:1206732-1208217(+)</td><td></td><td></td><td></td><td></td><td></td><td></td><td>chromosome:1207498-1207499(+):-0.209596[0.2777778,0.06818182]</td><td></td><td></td><td>3.34211</td><td>-0.857005</td><td>-0.51449</td></tr>
</table>

| | | | | | | | | | | | | | | |
|---|---|---|---|---|---|---|---|---|---|---|---|---|---|---|
| gene;ACNSPI_06715 | 16S ribosomal RNA | chromosome:1300731-1302283(+) | 1585-1301586(+):-0.210407[0.44117647,0.23076923];chromosome:1301794-1301795(+):-0.285714[0.42857143,0.14285715];chromosome:1301811-1301812(+):-0.203636[0.36363637,0.16];chromosome:1302276-1302277(+):-0.215517[0.46551725,0.25]\|\|\|chromosome:1301056-1301057(-):-0.308316[0.4117647,0.10344828];chromosome:1301573-1301574(-):+0.260307[0.44339624,0.7037037];chromosome:1301624-1301625(-):-0.568182[0.75,0. | | | \|\|\|chromosome:1301125-1301126(-):+0.320296[0.47457626,0.7948718] | | | \|\|\|chromosome:1301039-1301040(-):+0.307566[0.21875,0.5263158];chromosome:1301625-1301626(-):+0.222222[0.33333334,0.5555556] | | | -1.15527 | 0.862734 | 0.351627 |
| gene;ACNSPI_06720 | tRNA-Ile | chromosome:1302374-1302451(+) | chromosome:1302436-1302437(+):-0.207143[0.6,0.39285713] | | | chromosome:1302435-1302436(+):-0.222222[0.22222222,0] | | | | | | -1.39863 | 0.257417 | 0.519571 |
| gene;ACNSPI_06725 | tRNA-Ala | chromosome:1302469-1302545(+) | \|\|\|chromosome:1302486-1302487(-):-0.316667[0.73333335,0.41666666] | | | chromosome:1302516-1302517(+):-0.244186[0.5,0.25581396] | | | chromosome:1302516-1302517(+):+0.222591[0.3488372,0.5714286] | | | -0.445483 | 0.130338 | 0.254498 |

<table>
<tr><td>gene;ACNSPI_06730</td><td>23S ribosomal RNA</td><td>chromosome:1302758-1305681(+)</td><td>chromosome:1303686-1303687(+):+0.214286[0,0.21428572];chromosome:1305304-1305305(+):-0.212644[0.37931034,0.16666667];chromosome:1305401-1305402(+):+0.214286[0.5,0.71428573]|||chromosome:1303693-1303694(-):-0.252174[0.6,0.3478261];chromosome:1305319-1305320(-):+0.262931[0.36206895,0.625]</td><td>chromosome:1303971-1303972(+):-0.212454[0.30769232,0.0952381];chromosome:1304849-1304850(+):-0.223077[0.3,0.07692308]</td><td></td><td>4231-1304232(+):+0.280702[0.05263158,0.33333334];chromosome:1305010-1305011(+):-0.337607[0.7222222,0.3846154];chromosome:1305323-1305324(+):-0.269231[0.5,0.23076923]|||chromosome:1304236-1304237(-):+0.210606[0.1,0.31060606];chromosome:1304623-1304624(-):+0.260417[0.40625,0.6666667];chromosome:1304841-1304842(-):-0.333333[0.6,0.26666668];chromosome:1304940-1304941(-):-0.214286[0.31428573,0.1];chromosome:1304941-1304942(-</td><td></td><td></td><td>chromosome:1304617-1304618(+):+0.268421[0.6315789,0.9];chromosome:1304947-1304948(+):+0.411765[0,0.4117647];chromosome:1305010-1305011(+):-0.33913[0.73913044,0.4];chromosome:1305045-1305046(+):+0.253968[0.04761905,0.3015873];chromosome:1305318-1305319(+):+0.201299[0.72727275,0.9285714];chromosome:1305395-1305396(+):-0.209694[0.6712329,0.46153846]</td><td></td><td></td><td>-1.17992</td><td>1.55063</td><td>0.281765</td></tr>
<tr><td>gene;ACNSPI_06745</td><td>tRNA-Thr</td><td>chromosome:1305971-1306047(+)</td><td></td><td></td><td></td><td>chromosome:1306031-1306032(+):+0.537838[0.16216215,0.7]</td><td></td><td></td><td></td><td></td><td></td><td></td><td></td><td></td></tr>
<tr><td>gene;ACNSPI_06755</td><td>tRNA-Leu</td><td>chromosome:1306131-1306213(+)</td><td></td><td></td><td></td><td></td><td></td><td></td><td>chromosome:1306182-1306183(+):+0.201544[0.20754717,0.4090909]</td><td></td><td></td><td></td><td></td><td></td></tr>
<tr><td>gene;ACNSPI_06760</td><td>tRNA-Gly</td><td>chromosome:1306216-1306291(+)</td><td></td><td></td><td></td><td>|||chromosome:1306265-1306266(-):-0.213904[0.77272725,0.5588235]</td><td></td><td></td><td></td><td></td><td></td><td></td><td></td><td></td></tr>
</table>

| | | | | | | | | | | | | |
|---|---|---|---|---|---|---|---|---|---|---|---|---|
| gene;ACNSPI_06765 | tRNA-Leu | chromosome:1306301-1306390(+) | \|\|\|chromosome:1306318-1306319(-):+0.200669[0.53846157,0.73913044] | | | | | | chromosome:1306316-1306317(+):+0.279279[0.054054055,0.33333334];chromosome:1306369-1306370(+):-0.322344[0.84615386,0.52380955]\|\|\|chromosome:1306318-1306319(-):+0.311111[0.6333333,0.9444444] | | | |
| gene;ACNSPI_06780 | tRNA-Ala | chromosome:1306582-1306658(+) | \|\|\|chromosome:1306599-1306600(-):+0.251765[0.16,0.4117647] | | | \|\|\|chromosome:1306599-1306600(-):-0.318681[0.46153846,0.14285715] | | | | | | |
| gene;ACNSPI_06785 | tRNA-Met | chromosome:1306679-1306753(+) | chromosome:1306741-1306742(+):-0.406593[0.6923077,0.2857143] | | | chromosome:1306741-1306742(+):-0.336364[0.7,0.36363637] | | | | | | |
| gene;ACNSPI_06795 | tRNA-Ser | chromosome:1306864-1306954(+) | \|\|\|chromosome:1306931-1306932(-):+0.230769[0.07692308,0.30769232] | | | | | | | | | |
| gene;ACNSPI_06800 | tRNA-Asp | chromosome:1306964-1307040(+) | | | | \|\|\|chromosome:1307005-1307006(-):-0.222763[0.3548387,0.13207547] | | | | | | |
| gene;ACNSPI_06805 | tRNA-Ser | chromosome:1307078-1307171(+) | | | | chromosome:1307150-1307151(+):-0.518182[0.8181818,0.3]\|\|\|chromosome:1307145-1307146(-):-0.203661[0.42105263,0.2173913] | | | | | | 0.734174 |

<table>
<tr><th></th><th></th><th></th><th></th><th></th><th></th><th></th><th></th><th></th><th></th><th></th><th></th><th></th><th></th><th></th></tr>
<tr><td>gene;ACNSPI_06810</td><td>tRNA-Met</td><td>chromosome:1307185-1307259(+)</td><td>chromosome:1307226-1307227(+):-0.333333[0.7,0.36666667]|||chromosome:1307203-1307204(-):+0.232063[0.25974026,0.4918033]</td><td></td><td></td><td></td><td></td><td></td><td></td><td></td><td></td><td></td><td></td><td></td></tr>
<tr><td>gene;ACNSPI_06830</td><td>tRNA-Tyr</td><td>chromosome:1307523-1307604(+)</td><td>|||chromosome:1307579-1307580(-):+0.212474[0.13636364,0.3488372]</td><td></td><td></td><td></td><td></td><td></td><td></td><td></td><td></td><td></td><td></td><td></td></tr>
<tr><td>gene;ACNSPI_06835</td><td>tRNA-Trp</td><td>chromosome:1307619-1307693(+)</td><td></td><td></td><td></td><td></td><td></td><td></td><td>chromosome:1307678-1307679(+):+0.229651[0.375,0.60465115]</td><td></td><td></td><td></td><td></td><td></td></tr>
<tr><td>gene;ACNSPI_06840</td><td>tRNA-His</td><td>chromosome:1307695-1307768(+)</td><td>chromosome:1307735-1307736(+):+0.212815[0.26086956,0.47368422]</td><td></td><td></td><td></td><td></td><td></td><td></td><td></td><td></td><td></td><td></td><td></td></tr>
<tr><td>gene;ACNSPI_06865</td><td>tRNA-Leu</td><td>chromosome:1308228-1308312(+)</td><td>|||chromosome:1308295-1308296(-):+0.344444[0.5555556,0.9]</td><td></td><td></td><td>|||chromosome:1308282-1308283(-):-0.254634[0.34,0.085365854]</td><td></td><td></td><td></td><td></td><td></td><td></td><td></td><td></td></tr>
<tr><td>gene;ACNSPI_06885</td><td>amino acid ABC transporter ATP-binding protein</td><td>chromosome:1313027-1313750(+)</td><td></td><td></td><td></td><td>|||chromosome:1313614-1313615(-):-0.201677[0.7111111,0.509434]</td><td></td><td></td><td></td><td></td><td></td><td>-0.639222</td><td>-1.49967</td><td>-0.549562</td></tr>
<tr><td>gene;ACNSPI_06925</td><td>GAF domain-containing sensor histidine kinase</td><td>chromosome:1320637-1321750(+)</td><td></td><td></td><td></td><td></td><td></td><td>chromosome:1321674-1321675(+):+0.225732[0.045454547,0.27118644]</td><td></td><td></td><td></td><td>0.654943</td><td>0.192729</td><td>-0.291436</td></tr>
<tr><td>gene;ACNSPI_06930</td><td>response regulator</td><td>chromosome:1321771-1322395(+)</td><td>chromosome:1321916-1321917(+):+0.200466[0.030303031,0.23076923]|||chromosome:1322167-1322168(-):+0.299663[0.18181819,0.4814815]</td><td></td><td>chromosome:1321818-1321819(+):-0.213665[0.25714287,0.04347826]</td><td></td><td></td><td>chromosome:1321818-1321819(+):+0.218102[0.27027026,0.4883721];chromosome:1322155-1322156(+):-0.20595[0.68421054,0.47826087]</td><td></td><td></td><td></td><td>0.350477</td><td>0.142033</td><td>-0.137615</td></tr>
</table>

<table>
<tr><td>gene;ACNSPI_06970</td><td>peptidylprolyl isomerase</td><td>chromosome:1330883-1331846(-)</td><td>|||chromosome:1330897-1330898(+):+0.321839[0.3448276,0.6666667]</td><td></td><td></td><td></td><td></td><td></td><td></td><td></td><td></td><td>0.245533</td><td>3.04373</td><td>-0.205272</td></tr>
<tr><td>gene;ACNSPI_07065</td><td>tRNA-Glu</td><td>chromosome:1342568-1342640(+)</td><td>chromosome:1342573-1342574(+):-0.238095[0.23809524,0]</td><td></td><td></td><td></td><td></td><td></td><td></td><td></td><td></td><td></td><td></td><td></td></tr>
<tr><td>gene;ACNSPI_07140</td><td>DUF4888 domain-containing protein</td><td>chromosome:1358202-1358769(-)</td><td></td><td></td><td></td><td>|||chromosome:1358283-1358284(+):+0.287477[0.31858408,0.6060606]</td><td></td><td></td><td></td><td></td><td></td><td>2.32194</td><td>-0.147971</td><td>-0.192278</td></tr>
<tr><td>gene;ACNSPI_07255</td><td>DUF4352 domain-containing protein</td><td>chromosome:1378701-1379697(-)</td><td></td><td>chromosome:1379568-1379569(-):-0.227273[0.5,0.27272728]</td><td></td><td></td><td></td><td></td><td></td><td></td><td></td><td>-0.804507</td><td>-0.128065</td><td>-0.154682</td></tr>
<tr><td>gene;ACNSPI_07315</td><td>aldo/keto reductase</td><td>chromosome:1390769-1391603(+)</td><td>|||chromosome:1390834-1390835(-):+0.276819[0.4651163,0.7419355]</td><td></td><td></td><td></td><td></td><td></td><td></td><td></td><td></td><td>-0.437747</td><td>0.552265</td><td>0.391843</td></tr>
<tr><td>gene;ACNSPI_07415</td><td>proline dehydrogenase family protein</td><td>chromosome:1408604-1409606(-)</td><td>|||chromosome:1409522-1409523(+):-0.219512[0.2195122,0]</td><td></td><td></td><td>chromosome:1409529-1409530(-):+0.256966[0.05882353,0.31578946]</td><td></td><td></td><td></td><td></td><td></td><td>-0.950956</td><td>0.527425</td><td>0.634709</td></tr>
<tr><td>gene;ACNSPI_07445</td><td>TIGR01212 family radical SAM protein</td><td>chromosome:1413326-1414280(-)</td><td></td><td></td><td></td><td>chromosome:1414158-1414159(-):+0.208064[0.05,0.2580645]</td><td></td><td></td><td></td><td></td><td></td><td>-0.50648</td><td>0.762577</td><td>0.404513</td></tr>
<tr><td>gene;ACNSPI_07450</td><td>MDR family MFS transporter</td><td>chromosome:1414389-1415571(+)</td><td></td><td></td><td></td><td>chromosome:1414820-1414821(+):+0.3[0.3,0.6]</td><td></td><td></td><td>|||chromosome:1414764-1414765(-):-0.230519[0.32142857,0.09090909]</td><td></td><td></td><td>-0.00213862</td><td>-1.04297</td><td>0.439513</td></tr>
<tr><td>gene;ACNSPI_07590</td><td>acetate--CoA ligase</td><td>chromosome:1452550-1454257(+)</td><td></td><td></td><td></td><td></td><td></td><td></td><td>|||chromosome:1453942-1453943(-):+0.203911[0.33846155,0.5423729];chromosome:1454124-1454125(-):+0.298693[0.32352942,0.62222224]</td><td></td><td></td><td>-0.376642</td><td>1.18894</td><td>0.615328</td></tr>
</table>

| | | | | | | | | | | | | | | |
|---|---|---|---|---|---|---|---|---|---|---|---|---|---|---|
| gene;ACNSPI_07595 | formate--tetrahydrofolate ligase | chromosome:1454696-1456364(+) | chromosome:1454892-1454893(+):-0.238095[0.5714286,0.33333334] | | | | | | | | | -0.568768 | -0.638418 | 0.05022 |
| gene;ACNSPI_07905 | ATP-dependent Clp protease ATP-binding subunit ClpX | chromosome:1527628-1528891(+) | \|\|\|chromosome:1528300-1528301(-):+0.289744[0.07692308,0.36666667] | | | | | | | | | -0.43605 | -0.782794 | 0.373055 |
| gene;ACNSPI_07915 | glutamyl-tRNA reductase | chromosome:1529851-1531198(+) | chromosome:1530878-1530879(+):+0.270833[0.0625,0.33333334] | | | | | | | | | -0.255762 | -0.197793 | -0.0315608 |
| gene;ACNSPI_07975 | DNA repair protein RadC | chromosome:1543346-1544033(+) | \|\|\|chromosome:1543880-1543881(-):-0.2174[0.3580247,0.140625] | | | | | | | | | 0.958683 | 2.09724 | -0.0339513 |
| gene;ACNSPI_08065 | single-stranded-DNA-specific exonuclease RecJ | chromosome:1557185-1559459(+) | | | | | | | chromosome:1557528-1557529(+):+0.227848[0.1392405,0.36708862] | | | 0.0584542 | -1.03821 | 0.0459915 |
| gene;ACNSPI_08075 | bifunctional (p)ppGpp synthetase/guanosine-3'%2C5'-bis(diphosphate) 3'-pyrophosphohydrolase | chromosome:1560426-1562616(+) | chromosome:1561663-1561664(+):+0.215311[0.36363637,0.57894737] | | | | | | | | | -0.298867 | 0.4002 | -0.183046 |
| gene;ACNSPI_08140 | cysteine desulfurase family protein | chromosome:1572662-1573805(+) | \|\|\|chromosome:1572777-1572778(-):-0.402597[0.54545456,0.14285715] | | | | | | | | | 0.309217 | -0.131749 | 0.481485 |
| gene;ACNSPI_08160 | alanine--tRNA ligase | chromosome:1578875-1581506(+) | chromosome:1581419-1581420(+):-0.233987[0.4117647,0.17777778] | | | | | | | | | -0.527808 | -0.251743 | 0.347647 |
| gene;ACNSPI_08450 | DEAD/DEAH box helicase | chromosome:1634290-1635637(+) | | | | | | | \|\|\|chromosome:1635283-1635284(-):-0.240628[0.48387095,0.24324325] | | | -0.392111 | -0.677187 | 0.254574 |

| | | | | | | | | | | | | | | |
|---|---|---|---|---|---|---|---|---|---|---|---|---|---|---|
| gene;ACNSPI_08480 | peptidoglycan D%2CD-transpeptidase FtsI family protein | chromosome:1639746-1641822(+) | | chromosome:1641621-1641622(+):-0.260613[0.5625,0.3018868] | | | | | | | | -0.177445 | 0.585889 | -0.358407 |
| gene;ACNSPI_08570 | aminomethyl-transferring glycine dehydrogenase subunit GcvPB | chromosome:1653306-1654779(+) | | | | | | | \|\|\|chromosome:1654143-1654144(-):+0.200723[0.2278481,0.42857143] | | | -0.37315 | 0.189276 | -0.339362 |
| gene;ACNSPI_08610 | acetyl-CoA carboxylase biotin carboxylase subunit | chromosome:1660081-1661437(+) | chromosome:1660851-1660852(+):+0.370924[0.28125,0.65217394] | | | | | | | | | -0.454599 | 0.0362386 | 0.227457 |
| gene;ACNSPI_08620 | transcription antitermination factor NusB | chromosome:1661873-1662263(+) | | \|\|\|chromosome:1661937-1661938(-):+0.21627[0.3392857,0.5555556] | | | | | | | | -0.35889 | 0.0744205 | 0.107896 |
| gene;ACNSPI_08730 | pyrroline-5-carboxylate reductase | chromosome:1684554-1685370(-) | | | | \|\|\|chromosome:1684855-1684856(+):-0.47[0.72,0.25] | | | | | | -0.512315 | 1.59056 | -0.137004 |
| gene;ACNSPI_08755 | Fur family transcriptional regulator | chromosome:1688383-1688833(+) | | | | | | | chromosome:1688625-1688626(+):-0.232637[0.57746476,0.3448276] | | | 0.24569 | -1.12091 | -0.78984 |
| gene;ACNSPI_09065 | phage tail tape measure protein | chromosome:1726635-1732836(+) | chromosome:1727702-1727703(+):+0.242529[0.033333335,0.27586207];chromosome:1731106-1731107(+):+0.226891[0.05882353,0.2857143] | | chromosome:1730477-1730478(+):-0.318253[0.46341464,0.14516129];chromosome:1732665-1732666(+):-0.20097[0.2820513,0.08108108]\|\|\|chromosome:1731665-1731666(-):+0.20979[0.15384616,0.36363637] | \|\|\|chromosome:1731156-1731157(-):-0.21118[0.42857143,0.2173913];chromosome:1731497-1731498(-):-0.238095[0.32142857,0.083333336];chromosome:1731666-1731667(-):-0.235294[0.3529412,0.11764706] | | | | | | 0.755936 | 0.977326 | 0.3131 |
| gene;ACNSPI_09070 | phage tail family protein | chromosome:1732835-1733660(+) | | | chromosome:1733575-1733576(+):+0.200397[0.17460318,0.375] | | | | | | | 1.12801 | 0.966908 | 0.577901 |

<table>
<tr><th></th><th></th><th></th><th></th><th></th><th></th><th></th><th></th><th></th><th></th><th></th><th></th><th></th><th></th><th></th></tr>
<tr><td>gene;ACNSPI_09260</td><td>chorismate synthase</td><td>chromosome:1765250-1766417(+)</td><td>|||chromosome:1765339-1765340(-):+0.212197[0.10526316,0.31746033]</td><td></td><td></td><td></td><td></td><td></td><td></td><td></td><td></td><td>-0.457118</td><td>0.416062</td><td>0.223073</td></tr>
<tr><td>gene;ACNSPI_09305</td><td>CCA tRNA nucleotidyltransferase</td><td>chromosome:1774025-1775228(+)</td><td>chromosome:1774558-1774559(+):+0.213439[0.09090909,0.3043478]</td><td></td><td></td><td></td><td></td><td></td><td></td><td></td><td></td><td>-0.340771</td><td>-0.183264</td><td>0.165455</td></tr>
<tr><td>gene;ACNSPI_09320</td><td>asparagine--tRNA ligase</td><td>chromosome:1779224-1780517(+)</td><td></td><td></td><td></td><td>|||chromosome:1779882-1779883(-):-0.217334[0.3392857,0.12195122]</td><td></td><td></td><td></td><td></td><td></td><td>-0.326604</td><td>-0.322656</td><td>0.117261</td></tr>
<tr><td>gene;ACNSPI_09325</td><td>DnaD domain-containing protein</td><td>chromosome:1780844-1781531(+)</td><td></td><td></td><td>|||chromosome:1781135-1781136(-):-0.202369[0.7702703,0.56790125]</td><td></td><td></td><td></td><td></td><td></td><td></td><td>-0.424355</td><td>-0.679802</td><td>0.30254</td></tr>
<tr><td>gene;ACNSPI_09340</td><td>transglycosylase domain-containing protein</td><td>chromosome:1783091-1785275(-)</td><td>chromosome:1785060-1785061(-):+0.275862[0,0.27586207]</td><td></td><td></td><td></td><td></td><td></td><td></td><td></td><td></td><td>-0.309815</td><td>2.15876</td><td>0.329484</td></tr>
<tr><td>gene;ACNSPI_09385</td><td>PepSY-associated TM helix domain-containing protein</td><td>chromosome:1790119-1791460(-)</td><td>chromosome:1790716-1790717(-):+0.225624[0.1632653,0.3888889]</td><td></td><td></td><td></td><td></td><td></td><td></td><td></td><td></td><td>0.355168</td><td>-1.24167</td><td>1.60591</td></tr>
<tr><td>gene;ACNSPI_09415</td><td>multidrug efflux MFS transporter NorB</td><td>chromosome:1800339-1801731(+)</td><td></td><td></td><td></td><td>chromosome:1800785-1800786(+):+0.222222[0,0.22222222]</td><td></td><td></td><td></td><td></td><td></td><td>0.0327545</td><td>-0.409981</td><td>-0.306965</td></tr>
<tr><td>gene;ACNSPI_09420</td><td>hyperosmolarity resistance protein Ebh</td><td>chromosome:1802128-1833394(+)</td><td></td><td></td><td></td><td>|||chromosome:1808095-1808096(-):-0.22619[0.39285713,0.16666667]</td><td></td><td></td><td></td><td></td><td></td><td>0.250215</td><td>0.0403769</td><td>0.141976</td></tr>
<tr><td>gene;ACNSPI_09430</td><td>hypothetical protein</td><td>chromosome:1834198-1834330(+)</td><td>chromosome:1834204-1834205(+):+0.315942[0.2173913,0.53333336]</td><td></td><td></td><td></td><td></td><td></td><td>chromosome:1834206-1834207(+):-0.204503[0.5121951,0.30769232]</td><td></td><td></td><td></td><td></td><td></td></tr>
<tr><td>gene;ACNSPI_09540</td><td>2-oxoglutarate dehydrogenase E1 component</td><td>chromosome:1850230-1853029(+)</td><td>|||chromosome:1851244-1851245(-):-0.227273[0.5,0.27272728]</td><td>chromosome:1852942-1852943(+):-0.260163[0.33333334,0.07317073]</td><td></td><td></td><td></td><td></td><td></td><td></td><td></td><td>-0.610049</td><td>-0.079993</td><td>0.263047</td></tr>
</table>

<table>
<tr><td>gene;ACNSPI_09545</td><td>dihydrolipoyllysine-residue succinyltransferase</td><td>chromosome:1853042-1854311(+)</td><td></td><td></td><td></td><td></td><td></td><td></td><td>|||chromosome:1853539-1853540(-):+0.221269[0.41509435,0.6363636]</td><td></td><td></td><td>-0.560094</td><td>0.0445536</td><td>-0.287981</td></tr>
<tr><td>gene;ACNSPI_09565</td><td>hypothetical protein</td><td>chromosome:1856917-1858804(+)</td><td>|||chromosome:1857086-1857087(-):-0.308482[0.5294118,0.22093023]</td><td></td><td></td><td></td><td></td><td></td><td></td><td></td><td></td><td>-0.550428</td><td>-0.458075</td><td>0.620302</td></tr>
<tr><td>gene;ACNSPI_09665</td><td>phosphate ABC transporter substrate-binding protein PstS</td><td>chromosome:1877882-1878866(+)</td><td></td><td></td><td></td><td></td><td></td><td></td><td>chromosome:1877950-1877951(+):-0.200658[0.9375,0.7368421]</td><td></td><td></td><td>1.21231</td><td>3.08954</td><td>-0.218359</td></tr>
<tr><td>gene;ACNSPI_09710</td><td>nickel transporter permease</td><td>chromosome:1886233-1887064(+)</td><td>chromosome:1886891-1886892(+):+0.289474[0.5,0.7894737]|||chromosome:1886897-1886898(-):-0.205128[0.20512821,0]</td><td></td><td></td><td></td><td></td><td></td><td></td><td></td><td></td><td>0.520769</td><td>1.5673</td><td>0.59034</td></tr>
<tr><td>gene;ACNSPI_09765</td><td>anthranilate phosphoribosyltransferase</td><td>chromosome:1896824-1897823(-)</td><td></td><td></td><td></td><td></td><td></td><td></td><td>chromosome:1897587-1897588(-):+0.214912[0.36842105,0.5833333]</td><td></td><td></td><td>0.346296</td><td>0.45867</td><td>-0.69189</td></tr>
<tr><td>gene;ACNSPI_09785</td><td>prephenate dehydrogenase</td><td>chromosome:1901485-1902577(+)</td><td></td><td></td><td></td><td></td><td></td><td></td><td>|||chromosome:1902325-1902326(-):+0.229379[0.1923077,0.42168674]</td><td></td><td></td><td>1.64044</td><td>0.305113</td><td>0.280958</td></tr>
<tr><td>gene;ACNSPI_09800</td><td>LCP family protein</td><td>chromosome:1904396-1905380(-)</td><td></td><td></td><td></td><td>chromosome:1904705-1904706(-):+0.226601[0.3448276,0.5714286]</td><td></td><td></td><td></td><td></td><td></td><td>0.419301</td><td>2.50569</td><td>-0.445074</td></tr>
<tr><td>gene;ACNSPI_09830</td><td>alanine/glycine:cation symporter family protein</td><td>chromosome:1911925-1913386(-)</td><td>|||chromosome:1912510-1912511(+):-0.355556[0.7222222,0.36666667]</td><td></td><td></td><td></td><td></td><td></td><td></td><td></td><td></td><td>-0.382435</td><td>-0.504086</td><td>0.344904</td></tr>
<tr><td>gene;ACNSPI_10165</td><td>MIP/aquaporin family protein</td><td>chromosome:1977082-1977901(-)</td><td></td><td></td><td></td><td>chromosome:1977326-1977327(-):-0.222222[0.37037036,0.14814815]</td><td></td><td></td><td></td><td></td><td></td><td>-0.78216</td><td>0.445312</td><td>0.628419</td></tr>
</table>

| | | | | | | | | | | | | | | |
|---|---|---|---|---|---|---|---|---|---|---|---|---|---|---|
| gene;ACNSPI_10190 | RicAFT regulatory complex protein RicA family protein | chromosome:1984374-1984740(-) | \|\|\|chromosome:1984533-1984534(+):+0.224359[0.083333336,0.30769232] | | | | | | | | | -0.619693 | 0.149176 | 0.0112627 |
| gene;ACNSPI_10205 | 2-oxoacid:ferredoxin oxidoreductase subunit beta | chromosome:1986805-1987672(-) | | | | chromosome:1987622-1987623(-):+0.211429[0.42857143,0.64] | | | chromosome:1987622-1987623(-):+0.231482[0.25,0.4814815] | | | -0.362014 | -0.145442 | 0.697592 |
| gene;ACNSPI_10215 | TIGR00282 family metallophosphoesterase | chromosome:1989572-1990370(-) | \|\|\|chromosome:1990094-1990095(+):-0.236928[0.4722222,0.23529412] | | | | | | | | | -0.365516 | -1.3174 | -0.0831897 |
| gene;ACNSPI_10310 | translation initiation factor IF-2 | chromosome:2011705-2013823(-) | chromosome:2012120-2012121(-):+0.4[0.3,0.7] | | | | | | | | | -0.441635 | -1.26847 | 0.46772 |
| gene;ACNSPI_10330 | ribosome maturation factor RimP | chromosome:2015642-2016110(-) | \|\|\|chromosome:2015869-2015870(+):-0.234162[0.37209302,0.13793103] | | | | | | | | | -0.0125489 | -1.25672 | 0.240174 |
| gene;ACNSPI_10345 | RIP metalloprotease RseP | chromosome:2022696-2023983(-) | | | | \|\|\|chromosome:2023127-2023128(+):+0.276406[0.35820895,0.63461536] | | | | | | -0.328663 | 0.98036 | 0.375199 |
| gene;ACNSPI_10415 | type I DNA topoisomerase | chromosome:2035365-2037441(-) | chromosome:2036939-2036940(-):-0.200664[0.82857144,0.627907] | | | | | | | | | -0.0908453 | 1.03708 | 0.296535 |
| gene;ACNSPI_10460 | 50S ribosomal protein L19 | chromosome:2048367-2048718(-) | | | | \|\|\|chromosome:2048552-2048553(+):-0.304167[0.4375,0.13333334] | | | | | | -0.0920605 | -0.665943 | 0.198617 |
| gene;ACNSPI_10610 | bifunctional phosphopantothenoylcysteine decarboxylase/phosphopantothenate--cysteine ligase CoaBC | chromosome:2081151-2082351(-) | \|\|\|chromosome:2081465-2081466(+):-0.212982[0.44827586,0.23529412] | | | | | | | | | -0.286723 | 0.219451 | 0.0143329 |
| gene;ACNSPI_10735 | cell division protein FtsZ | chromosome:2107483-2108656(-) | \|\|\|chromosome:2108328-2108329(+):-0.299423[0.6111111,0.3116883] | | | | | | | | | -0.364826 | -0.19836 | -0.0946943 |

| | | | | | | | | | | | | | | |
|---|---|---|---|---|---|---|---|---|---|---|---|---|---|---|
| gene;ACNSPI_10810 | tRNA-Arg | chromosome:2122170-2122244(+) | | | | \|\|\|chromosome:2122221-2122222(-):-0.457143[0.6,0.14285715] | | | chromosome:2122232-2122233(+):+0.266667[0.53333336,0.8] | | | 1.05042 | 0.796143 | 0.978121 |
| gene;ACNSPI_11210 | transketolase C-terminal domain-containing protein | chromosome:2199579-2200557(-) | \|\|\|chromosome:2200196-2200197(+):-0.301099[0.6857143,0.3846154] | | | | | | \|\|\|chromosome:2200196-2200197(+):+0.224828[0.62,0.8448276] | | | -0.469226 | -0.348409 | 0.278942 |
| gene;ACNSPI_11265 | phosphoenolpyruvate--protein phosphotransferase | chromosome:2209970-2211689(-) | | | | | | | chromosome:2211237-2211238(-):+0.218112[0.3125,0.53061223] | | | -0.366655 | 0.176455 | 0.238574 |
| gene;ACNSPI_11300 | ABC transporter ATP-binding protein | chromosome:2217059-2218460(+) | \|\|\|chromosome:2217763-2217764(-):-0.262121[0.34545454,0.083333336] | | | | | | chromosome:2217753-2217754(+):+0.236702[0.63829786,0.875] | | | -0.56125 | 1.46098 | -0.0474688 |
| gene;ACNSPI_11310 | phosphoribosylamine--glycine ligase | chromosome:2219523-2220771(-) | | | chromosome:2220651-2220652(-):+0.205128[0,0.20512821] | | | \|\|\|chromosome:2220711-2220712(+):+0.258064[0,0.2580645] | | | | -0.411131 | -0.288292 | -0.73513 |
| gene;ACNSPI_11315 | bifunctional phosphoribosylaminoimidazolecarboxamide formyltransferase/IMP cyclohydrolase | chromosome:2220792-2222271(-) | chromosome:2220809-2220810(-):+0.217391[0,0.2173913] | | chromosome:2220932-2220933(-):+0.243902[0,0.24390244];chromosome:2221212-2221213(-):+0.322727[0.22727273,0.55] | | | | | | | -0.6268 | -0.422331 | -0.363361 |
| gene;ACNSPI_11325 | phosphoribosylformylglycinamidine cyclo-ligase | chromosome:2222854-2223883(-) | chromosome:2223110-2223111(-):+0.233397[0.4117647,0.6451613] | | | | | | | | | -0.816087 | -0.29201 | -0.181982 |
| gene;ACNSPI_11420 | bifunctional autolysin | chromosome:2242215-2245986(+) | | | | chromosome:2243461-2243462(+):+0.274854[0.44444445,0.71929824] | | | | | | -0.260954 | 0.255655 | 0.590088 |
| gene;ACNSPI_11595 | tRNA-Ser | chromosome:2271356-2271445(+) | chromosome:2271434-2271435(+):+0.254808[0.4375,0.6923077] | | | \|\|\|chromosome:2271428-2271429(-):+0.414474[0.375,0.7894737] | | | | | | 3.32704 | 1.34882 | 1.50654 |

| | | | | | | | | | | | | | | |
|---|---|---|---|---|---|---|---|---|---|---|---|---|---|---|
| gene;ACNSPI_11870 | helicase-exonuclease AddAB subunit AddA | chromosome:2334565-2338219(-) | chromosome:2338126-2338127(-):+0.234375[0,0.234375] | | | | | | | | | -0.113214 | 0.589717 | 0.238804 |
| gene;ACNSPI_11950 | Na+/H+ antiporter Mnh1 subunit B | chromosome:2358507-2358936(+) | | | | chromosome:2358626-2358627(+):+0.380952[0.2857143,0.6666667] | | | | | | -0.191439 | 1.05441 | -0.238143 |
| gene;ACNSPI_11955 | Na+/H+ antiporter Mnh1 subunit C | chromosome:2358935-2359277(+) | | | | chromosome:2359263-2359264(+):-0.23913[0.3478261,0.10869565] | | | | | | -0.269803 | 0.876455 | -0.231059 |
| gene;ACNSPI_11960 | Na+/H+ antiporter Mnh1 subunit D | chromosome:2359269-2360766(+) | chromosome:2360433-2360434(+):+0.279352[0.10526316,0.3846154]\|\|\|chromosome:2360220-2360221(-):+0.203571[0.071428575,0.275] | | | chromosome:2360144-2360145(+):+0.241071[0.1875,0.42857143] | | chromosome:2360247-2360248(+):-0.233333[0.5833333,0.35] | chromosome:2360144-2360145(+):-0.254386[0.42105263,0.16666667] | | | -0.14228 | 0.889658 | -0.21249 |
| gene;ACNSPI_11980 | FAD/NAD(P)-binding protein | chromosome:2362122-2363277(+) | chromosome:2363228-2363229(+):-0.206767[0.42105263,0.21428572] | | | | | | | | | 0.551066 | -0.00351339 | -0.950466 |
| gene;ACNSPI_12110 | NAD(P)H-dependent flavin oxidoreductase | chromosome:2383780-2384848(-) | \|\|\|chromosome:2384259-2384260(+):-0.25[0.41666666,0.16666667] | | | | | | \|\|\|chromosome:2383839-2383840(+):+0.269231[0.23076923,0.5];chromosome:2383846-2383847(+):+0.296154[0.15384616,0.45] | | | 0.150138 | 1.24464 | -0.0497249 |
| gene;ACNSPI_12135 | Fe-S cluster assembly sulfur transfer protein SufU | chromosome:2388458-2388923(-) | | | | | | chromosome:2388757-2388758(-):+0.208244[0.23423423,0.44247788] | | | | -0.272007 | -0.398242 | 0.249192 |
| gene;ACNSPI_12445 | transfer-messenger RNA | chromosome:2435382-2435744(-) | \|\|\|chromosome:2435721-2435722(+):-0.256158[0.82758623,0.5714286] | | | chromosome:2435714-2435715(-):-0.217054[0.33333334,0.11627907] | | | | | | 0.203945 | 0.904434 | 0.105812 |
| gene;ACNSPI_12770 | DNA helicase RecQ | chromosome:2507171-2508953(-) | | | | | | | chromosome:2508003-2508004(-):+0.3[0.3,0.6] | | | -0.0610127 | 0.434976 | 0.152102 |

| | | | | | | | | | | | | | | |
|---|---|---|---|---|---|---|---|---|---|---|---|---|---|---|
| gene;ACNSPI_12780 | polyglycerol-phosphate lipoteichoic acid synthase LtaS | chromosome:2511118-2513059(-) | | | \|\|\|chromosome:2512506-2512507(+):+0.217474[0.35042736,0.56790125] | | | | | | | 0.222912 | 1.00222 | -0.319654 |
| gene;ACNSPI_12840 | DoxX family membrane protein | chromosome:2521828-2522302(+) | \|\|\|chromosome:2522044-2522045(-):+0.201357[0.5294118,0.7307692] | | | | | | | | | 0.024495 | 1.11923 | -0.446147 |
| gene;ACNSPI_12875 | fructose-specific PTS transporter subunit EIIC | chromosome:2529324-2531283(-) | chromosome:2530966-2530967(-):-0.310924[0.42857143,0.11764706] | | | | | | chromosome:2530966-2530967(-):+0.20802[0.31578946,0.52380955] | | | -0.239652 | -0.0339338 | -0.274579 |
| gene;ACNSPI_12890 | Cys-tRNA(Pro) deacylase | chromosome:2533219-2533702(-) | chromosome:2533385-2533386(-):-0.242424[0.33333334,0.09090909] | | | chromosome:2533382-2533383(-):-0.206897[0.20689656,0] | | | | | | 0.248382 | 0.339002 | -0.248564 |
| gene;ACNSPI_12975 | TIGR00730 family Rossman fold protein | chromosome:2549714-2550281(+) | | | | chromosome:2550027-2550028(+):-0.22488[0.90909094,0.68421054] | | | | | | 0.191327 | 0.475038 | -0.0888593 |
| gene;ACNSPI_13130 | dihydroxyacetone kinase subunit DhaK | chromosome:2578020-2578989(-) | \|\|\|chromosome:2578829-2578830(+):+0.214912[0.41666666,0.6315789] | | | | | | | | | -0.56871 | 0.844232 | 1.05484 |
| gene;ACNSPI_13210 | metal-dependent transcriptional regulator | chromosome:2595427-2596072(-) | chromosome:2595698-2595699(-):-0.203007[0.6315789,0.42857143]\|\|\|chromosome:2595690-2595691(+):+0.203209[0.09090909,0.29411766] | | | | | | | | | 0.298359 | -0.603743 | -0.463478 |
| gene;ACNSPI_13465 | DUF443 family protein | chromosome:2641381-2642002(-) | | | | \|\|\|chromosome:2641447-2641448(+):+0.231579[0.36842105,0.6] | | | | | | 0.0632985 | -0.177717 | -0.0580376 |
| gene;ACNSPI_13670 | MSCRAMM family adhesin SdrC | chromosome:2685054-2687898(-) | \|\|\|chromosome:2685282-2685283(+):+0.209755[0.26785713,0.47761193] | | | \|\|\|chromosome:2685552-2685553(+):+0.302083[0.53125,0.8333333] | | | | | | 0.387891 | -2.15929 | -0.299623 |

| | | | | | | | | | | | | | | |
|---|---|---|---|---|---|---|---|---|---|---|---|---|---|---|
| gene;ACNSPI_13705 | branched-chain amino acid aminotransferase | chromosome:2692789-2693866(-) | | | | chromosome:2693296-2693297(-):+0.239011[0.15384616,0.39285713] | | | | | | -0.866204 | 0.819974 | -0.220183 |
| gene;ACNSPI_13775 | 50S ribosomal protein L7/L12 | chromosome:2714487-2714856(-) | chromosome:2714718-2714719(-):+0.229946[0.5882353,0.8181818] | | | | | | | | | -0.367147 | -1.68279 | 0.0468479 |
| gene;ACNSPI_13850 | DNA repair protein RadA | chromosome:2726118-2727483(-) | chromosome:2727193-2727194(-):-0.286765[0.4117647,0.125] | | | | | | | | | -0.0533731 | -0.062598 | 0.308253 |
| gene;ACNSPI_13905 | 23S ribosomal RNA | chromosome:2737709-2740632(-) | 7988-2737989(-):-0.203661[0.7826087,0.57894737];chromosome:2738104-2738105(-):-0.475[0.875,0.4];chromosome:2738106-2738107(-):+0.287356[0.33333334,0.62068963];chromosome:2738540-2738541(-):+0.20979[0.15384616,0.36363637];chromosome:2740517-2740518(-):+0.209239[0.3125,0.5217391]\|\|\|chromosome:2738062-2738063(+):+0.25[0,0.25];chromosome:2738449-2738450(+):-0.256685[0.5294118,0.27272728];chromosome:2738766-2738767(+):- | | | chromosome:2738379-2738380(-):-0.206349[0.7619048,0.5555556];chromosome:2739305-2739306(-):+0.267943[0.36363637,0.6315789]\|\|\|chromosome:2738354-2738355(+):+0.398649[0.35135135,0.75];chromosome:2739707-2739708(+):+0.253205[0.03846154,0.29166666] | | | chromosome:2737988-2737989(-):-0.203931[0.5675676,0.36363637];chromosome:2738084-2738085(-):+0.233333[0.1,0.33333334] | | | -0.847423 | 1.53133 | 0.272664 |

<table>
<tr><td>gene;ACNSPI_13910</td><td>16S ribosomal RNA</td><td>chromosome:2740936-2742488(-)</td><td></td><td></td><td></td><td>chromosome:2740963-2740964(-):+0.229677[0.29032257,0.52]|||chromosome:2741593-2741594(+):+0.269231[0.46153846,0.7307692]</td><td>chromosome:2741544-2741545(-):+0.270588[0.2,0.47058824]</td><td></td><td>chromosome:2742080-2742081(-):+0.2875[0.4,0.6875]|||chromosome:2741074-2741075(+):+0.232143[0.64285713,0.875]</td><td></td><td></td><td>-0.968995</td><td>1.14458</td><td>0.305218</td></tr>
<tr><td>gene;ACNSPI_13920</td><td>23S ribosomal RNA</td><td>chromosome:2742887-2745810(-)</td><td>chromosome:2743171-2743172(-):+0.25[0,0.25];chromosome:2743380-2743381(-):-0.222222[0.33333334,0.11111111];chromosome:2743522-2743523(-):+0.21448[0.13846155,0.3529412];chromosome:2743557-2743558(-):-0.266667[0.6666667,0.4]|||chromosome:2744885-2744886(+):-0.234604[0.36363637,0.12903225]</td><td></td><td></td><td>|||chromosome:2743627-2743628(+):+0.208539[0.1724138,0.3809524];chromosome:2743726-2743727(+):+0.361111[0.5555556,0.9166667];chromosome:2744874-2744875(+):+0.224026[0.20454545,0.42857143]</td><td></td><td></td><td>chromosome:2744875-2744876(-):+0.302564[0.46666667,0.7692308]|||chromosome:2745365-2745366(+):+0.201593[0.39215687,0.59375]</td><td></td><td></td><td>-1.09038</td><td>1.44294</td><td>0.530879</td></tr>
<tr><td>gene;ACNSPI_13925</td><td>tRNA-Ile</td><td>chromosome:2745977-2746054(-)</td><td></td><td></td><td></td><td>chromosome:2745991-2745992(-):-0.218301[0.5294118,0.31111112]</td><td></td><td></td><td>chromosome:2745991-2745992(-):+0.28882[0.2826087,0.5714286]</td><td></td><td></td><td>-1.1337</td><td>0.0491707</td><td>0.189601</td></tr>
</table>

<table>
<tr><th></th><th></th><th></th><th></th><th></th><th></th><th></th><th></th><th></th><th></th><th></th><th></th><th></th><th></th><th></th></tr>
<tr><td>gene;ACNSPI_13930</td><td>16S ribosomal RNA</td><td>chromosome:2746148-2747700(-)</td><td>chromosome:2747292-2747293(-):+0.25[0.33333334,0.5833333];chromosome:2747664-2747665(-):-0.35942[0.53333336,0.17391305]|||chromosome:2746423-2746424(+):-0.226382[0.5294118,0.3030303];chromosome:2746659-2746660(+):-0.25[0.7,0.45]</td><td></td><td></td><td>chromosome:2747292-2747293(-):+0.215385[0.4,0.61538464];chromosome:2747664-2747665(-):+0.26087[0,0.26086956]|||chromosome:2746807-2746808(+):+0.352941[0.64705884,1];chromosome:2747375-2747376(+):+0.210084[0.14285715,0.3529412]</td><td></td><td></td><td>chromosome:2746845-2746846(-):-0.208791[0.6730769,0.4642857]|||chromosome:2746658-2746659(+):+0.317073[0,0.31707317];chromosome:2746659-2746660(+):+0.525641[0.30769232,0.8333333]</td><td></td><td></td><td>-1.02789</td><td>1.37102</td><td>0.411081</td></tr>
<tr><td>gene;ACNSPI_13935</td><td>tRNA-Ala</td><td>chromosome:2747822-2747898(-)</td><td>chromosome:2747850-2747851(-):+0.222222[0,0.22222222];chromosome:2747870-2747871(-):-0.241667[0.825,0.5833333];chromosome:2747885-2747886(-):+0.351515[0.18181819,0.53333336]</td><td></td><td></td><td></td><td></td><td></td><td>|||chromosome:2747880-2747881(+):+0.215294[0.52,0.7352941]</td><td></td><td></td><td>1.09566</td><td>0.163605</td><td>-0.475051</td></tr>
<tr><td>gene;ACNSPI_13940</td><td>tRNA-Pro</td><td>chromosome:2747921-2747995(-)</td><td>|||chromosome:2747958-2747959(+):+0.259511[0.21875,0.47826087]</td><td></td><td></td><td></td><td></td><td></td><td></td><td></td><td></td><td></td><td></td><td></td></tr>
<tr><td>gene;ACNSPI_13950</td><td>tRNA-Leu</td><td>chromosome:2748097-2748186(-)</td><td></td><td></td><td></td><td>|||chromosome:2748174-2748175(+):+0.330189[0.5,0.8301887]</td><td></td><td></td><td></td><td></td><td></td><td></td><td></td><td></td></tr>
<tr><td>gene;ACNSPI_13955</td><td>tRNA-Gly</td><td>chromosome:2748193-2748268(-)</td><td>|||chromosome:2748218-2748219(+):+0.21875[0.40625,0.625];chromosome:2748225-2748226(+):-0.202083[0.46875,0.26666668]</td><td></td><td></td><td>chromosome:2748219-2748220(-):+0.209091[0.09090909,0.3]|||chromosome:2748225-2748226(+):-0.200855[0.42307693,0.22222222]</td><td></td><td></td><td></td><td></td><td></td><td></td><td></td><td></td></tr>
</table>

| | | | | | | | | | | | | | | |
|---|---|---|---|---|---|---|---|---|---|---|---|---|---|---|
| gene;ACNSPI_13990 | dihydroneopterin aldolase | chromosome:2751731-2752097(-) | chromosome:2751789-2751790(-):+0.277778[0.11111111,0.3888889] | | | | | | | | | -0.336715 | -0.427832 | 0.189332 |
| gene;ACNSPI_14010 | ATP-dependent zinc metalloprotease FtsH | chromosome:2755314-2757408(-) | | | | chromosome:2756994-2756995(-):-0.222222[0.4,0.17777778] | | | | | | -0.33719 | 0.336275 | 0.476378 |
| gene;ACNSPI_14065 | ribose-phosphate diphosphokinase | chromosome:2768760-2769726(-) | | | | \|\|\|chromosome:2769182-2769183(+):-0.204319[0.41860464,0.21428572] | | | | | | -0.274428 | -0.390289 | 0.0825488 |
| gene;ACNSPI_14095 | 4-(cytidine 5'-diphospho)-2-C-methyl-D-erythritol kinase | chromosome:2773177-2774026(-) | \|\|\|chromosome:2773728-2773729(+):-0.276973[0.4074074,0.13043478] | | | | | | | | | 0.219519 | 0.369709 | -0.491523 |
| gene;ACNSPI_14180 | 23S ribosomal RNA | chromosome:2787139-2790062(-) | chromosome:2787631-2787632(-):-0.237143[0.88,0.64285713];chromosome:2788202-2788203(-):+0.220833[0.46666667,0.6875] | | | chromosome:2787536-2787537(-):+0.246154[0.15384616,0.4] | | | chromosome:2787536-2787537(-):+0.21371[0.625,0.83870965];chromosome:2787631-2787632(-):+0.248252[0.6363636,0.88461536];chromosome:2789132-2789133(-):-0.20904[0.5423729,0.33333334]\|\|\|chromosome:2787878-2787879(+):+0.201238[0.0882353,0.28947368];chromosome:2787879-2787880(+):+0.264872[0.2244898,0.4893617] | | chromosome:2787657-2787658(-):-0.201839[0.76635516,0.5645161] | -0.647294 | 0.79373 | 0.380153 |

| | | | | | | | | | | | | | | |
|---|---|---|---|---|---|---|---|---|---|---|---|---|---|---|
| gene;ACNSPI_14185 | 16S ribosomal RNA | chromosome:2790428-2791980(-) | chromosome:2790904-2790905(-):+0.235294[0,0.23529412];chromosome:2791572-2791573(-):+0.202381[0.4642857,0.6666667]\|\|\|chromosome:2790566-2790567(+):+0.272727[0.36363637,0.6363636];chromosome:2791045-2791046(+):-0.240891[0.8947368,0.65384614] | | | chromosome:2791040-2791041(-):+0.223684[0.5263158,0.75]\|\|\|chromosome:2790939-2790940(+):+0.238462[0.3,0.53846157];chromosome:2791626-2791627(+):+0.402027[0.125,0.527027] | | | chromosome:2791128-2791129(-):+0.254372[0.21621622,0.47058824]\|\|\|chromosome:2791046-2791047(+):-0.32971[0.41666666,0.08695652];chromosome:2791085-2791086(+):+0.241071[0.32142857,0.5625];chromosome:2791585-2791586(+):+0.220034[0.4375,0.65753424];chromosome:2791655-2791656(+):+0.22824[0.09090909,0.31914893] | | | 0.0185868 | 1.0081 | 0.442043 |
| gene;ACNSPI_14240 | tRNA-Ser | chromosome:2801127-2801220(-) | chromosome:2801147-2801148(-):+0.258152[0.4375,0.6956522]\|\|\|chromosome:2801153-2801154(+):-0.217433[0.5277778,0.31034482] | | | | | | | | | 1.51937 | 1.54234 | -0.640062 |
| gene;ACNSPI_14240 | glutamate synthase subunit beta | chromosome:2801488-2802952(-) | | | | \|\|\|chromosome:2802511-2802512(+):+0.218474[0.12195122,0.34042552] | | | | | | -0.351208 | 2.34188 | -0.967865 |
| gene;ACNSPI_14245 | glutamate synthase large subunit | chromosome:2802969-2807469(-) | chromosome:2807322-2807323(-):+0.2125[0.1875,0.4]\|\|\|chromosome:2806777-2806778(+):-0.244939[0.55263156,0.30769232] | | | | | | chromosome:2807316-2807317(-):-0.208333[0.20833333,0];chromosome:2807322-2807323(-):+0.20761[0.17948718,0.38709676] | | | -1.35375 | 3.06566 | -0.887405 |
| gene;ACNSPI_14280 | autolysin/adhesin Aaa | chromosome:2812187-2813192(-) | \|\|\|chromosome:2812726-2812727(+):+0.219002[0.5217391,0.7407407] | | | | | | | | | 0.0811539 | -1.65117 | 1.05374 |

| | | | | | | | | | | | | | | |
|---|---|---|---|---|---|---|---|---|---|---|---|---|---|---|
| gene;ACNSPI_14340 | DUF2309 domain-containing protein | chromosome:2823086-2825792(-) | | | | chromosome:2823579-2823580(-):+0.264706[0.23529412,0.5] | | | chromosome:2823579-2823580(-):+0.35[0.15,0.5] | | | -0.0084627 | 1.94358 | 0.178382 |
| gene;ACNSPI_14505 | superantigen-like protein SSL4 | chromosome:2854127-2855054(-) | \|\|\|chromosome:2854964-2854965(+):+0.342995[0.4347826,0.7777778] | | | \|\|\|chromosome:2854964-2854965(+):+0.20202[0.57575756,0.7777778] | | | | | | 0.822472 | 2.82105E-05 | -0.23837 |
| gene;ACNSPI_14515 | superantigen-like protein SSL3 | chromosome:2855418-2856477(-) | \|\|\|chromosome:2856107-2856108(+):-0.228758[0.61764705,0.3888889] | | | \|\|\|chromosome:2856107-2856108(+):+0.229798[0.4090909,0.6388889] | | | | | | 0.791979 | 1.07449 | -0.831418 |
| gene;ACNSPI_14565 | IMP dehydrogenase | chromosome:2865043-2866510(-) | chromosome:2865189-2865190(-):+0.272941[0.08,0.3529412] | | | | | | | | | -0.163961 | -0.187529 | 0.686934 |
| gene;ACNSPI_14570 | xanthine permease PbuX | chromosome:2866547-2867816(-) | chromosome:2866653-2866654(-):+0.207046[0.36158192,0.5686275] | | | | | | | | | 0.234582 | -0.569422 | 0.718691 |
| gene;ACNSPI_14600 | L-cystine transporter | chromosome:2872169-2873558(+) | chromosome:2873325-2873326(+):+0.246667[0.23333333,0.48] | | | | | | | | | -0.153033 | -1.67181 | -0.22172 |
| terminator;1 | alkyl hydroperoxide reductase subunit C;alkyl hydroperoxide reductase subunit F | chromosome:486-2594(+) | | | | | | | chromosome:2136-2137(+):-0.27451[0.33333334,0.05882353] | | | -0.247435 to -0.24055 | -1.49652 to -1.42397 | +0.524968 to +0.551929 |
| terminator;1001 | prephenate dehydrogenase | chromosome:1901486-1902577(+) | | | | | | | \|\|\|chromosome:1902325-1902326(-):+0.229379[0.1923077,0.42168674] | | | +1.64044 to +1.64044 | +0.305113 to +0.305113 | +0.280958 to +0.280958 |
| terminator;1004 | LCP family protein | chromosome:1904397-1905380(-) | | | | chromosome:1904705-1904706(-):+0.226601[0.3448276,0.5714286] | | | | | | +0.419301 to +0.419301 | +2.50569 to +2.50569 | -0.445074 to -0.445074 |
| terminator;1008 | alanine/glycine:cation symporter family protein | chromosome:1911926-1913386(-) | \|\|\|chromosome:1912510-1912511(+):-0.355556[0.7222222,0.36666667] | | | | | | | | | -0.382435 to -0.382435 | -0.504086 to -0.504086 | +0.344904 to +0.344904 |

| | | | | | | | | | | | | | | |
|---|---|---|---|---|---|---|---|---|---|---|---|---|---|---|
| terminator;104 | MurR/RpiR family transcriptional regulator;PTS transporter subunit EIIC;N-acetylmuramic acid 6-phosphate etherase;6-phospho-N-acetylmuramidase | chromosome:202207-206502(-) | chromosome:203915-203916(-):-0.200669[0.46153846,0.26086956] | | | | | | | | | -0.603796 to +0.0354416 | -0.592446 to +0.198498 | -0.327153 to +0.893391 |
| terminator;105 | glucose-specific PTS transporter subunit IIBC | chromosome:207088-209133(+) | \|\|\|chromosome:207513-207514(-):-0.246956[0.5869565,0.34] | | | | | | | | | -0.850266 to -0.850266 | -0.450807 to -0.450807 | +0.411313 to +0.411313 |
| terminator;1052 | glycerol-3-phosphate dehydrogenase/oxidase;glycerol kinase GlpK;MIP/aquaporin family protein | chromosome:1973627-1977901(-) | | | | chromosome:1977326-1977327(-):-0.222222[0.37037036,0.14814815] | | | | | | -0.78216 to -0.593891 | +0.445312 to +1.14128 | +0.219915 to +0.668719 |
| terminator;1054 | energy coupling factor transporter S component ThiW;RicAFT regulatory complex protein RicA family protein;tRNA (N6-isopentenyl adenosine(37)-C2)-methylthiotransferase MiaB | chromosome:1983857-1986285(-) | \|\|\|chromosome:1984533-1984534(+):+0.224359[0.083333336,0.30769232] | | | | | | | | | -0.619693 to -0.44152 | -0.0298335 to +0.248505 | +0.0112627 to +0.160468 |
| terminator;1056 | 2-oxoacid:ferredoxin oxidoreductase subunit beta;2-oxoacid:acceptor oxidoreductase subunit alpha | chromosome:1986806-1989433(-) | | | | chromosome:1987622-1987623(-):+0.211429[0.42857143,0.64] | | | chromosome:1987622-1987623(-):+0.231482[0.25,0.4814815] | | | -0.362014 to -0.350014 | -0.145442 to -0.138642 | +0.697592 to +0.881514 |
| terminator;1057 | TIGR00282 family metallophosphoesterase | chromosome:1989573-1990370(-) | \|\|\|chromosome:1990094-1990095(+):-0.236928[0.4722222,0.23529412] | | | | | | | | | -0.365516 to -0.365516 | -1.3174 to -1.3174 | -0.0831897 to -0.0831897 |

<table>
<tr><th></th><th></th><th></th><th></th><th></th><th></th><th></th><th></th><th></th><th></th><th></th><th></th><th></th><th></th><th></th></tr>
<tr><td>terminator;1063</td><td>ribonuclease J;polyribonucleotide nucleotidyltransferase;30S ribosomal protein S15;riboflavin biosynthesis protein RibF;tRNA pseudouridine(55) synthase TruB;30S ribosome-binding factor RbfA;translation initiation factor IF-2;YlxQ family RNA-binding protein;YlxR family RNase P modulator;transcription termination factor NusA;ribosome maturation factor RimP</td><td>chromosome:2004139-2016110(-)</td><td>chromosome:2012120-2012121(-):+0.4[0.3,0.7]|||chromosome:2015869-2015870(+):-0.234162[0.37209302,0.13793103]</td><td></td><td></td><td></td><td></td><td></td><td></td><td></td><td></td><td>-0.475835 to +0.127065</td><td>-1.62529 to +0.323288</td><td>-0.171794 to +0.632799</td></tr>
<tr><td>terminator;1065</td><td>proline--tRNA ligase;RIP metalloprotease RseP;phosphatidate cytidylyltransferase;isoprenyl transferase;ribosome recycling factor;UMP kinase;translation elongation factor Ts;hypothetical protein;30S ribosomal protein S2</td><td>chromosome:2020974-2029439(-)</td><td></td><td></td><td></td><td>|||chromosome:2023127-2023128(+):+0.276406[0.35820895,0.63461536]</td><td></td><td></td><td></td><td></td><td></td><td>-0.444252 to +0.1912</td><td>-0.830711 to +1.27728</td><td>-0.145678 to +0.479257</td></tr>
<tr><td>terminator;1067</td><td>methylenetetrahydrofolate--tRNA-(uracil(54)-C(5))-methyltransferase (FADH(2)-oxidizing) TrmFO;type I DNA topoisomerase;DNA-processing protein DprA</td><td>chromosome:2033903-2038487(-)</td><td>chromosome:2036939-2036940(-):-0.200664[0.82857144,0.627907]</td><td></td><td></td><td></td><td></td><td></td><td></td><td></td><td></td><td>-0.254514 to +0.625492</td><td>+0.705186 to +1.76536</td><td>-0.181715 to +0.530363</td></tr>
</table>

<table>
<tr><td>terminator;1072</td><td>protein L19;tRNA (guanosine(37)-N1)-methyltransferase TrmD;ribosome maturation factor RimM;30S ribosomal protein S16;signal recognition particle protein;putative DNA-binding protein;signal recognition particle-docking protein FtsY;chromosome segregation protein SMC;ribonuclease III;acyl carrier protein;3-oxoacyl-[acyl-carrier-protein] reductase;ACP S-malonyltransferase;phosphate acyltransferase PlsX;transcription</td><td>chromosome:2048368-2062166(-)</td><td></td><td></td><td></td><td>|||chromosome:2048552-2048553(+):-0.304167[0.4375,0.13333334]</td><td></td><td></td><td></td><td></td><td></td><td>-0.769574 to -0.0920605</td><td>-2.10483 to -0.232354</td><td>-0.334612 to +1.27695</td></tr>
<tr><td>terminator;1078</td><td>primosomal protein N';bifunctional phosphopantothenoylcysteine decarboxylase/phosphopantothenate--cysteine ligase CoaBC</td><td>chromosome:2078744-2082351(-)</td><td>|||chromosome:2081465-2081466(+):-0.212982[0.44827586,0.23529412]</td><td></td><td></td><td></td><td></td><td></td><td></td><td></td><td></td><td>-0.33239 to -0.286723</td><td>+0.0143642 to +0.219451</td><td>+0.0143329 to +0.115988</td></tr>
</table>

| | | | | | | | | | | | | | | |
|---|---|---|---|---|---|---|---|---|---|---|---|---|---|---|
| terminator;1086 | cell division protein FtsZ;cell division protein FtsA;cell division protein FtsQ/DivIB;UDP-N-acetylmuramoyl-L-alanine--D-glutamate ligase;phospho-N-acetylmuramoyl-pentapeptide-transferase;penicillin-binding protein;cell division protein FtsL;16S rRNA (cytosine(1402)-N(4))-methyltransferase RsmH;division/cell wall cluster transcriptional repressor MraZ | chromosome:2107484-2118162(-) | \|\|\|chromosome:2108328-2108329(+):-0.299423[0.6111111,0.3116883] | | | | | | | | | -0.367098 to +0.498964 | -0.19836 to +0.630972 | -1.09948 to -0.0946943 |
| terminator;1091 | tRNA-Arg | chromosome:2122171-2122244(+) | | | | \|\|\|chromosome:2122221-2122222(-):-0.457143[0.6,0.14285715] | | | chromosome:2122232-2122233(+):+0.266667[0.53333336,0.8] | | | +1.05042 to +1.05042 | +0.796143 to +0.796143 | +0.978121 to +0.978121 |
| terminator;111 | YagU family protein | chromosome:218112-218591(+) | chromosome:218495-218496(+):-0.210526[0.21052632,0] | | | | | | | | | -0.469805 to -0.469805 | +0.853675 to +0.853675 | -0.586348 to -0.586348 |
| terminator;112 | 4'-phosphopantetheinyl transferase superfamily protein;non-ribosomal peptide synthetase | chromosome:218919-226751(-) | chromosome:224018-224019(-):+0.221429[0.17857143,0.4] | | | chromosome:224018-224019(-):+0.3125[0.25,0.5625]\|\|\|chromosome:224535-224536(+):+0.257143[0.1,0.35714287];chromosome:224708-224709(+):-0.210237[0.34210527,0.13186814] | | | | | | +0.648318 to +0.772194 | +0.671522 to +1.32236 | +0.160127 to +0.390159 |

| | | | | | | | | | | | | | | |
|---|---|---|---|---|---|---|---|---|---|---|---|---|---|---|
| terminator;1142 | dihydrolipoyl dehydrogenase;dihydrolipoamide acetyltransferase family protein;transketolase C-terminal domain-containing protein;pyruvate dehydrogenase (acetyl-transferring) E1 component subunit alpha | chromosome:2196787-2201673(-) | \|\|\|chromosome:2200196-2200197(+):-0.301099[0.6857143,0.3846154] | | | | | | \|\|\|chromosome:2200196-2200197(+):+0.224828[0.62,0.8448276] | | | -0.469226 to -0.285952 | -0.68573 to -0.348409 | +0.0528753 to +0.357024 |
| terminator;1148 | phosphoenolpyruvate--protein phosphotransferase;phosphocarrier protein HPr | chromosome:2209971-2211958(-) | | | | | | | chromosome:2211237-2211238(-):+0.218112[0.3125,0.53061223] | | | -0.366655 to -0.213315 | +0.032128 to +0.176455 | +0.21379 to +0.238574 |
| terminator;1152 | ECF transporter S component;ABC transporter ATP-binding protein;energy-coupling factor transporter transmembrane protein EcfT | chromosome:2216470-2219259(+) | \|\|\|chromosome:2217763-2217764(-):-0.262121[0.34545454,0.083333336] | | | | | | chromosome:2217753-2217754(+):+0.236702[0.63829786,0.875] | | | -0.56125 to -0.3328 | +1.09181 to +1.53411 | -0.121314 to -0.00558305 |

| | | | | | | | | | | | | | | |
|---|---|---|---|---|---|---|---|---|---|---|---|---|---|---|
| terminator;1153 | [illegible]mine--glycine ligase;bifunctional phosphoribosylaminoimidazolecarboxamide formyltransferase/IMP cyclohydrolase;phosphoribosylglycinamide formyltransferase;phosphoribosylformylglycinamidine cyclo-ligase;amidophosphoribosyltransferase;phosphoribosylformylglycinamidine synthase subunit PurL;phosphoribosylformylglycinamidine synthase I;phosphoribosylformylglycinamidine synthase subunit PurS;phosphoribosylaminoimidaz | chromosome:2219524-2230758(-) | chromosome:2220809-2220810(-):+0.217391[0,0.2173913];chromosome:2223110-2223111(-):+0.233397[0.4117647,0.6451613] | | chromosome:2220651-2220652(-):+0.205128[0,0.20512821];chromosome:2220932-2220933(-):+0.243902[0,0.24390244];chromosome:2221212-2221213(-):+0.322727[0.22727273,0.55] | | | \|\|\|chromosome:2220711-2220712(+):+0.258064[0,0.2580645] | | | | -1.01407 to -0.411131 | -0.686768 to -0.200232 | -0.786826 to -0.0151069 |
| terminator;1161 | bifunctional autolysin | chromosome:2242216-2245986(+) | | | | chromosome:2243461-2243462(+):+0.274854[0.44444445,0.71929824] | | | | | | -0.260954 to -0.260954 | +0.255655 to +0.255655 | +0.590088 to +0.590088 |
| terminator;1182 | tRNA-Asn;tRNA-Ser | chromosome:2271280-2271445(+) | chromosome:2271434-2271435(+):+0.254808[0.4375,0.6923077] | | | \|\|\|chromosome:2271428-2271429(-):+0.414474[0.375,0.7894737] | | | | | | +3.23843 to +3.32704 | +1.34882 to +5.24239 | -1.03431 to +1.50654 |
| terminator;119 | cation diffusion facilitator family transporter | chromosome:235436-236395(-) | | | | chromosome:235807-235808(-):+0.201243[0.08080808,0.2820513] | | | | | | +0.168162 to +0.168162 | -0.626488 to -0.626488 | -0.0623164 to -0.0623164 |
| terminator;1213 | helicase-exonuclease AddAB subunit AddA;helicase-exonuclease AddAB subunit AddB | chromosome:2334566-2341696(-) | chromosome:2338126-2338127(-):+0.234375[0,0.234375] | | | | | | | | | -0.248874 to -0.113214 | +0.589717 to +0.624849 | +0.238804 to +0.266356 |

| | | | | | | | | | | | | | | |
|---|---|---|---|---|---|---|---|---|---|---|---|---|---|---|
| terminator;1223 | Na+/H+ antiporter Mnh1 subunit A;Na+/H+ antiporter Mnh1 subunit B;Na+/H+ antiporter Mnh1 subunit C;Na+/H+ antiporter Mnh1 subunit D;Na+/H+ antiporter Mnh1 subunit E;Na+/H+ antiporter Mnh1 subunit F;Na+/H+ antiporter Mnh1 subunit G | chromosome:2356110-2361874(+) | chromosome:2360433-2360434(+):+0.279352[0.10526316,0.3846154]\|\|\|chromosome:2360220-2360221(-):+0.203571[0.071428575,0.275] | | | chromosome:2358626-2358627(+):+0.380952[0.2857143,0.6666667];chromosome:2359263-2359264(+):-0.23913[0.3478261,0.10869565];chromosome:2360144-2360145(+):+0.241071[0.1875,0.42857143] | | chromosome:2360247-2360248(+):-0.233333[0.5833333,0.35] | chromosome:2360144-2360145(+):-0.254386[0.42105263,0.16666667] | | | -0.269803 to -0.0543111 | +0.679203 to +1.05441 | -0.454642 to -0.103825 |
| terminator;1224 | FAD/NAD(P)-binding protein | chromosome:2362123-2363277(+) | chromosome:2363228-2363229(+):-0.206767[0.42105263,0.21428572] | | | | | | | | | +0.551066 to +0.551066 | -0.00351339 to -0.00351339 | -0.950466 to -0.950466 |
| terminator;1237 | NAD(P)H-dependent flavin oxidoreductase;HlyC/CorC family transporter | chromosome:2383781-2385902(-) | \|\|\|chromosome:2384259-2384260(+):-0.25[0.41666666,0.16666667] | | | | | | \|\|\|chromosome:2383839-2383840(+):+0.269231[0.23076923,0.5];chromosome:2383846-2383847(+):+0.296154[0.15384616,0.45] | | | +0.110421 to +0.150138 | +1.24464 to +1.73532 | -0.0497249 to +0.157242 |
| terminator;1240 | Fe-S cluster assembly protein SufB;Fe-S cluster assembly sulfur transfer protein SufU;cysteine desulfurase;Fe-S cluster assembly protein SufD;Fe-S cluster assembly ATPase SufC | chromosome:2386911-2392435(-) | | | | | | chromosome:2388757-2388758(-):+0.208244[0.23423423,0.44247788] | | | | -0.323648 to +0.0695922 | -0.398242 to -0.0965561 | +0.244976 to +0.423215 |

| | | | | | | | | | | | | | | |
|---|---|---|---|---|---|---|---|---|---|---|---|---|---|---|
| terminator;127 | phosphate/phosphite/phosphonate ABC transporter substrate-binding protein;phosphonate ABC transporter ATP-binding protein;phosphonate ABC transporter%2C permease protein PhnE;phnE_2 | chromosome:265686-269243(+) | \|\|\|chromosome:265972-265973(-):-0.245645[0.31707317,0.071428575] | | | | | | chromosome:268286-268287(+):-0.26186[0.44186047,0.18] | | | +1.29349 to +1.78431 | -0.497782 to +0.338704 | -0.197824 to +0.143778 |
| terminator;1315 | DNA helicase RecQ;ABC-F family ATP-binding cassette domain-containing protein | chromosome:2507172-2510848(-) | | | | | | | chromosome:2508003-2508004(-):+0.3[0.3,0.6] | | | -0.0610127 to +0.143088 | +0.434976 to +0.862431 | -0.0158555 to +0.152102 |
| terminator;1316 | polyglycerol-phosphate lipoteichoic acid synthase LtaS | chromosome:2511119-2513059(-) | | | \|\|\|chromosome:2512506-2512507(+):+0.217474[0.35042736,0.56790125] | | | | | | | +0.222912 to +0.222912 | +1.00222 to +1.00222 | -0.319654 to -0.319654 |
| terminator;1323 | DoxX family membrane protein;response regulator transcription factor SaeR;two-component system sensor histidine kinase SaeS | chromosome:2521829-2524018(+) | \|\|\|chromosome:2522044-2522045(-):+0.201357[0.5294118,0.7307692] | | | | | | | | | -0.231581 to +0.024495 | +0.42917 to +1.11923 | -0.446147 to +0.00433084 |
| terminator;1326 | N-acetylglucosamine-6-phosphate deacetylase;fructose-specific PTS transporter subunit EIIC;1-phosphofructokinase;DeoR/GlpR family DNA-binding transcription regulator | chromosome:2527836-2532970(-) | chromosome:2530966-2530967(-):-0.310924[0.42857143,0.11764706] | | | | | | chromosome:2530966-2530967(-):+0.20802[0.31578946,0.52380955] | | | -0.339885 to +0.111211 | -0.314761 to +0.432245 | -1.01328 to +0.352149 |
| terminator;1327 | Cys-tRNA(Pro) deacylase | chromosome:2533220-2533702(-) | chromosome:2533385-2533386(-):-0.242424[0.33333334,0.09090909] | | | chromosome:2533382-2533383(-):-0.206897[0.20689656,0] | | | | | | +0.248382 to +0.248382 | +0.339002 to +0.339002 | -0.248564 to -0.248564 |

| | | | | | | | | | | | | | | |
|---|---|---|---|---|---|---|---|---|---|---|---|---|---|---|
| terminator;133 | oligosaccharide flippase family protein;O-antigen ligase family protein;glycosyltransferase family 4 protein;sugar transferase;NAD-dependent epimerase/dehydratase family protein | chromosome:276532-282182(-) | chromosome:277851-277852(-):+0.318681[0.14285715,0.46153846] | | | | | | | | | +0.278343 to +1.95271 | -1.18682 to +0.0244273 | -0.690372 to -0.0576255 |
| terminator;1338 | hypothetical protein;YaiI/YqxD family protein;TIGR00730 family Rossman fold protein | chromosome:2548569-2550281(+) | | | | chromosome:2550027-2550028(+):-0.22488[0.90909094,0.68421054] | | | | | | -0.233967 to +0.341485 | +0.475038 to +1.26756 | -0.245376 to -0.0888593 |
| terminator;1360 | dihydroxyacetone kinase phosphoryl donor subunit DhaM;dihydroxyacetone kinase subunit DhaL;dihydroxyacetone kinase subunit DhaK | chromosome:2577040-2578989(-) | \|\|\|chromosome:2578829-2578830(+):+0.214912[0.41666666,0.6315789] | | | | | | | | | -0.56871 to -0.290474 | +0.604115 to +0.844232 | +0.954501 to +1.05484 |
| terminator;1371 | metal-dependent transcriptional regulator | chromosome:2595428-2596072(-) | chromosome:2595698-2595699(-):-0.203007[0.6315789,0.42857143]\|\|\|chromosome:2595690-2595691(+):+0.203209[0.09090909,0.29411766] | | | | | | | | | +0.298359 to +0.298359 | -0.603743 to -0.603743 | -0.463478 to -0.463478 |

| | | | | | | | | | | | | | | |
|---|---|---|---|---|---|---|---|---|---|---|---|---|---|---|
| terminator;1393 | DUF443 domain-containing protein;DUF443 family protein;hypothetical protein;DUF443 domain-containing protein;DUF443 family protein;DUF443 domain-containing protein;DUF443 family protein;T7SS effector LXG polymorphic toxin | chromosome:2640610-2646413(-) | | | | \|\|\|chromosome:2641447-2641448(+):+0.231579[0.36842105,0.6] | | | | | | -0.320961 to +0.535215 | -0.258104 to +0.43725 | -0.23758 to +0.0886232 |
| terminator;141 | L-lactate permease | chromosome:302628-304220(-) | | | | | | | chromosome:303720-303721(-):-0.25[0.55,0.3] | | | +0.219422 to +0.219422 | -0.0746335 to -0.0746335 | -0.6902 to -0.6902 |
| terminator;1414 | MSCRAMM family adhesin SdrE;MSCRAMM family adhesin SdrD;MSCRAMM family adhesin SdrC | chromosome:2676685-2687898(-) | \|\|\|chromosome:2685282-2685283(+):+0.209755[0.26785713,0.47761193] | | | \|\|\|chromosome:2685552-2685553(+):+0.302083[0.53125,0.8333333] | | | | | | -1.23091 to +0.387891 | -4.56531 to +0.250321 | -0.299623 to +1.01922 |
| terminator;1419 | branched-chain amino acid aminotransferase | chromosome:2692790-2693866(-) | | | | chromosome:2693296-2693297(-):+0.239011[0.15384616,0.39285713] | | | | | | -0.866204 to -0.866204 | +0.819974 to +0.819974 | -0.220183 to -0.220183 |

| | | | | | | | | | | | | | | |
|---|---|---|---|---|---|---|---|---|---|---|---|---|---|---|
| terminator;1426 | class I SAM-dependent methyltransferase;50S ribosomal protein L7/L12;50S ribosomal protein L10;50S ribosomal protein L1;50S ribosomal protein L11;transcription termination/antitermination protein NusG;preprotein translocase subunit SecE;50S ribosomal protein L33 | chromosome:2713705-2718116(-) | chromosome:2714718-2714719(-):+0.229946[0.5882353,0.8181818] | | | | | | | | | -0.415119 to +0.191552 | -1.68279 to -0.378908 | -0.342098 to +0.379181 |
| terminator;1430 | PIN/TRAM domain-containing protein;DNA repair protein RadA | chromosome:2725021-2727483(-) | chromosome:2727193-2727194(-):-0.286765[0.4117647,0.125] | | | | | | | | | -0.193952 to -0.0533731 | -0.062598 to +0.122994 | +0.308253 to +0.482629 |

| | | | | | | | | | | | | | | |
|---|---|---|---|---|---|---|---|---|---|---|---|---|---|---|
| terminator;1435 | 5S ribosomal RNA;23S ribosomal RNA;16S ribosomal RNA;5S ribosomal RNA;23S ribosomal RNA;tRNA-Ile;16S ribosomal RNA;tRNA-Ala;tRNA-Pro;tRNA-Arg;tRNA-Leu;tRNA-Gly;tRNA-Lys;tRNA-Thr;tRNA-Val;5S ribosomal RNA | chromosome:2737523-2748675(-) | 7988-2737989(-):-0.203661[0.7826087,0.57894737];chromosome:2738104-2738105(-):-0.475[0.875,0.4];chromosome:2738106-2738107(-):+0.287356[0.33333334,0.62068963];chromosome:2738540-2738541(-):+0.20979[0.15384616,0.36363637];chromosome:2740517-2740518(-):+0.209239[0.3125,0.5217391];chromosome:2743171-2743172(-):+0.25[0,0.25];chromosome:2743380-2743381(-):-0.222222[0.33333334,0.11111111];chromosome:2743522-2743523(-):+0.21448[0.13846155,0.3529412 | | | 8379-2738380(-):-0.206349[0.7619048,0.5555556];chromosome:2739305-2739306(-):+0.267943[0.36363637,0.6315789];chromosome:2740963-2740964(-):+0.229677[0.29032257,0.52];chromosome:2745991-2745992(-):-0.218301[0.5294118,0.31111112];chromosome:2747292-2747293(-):+0.215385[0.4,0.61538464];chromosome:2747664-2747665(-):+0.26087[0,0.26086956];chromosome:2748219-2748220(-):+0.209091[0.09090909,0.3]\|\|\|chromosome:2738354-2738355(+):+0.3 | chromosome:2741544-2741545(-):+0.270588[0.2,0.47058824] | | 7988-2737989(-):-0.203931[0.5675676,0.36363637];chromosome:2738084-2738085(-):+0.233333[0.1,0.33333334];chromosome:2742080-2742081(-):+0.2875[0.4,0.6875];chromosome:2744875-2744876(-):+0.302564[0.46666667,0.7692308];chromosome:2745991-2745992(-):+0.28882[0.2826087,0.5714286];chromosome:2746845-2746846(-):-0.208791[0.6730769,0.4642857]\|\|\|chromosome:2741074-2741075(+):+0.232143[0.64285713,0.875];chromosome:2745365-2745366(+):+0.2 | | | -1.1337 to +2.56788 | -2.48442 to +1.53133 | -0.475051 to +0.530879 |
| terminator;1437 | 2-amino-4-hydroxy-6-hydroxymethyldihydropteridine diphosphokinase;dihydroneopterin aldolase;dihydropteroate synthase | chromosome:2751259-2752878(-) | chromosome:2751789-2751790(-):+0.277778[0.11111111,0.3888889] | | | | | | | | | -0.336715 to -0.0859394 | -0.621542 to -0.427832 | -0.0508421 to +0.234096 |
| terminator;1440 | ATP-dependent zinc metalloprotease FtsH | chromosome:2755315-2757408(-) | | | | chromosome:2756994-2756995(-):-0.222222[0.4,0.17777778] | | | | | | -0.33719 to -0.33719 | +0.336275 to +0.336275 | +0.476378 to +0.476378 |

| | | | | | | | | | | | | | | |
|---|---|---|---|---|---|---|---|---|---|---|---|---|---|---|
| terminator;1442 | containing RNA-binding protein;cell division protein DivIC;RNA-binding S4 domain-containing protein;MazG nucleotide pyrophosphohydrolase domain-containing protein;polysaccharide biosynthesis protein;transcription-repair coupling factor;aminoacyl-tRNA hydrolase;50S ribosomal protein L25/general stress protein Ctc;ribose-phosphate diphosphokinase;bifunctional UDP-N- | chromosome:2759684-2771379(-) | | | | \|\|\|chromosome:2769182-2769183(+):-0.204319[0.41860464,0.21428572] | | | | | | -0.511782 to +1.18169 | -0.409198 to +0.647847 | -0.524228 to +0.176025 |
| terminator;1443 | septation regulator SpoVG;RidA family protein;pur operon repressor;4-(cytidine 5'-diphospho)-2-C-methyl-D-erythritol kinase | chromosome:2771569-2774026(-) | \|\|\|chromosome:2773728-2773729(+):-0.276973[0.4074074,0.13043478] | | | | | | | | | -0.224314 to +0.219519 | -0.211917 to +1.04101 | -0.491523 to -0.223435 |
| terminator;145 | oleate hydratase | chromosome:307701-309476(+) | chromosome:307931-307932(+):+0.238095[0.42857143,0.6666667] | | | chromosome:307931-307932(+):-0.235556[0.68,0.44444445] | | | | | | +1.06271 to +1.06271 | +1.56722 to +1.56722 | -0.0269741 to -0.0269741 |

| | | | | | | | | | | | | | | |
|---|---|---|---|---|---|---|---|---|---|---|---|---|---|---|
| terminator;1450 | 5S ribosomal RNA;23S ribosomal RNA;16S ribosomal RNA | chromosome:2786953-2791980(-) | chromosome:2787631-2787632(-):-0.237143[0.88,0.64285713];chromosome:2788202-2788203(-):+0.220833[0.46666667,0.6875];chromosome:2790904-2790905(-):+0.235294[0,0.23529412];chromosome:2791572-2791573(-):+0.202381[0.4642857,0.6666667]\|\|\|chromosome:2790566-2790567(+):+0.272727[0.36363637,0.6363636];chromosome:2791045-2791046(+):-0.240891[0.8947368,0.65384614] | | | chromosome:2787536-2787537(-):+0.246154[0.15384616,0.4];chromosome:2791040-2791041(-):+0.223684[0.5263158,0.75]\|\|\|chromosome:2790939-2790940(+):+0.238462[0.3,0.53846157];chromosome:2791626-2791627(+):+0.402027[0.125,0.527027] | | | 7536-2787537(-):+0.21371[0.625,0.83870965];chromosome:2787631-2787632(-):+0.248252[0.6363636,0.88461536];chromosome:2789132-2789133(-):-0.20904[0.5423729,0.33333334];chromosome:2791128-2791129(-):+0.254372[0.21621622,0.47058824]\|\|\|chromosome:2787878-2787879(+):+0.201238[0.0882353,0.28947368];chromosome:2787879-2787880(+):+0.264872[0.2244898,0.4893617];chromosome:2791046-2791047(+):-0.32971[0.41666666,0.08695652]; | | chromosome:2787657-2787658(-):-0.201839[0.76635516,0.5645161] | -0.647294 to +0.450594 | -2.85863 to +1.0081 | -0.516034 to +0.442043 |
| terminator;1456 | tRNA-Ser | chromosome:2801128-2801220(-) | chromosome:2801147-2801148(-):+0.258152[0.4375,0.6956522]\|\|\|chromosome:2801153-2801154(+):-0.217433[0.5277778,0.31034482] | | | | | | | | | +1.51937 to +1.51937 | +1.54234 to +1.54234 | -0.640062 to -0.640062 |
| terminator;1457 | glutamate synthase subunit beta;glutamate synthase large subunit | chromosome:2801489-2807469(-) | chromosome:2807322-2807323(-):+0.2125[0.1875,0.4]\|\|\|chromosome:2806777-2806778(+):-0.244939[0.55263156,0.30769232] | | | \|\|\|chromosome:2802511-2802512(+):+0.218474[0.12195122,0.34042552] | | | chromosome:2807316-2807317(-):-0.208333[0.20833333,0];chromosome:2807322-2807323(-):+0.20761[0.17948718,0.38709676] | | | -1.35375 to -0.351208 | +2.34188 to +3.06566 | -0.967865 to -0.887405 |
| terminator;1462 | autolysin/adhesin Aaa | chromosome:2812188-2813192(-) | \|\|\|chromosome:2812726-2812727(+):+0.219002[0.5217391,0.7407407] | | | | | | | | | +0.0811539 to +0.0811539 | -1.65117 to -1.65117 | +1.05374 to +1.05374 |

| | | | | | | | | | | | | | | |
|---|---|---|---|---|---|---|---|---|---|---|---|---|---|---|
| terminator;1470 | DUF2309 domain-containing protein;NADH dehydrogenase subunit 5 | chromosome:2823087-2827289(-) | | | | chromosome:2823579-2823580(-):+0.264706[0.23529412,0.5] | | | chromosome:2823579-2823580(-):+0.35[0.15,0.5] | | | -0.0084627 to +0.115784 | +1.83142 to +1.94358 | -0.0992706 to +0.178382 |
| terminator;1490 | superantigen-like protein SSL4 | chromosome:2854128-2855054(-) | \|\|\|chromosome:2854964-2854965(+):+0.342995[0.4347826,0.7777778] | | | \|\|\|chromosome:2854964-2854965(+):+0.20202[0.57575756,0.7777778] | | | | | | +0.822472 to +0.822472 | +2.82105e-05 to +2.82105e-05 | -0.23837 to -0.23837 |
| terminator;1492 | superantigen-like protein SSL3 | chromosome:2855419-2856477(-) | \|\|\|chromosome:2856107-2856108(+):-0.228758[0.61764705,0.3888889] | | | \|\|\|chromosome:2856107-2856108(+):+0.229798[0.4090909,0.6388889] | | | | | | +0.791979 to +0.791979 | +1.07449 to +1.07449 | -0.831418 to -0.831418 |
| terminator;1500 | glutamine-hydrolyzing GMP synthase;IMP dehydrogenase;xanthine permease PbuX;xanthine phosphoribosyltransferase | chromosome:2863478-2868394(-) | chromosome:2865189-2865190(-):+0.272941[0.08,0.3529412];chromosome:2866653-2866654(-):+0.207046[0.36158192,0.5686275] | | | | | | | | | -0.198487 to +0.329876 | -0.658001 to -0.187529 | +0.541665 to +0.718691 |
| terminator;1505 | L-cystine transporter | chromosome:2872170-2873558(+) | chromosome:2873325-2873326(+):+0.246667[0.23333333,0.48] | | | | | | | | | -0.153033 to -0.153033 | -1.67181 to -1.67181 | -0.22172 to -0.22172 |
| terminator;1527 | aminoglycoside O-phosphotransferase APH(3')-IIIa | pUSA300-1:22524-23318(-) | \|\|\|pUSA300-1:22744-22745(+):+0.233333[0.3,0.53333336] | | | pUSA300-1:23030-23031(-):+0.269231[0.1923077,0.46153846] | | | | | | +0.0199552 to +0.0199552 | +0.5428 to +0.5428 | +0.0504578 to +0.0504578 |
| terminator;168 | ABC transporter ATP-binding protein;ATP-binding cassette domain-containing protein;ABC transporter permease;ABC transporter permease;ABC transporter substrate-binding protein;class I SAM-dependent methyltransferase | chromosome:346253-351583(-) | | | | | | | chromosome:349048-349049(-):+0.43089[0.41121495,0.84210527] | | | +0.728541 to +1.12902 | -0.812046 to +0.255066 | -0.614111 to +0.160026 |

| | | | | | | | | | | | | | | |
|---|---|---|---|---|---|---|---|---|---|---|---|---|---|---|
| terminator;173 | arginine repressor;arginine deiminase;arginine-ornithine antiporter;Crp/Fnr family transcriptional regulator;ornithine carbamoyltransferase;carbamate kinase | chromosome:356225-362398(+) | chromosome:359221-359222(+):-0.204872[0.49333334,0.28846154] | | | chromosome:361936-361937(+):+0.247412[0.071428575,0.3188406] | | | | | | -0.0319636 to +1.08327 | -0.307924 to +1.09104 | -0.126835 to +2.58187 |
| terminator;227 | anion permease | chromosome:449847-451265(-) | | | | \|\|\|chromosome:450064-450065(+):-0.268092[0.53125,0.2631579];chromosome:450070-450071(+):+0.253232[0.1521739,0.4054054] | | | | | | -0.283639 to -0.283639 | -1.72314 to -1.72314 | -0.488766 to -0.488766 |
| terminator;234 | S-adenosyl-l-methionine hydroxide adenosyltransferase family protein;ECF-type riboflavin transporter substrate-binding protein;ABC transporter ATP-binding protein;energy-coupling factor transporter transmembrane protein EcfT;GNAT family protein | chromosome:457353-461923(+) | | | | chromosome:457566-457567(+):-0.204678[0.31578946,0.11111111] | | | | | | -1.93963 to -0.659642 | +1.01762 to +1.91343 | -1.55093 to -1.19067 |

<table>
<tr><td>terminator;239</td><td>polysaccharide intercellular adhesin biosynthesis/export protein IcaC;intercellular adhesin biosynthesis polysaccharide N-deacetylase;intracellular adhesion protein IcaD;poly-beta-1%2C6 N-acetyl-D-glucosamine synthase IcaA</td><td>chromosome:473977-477392(-)</td><td></td><td></td><td></td><td></td><td></td><td></td><td>|||chromosome:476169-476170(+):-0.308271[0.7368421,0.42857143]</td><td></td><td></td><td>-0.508855 to +0.89835</td><td>-0.195611 to +0.940414</td><td>-0.569595 to +0.486224</td></tr>
<tr><td>terminator;24</td><td>hypothetical protein</td><td>chromosome:35335-35631(-)</td><td></td><td></td><td></td><td>|||chromosome:35409-35410(+):+0.303571[0.525,0.82857144]</td><td></td><td></td><td>|||chromosome:35464-35465(+):+0.216247[0.1521739,0.36842105]</td><td></td><td></td><td>+0.64641 to +0.64641</td><td>-0.948721 to -0.948721</td><td>-1.66085 to -1.66085</td></tr>
<tr><td>terminator;247</td><td>serine-rich repeat glycoprotein adhesin SasA</td><td>chromosome:485821-492636(+)</td><td>chromosome:486991-486992(+):-0.297203[0.61538464,0.3181818]</td><td></td><td></td><td></td><td></td><td></td><td></td><td></td><td></td><td>-0.151388 to -0.151388</td><td>+0.499412 to +0.499412</td><td>+1.03656 to +1.03656</td></tr>
<tr><td>terminator;252</td><td>YhgE/Pip family protein</td><td>chromosome:508507-511488(+)</td><td>chromosome:509629-509630(+):+0.254386[0.078947365,0.33333334]</td><td></td><td></td><td></td><td></td><td></td><td></td><td></td><td></td><td>+0.270579 to +0.270579</td><td>-0.271865 to -0.271865</td><td>-0.0659837 to -0.0659837</td></tr>
<tr><td>terminator;253</td><td>mannose-6-phosphate isomerase%2C class I;fructose-specific PTS transporter subunit EIIC;transcription antiterminator</td><td>chromosome:511600-516455(-)</td><td>chromosome:513224-513225(-):-0.202381[0.2857143,0.083333336]</td><td></td><td></td><td></td><td></td><td></td><td></td><td></td><td></td><td>-0.389623 to +0.231002</td><td>-0.67942 to +0.34131</td><td>+0.0748817 to +0.431894</td></tr>
<tr><td>terminator;256</td><td>zinc metalloproteinase aureolysin</td><td>chromosome:518362-519891(+)</td><td>|||chromosome:519561-519562(-):+0.216667[0.33333334,0.55]</td><td></td><td></td><td></td><td></td><td></td><td></td><td></td><td></td><td>+0.358435 to +0.358435</td><td>-2.22223 to -2.22223</td><td>-1.09297 to -1.09297</td></tr>
<tr><td>terminator;259</td><td>MSCRAMM family adhesin clumping factor ClfB</td><td>chromosome:526964-529663(+)</td><td>chromosome:528665-528666(+):+0.221839[0.43333334,0.6551724]|||chromosome:529332-529333(-):+0.205128[0.7948718,1]</td><td></td><td></td><td>|||chromosome:528637-528638(-):+0.266667[0.33333334,0.6]</td><td></td><td></td><td>|||chromosome:528615-528616(-):+0.215311[0.42105263,0.6363636];chromosome:528643-528644(-):+0.331933[0.38235295,0.71428573]</td><td></td><td></td><td>-0.59428 to -0.59428</td><td>-1.04366 to -1.04366</td><td>+0.396314 to +0.396314</td></tr>
</table>

| | | | | | | | | | | | | | | |
|---|---|---|---|---|---|---|---|---|---|---|---|---|---|---|
| terminator;26 | NADPH-dependent FMN reductase;LLM class flavin-dependent oxidoreductase;VOC family protein | chromosome:36784-39365(-) | | | | chromosome:37847-37848(-):+0.302083[0.16666667,0.46875] | | | | | | -0.523412 to -0.289127 | -0.259554 to -0.0336714 | +0.44922 to +0.600254 |
| terminator;269 | assimilatory sulfite reductase (NADPH) flavoprotein subunit;NAD(P)-binding protein | chromosome:540110-542658(+) | | | | chromosome:542114-542115(+):+0.228889[0.16,0.3888889] | | | | | | +0.210059 to +0.272889 | +1.8791 to +2.24773 | +0.184641 to +0.399902 |
| terminator;291 | oxidoreductase | chromosome:567353-568213(-) | | | | chromosome:568187-568188(-):+0.202797[0.18181819,0.3846154] | | | | | | -0.193633 to -0.193633 | +0.352174 to +0.352174 | -0.556543 to -0.556543 |
| terminator;299 | fructosamine kinase family protein | chromosome:576368-577234(+) | \|\|\|chromosome:576647-576648(-):+0.301314[0.046511628,0.3478261] | | | | | chromosome:576580-576581(+):-0.212403[0.27906978,0.06666667] | | | | +0.158709 to +0.158709 | +0.269303 to +0.269303 | -0.13211 to -0.13211 |
| terminator;30 | transcription antiterminator;PTS sugar transporter subunit IIA;PTS sugar transporter subunit IIB;PTS ascorbate transporter subunit IIC | chromosome:43751-47801(+) | \|\|\|chromosome:45595-45596(-):-0.22[0.3,0.08] | | chromosome:45603-45604(+):-0.29064[0.42857143,0.13793103];chromosome:46139-46140(+):-0.212698[0.6571429,0.44444445];chromosome:46146-46147(+):-0.240602[0.5263158,0.2857143] | | | chromosome:46187-46188(+):+0.20979[0.15384616,0.36363637]\|\|\|chromosome:45654-45655(-):+0.21657[0.08955224,0.30612245] | | | | -0.150827 to +0.972412 | +0.285616 to +1.14763 | +0.563854 to +1.05881 |
| terminator;316 | aminotransferase class I/II-fold pyridoxal phosphate-dependent enzyme;D-lactate dehydrogenase | chromosome:602291-604466(+) | \|\|\|chromosome:602404-602405(-):+0.315556[0.24,0.5555556] | | | | | | | | | -0.382514 to -0.305956 | +0.41423 to +0.839202 | -1.03998 to -0.674224 |

| | | | | | | | | | | | | | | |
|---|---|---|---|---|---|---|---|---|---|---|---|---|---|---|
| terminator;341 | PTS sugar transporter subunit IIC;L-serine ammonia-lyase%2C iron-sulfur-dependent subunit beta;L-serine ammonia-lyase%2C iron-sulfur-dependent%2C subunit alpha;hypothetical protein;hypothetical protein | chromosome:635958-638946(+) | chromosome:638233-638234(+):-0.300926[0.7083333,0.4074074] | | | | | | | | | -0.176469 to +3.03738 | -0.821197 to +0.153389 | -1.93667 to +0.673111 |
| terminator;362 | hypothetical protein;fibronectin-binding protein FnbA | chromosome:668117-671258(+) | \|\|\|chromosome:670955-670956(-):-0.264444[0.32,0.055555556];chromosome:670969-670970(-):+0.233333[0.5,0.73333335] | | | \|\|\|chromosome:670870-670871(-):+0.20903[0.26923078,0.47826087] | | | | | | -0.415707 to +2.72622 | -0.891944 to +2.28516 | +0.52543 to +1.73926 |
| terminator;363 | fibronectin-binding protein FnbB | chromosome:671940-674762(+) | chromosome:674388-674389(+):-0.285362[0.390625,0.10526316] | | | \|\|\|chromosome:674453-674454(-):-0.216783[0.4090909,0.1923077] | | | | | | -0.288636 to -0.288636 | -0.467591 to -0.467591 | +1.14657 to +1.14657 |
| terminator;385 | staphylopine-dependent metal ABC transporter substrate-binding protein CntA;nickel/cobalt ABC transporter permease;nickel/cobalt ABC transporter permease;ABC transporter ATP-binding protein;ABC transporter ATP-binding protein;MFS transporter | chromosome:706650-712821(+) | \|\|\|chromosome:711222-711223(-):-0.22686[0.37579617,0.14893617] | | | | | | | | | +0.177084 to +1.06419 | -1.42715 to +0.0912527 | -0.119764 to +0.528331 |
| terminator;397 | hypothetical protein | chromosome:726426-726545(+) | | | | | | | chromosome:726430-726431(+):+0.246753[0.18181819,0.42857143] | | | +0.751325 to +0.751325 | +0.932991 to +0.932991 | -0.354686 to -0.354686 |

| | | | | | | | | | | | | | | |
|---|---|---|---|---|---|---|---|---|---|---|---|---|---|---|
| terminator;399 | MFS transporter | chromosome:730677-732077(+) | chromosome:731679-731680(+):+0.241597[0.29411766,0.53571427];chromosome:731856-731857(+):+0.214973[0.09090909,0.30588236] | | | | | | | | | -0.575974 to -0.575974 | +0.776895 to +0.776895 | -0.529676 to -0.529676 |
| terminator;407 | C39 family peptidase | chromosome:742980-743636(-) | | | | chromosome:743481-743482(-):-0.216346[0.5625,0.34615386] | | | | | | -0.547985 to -0.547985 | -0.393463 to -0.393463 | +0.742507 to +0.742507 |
| terminator;42 | PTS sugar transporter subunit IIC | chromosome:68961-69950(-) | | | | | | | \|\|\|chromosome:69775-69776(+):-0.218254[0.8055556,0.5873016] | | | -0.564456 to -0.564456 | -0.798991 to -0.798991 | -0.275979 to -0.275979 |
| terminator;433 | nitrate reductase subunit alpha;nitrate reductase subunit beta;nitrate reductase molybdenum cofactor assembly chaperone;respiratory nitrate reductase subunit gamma;nitrate respiration regulation accessory nitrate sensor NreA;nitrate respiration regulation sensor histidine kinase NreB;nitrate respiration regulation response regulator NreC | chromosome:783520-792202(+) | | | | chromosome:785195-785196(+):+0.485714[0.21428572,0.7] | | | | | | -0.452456 to +0.0187975 | -0.602408 to +0.514277 | -2.15677 to -1.60674 |
| terminator;448 | hypothetical protein;NAD(P)/FAD-dependent oxidoreductase;GNAT family N-acetyltransferase | chromosome:810050-811648(+) | | | | | | | chromosome:810210-810211(+):+0.209549[0.4827586,0.6923077] | | | -0.255202 to +1.78506 | -0.528499 to +1.85744 | +0.310394 to +1.71926 |

| | | | | | | | | | | | | | | |
|---|---|---|---|---|---|---|---|---|---|---|---|---|---|---|
| terminator;454 | malate dehydrogenase (quinone) | chromosome:818469-819947(+) | chromosome:819357-819358(+):-0.210084[0.3529412,0.14285715] | | | | | | | | | -0.411586 to -0.411586 | -0.83311 to -0.83311 | -0.231215 to -0.231215 |
| terminator;473 | ribose 5-phosphate isomerase A | chromosome:847657-848343(+) | | | | | | | chromosome:847965-847966(+):-0.219048[0.26666668,0.04761905] | | | -0.20824 to -0.20824 | -0.994181 to -0.994181 | +0.181885 to +0.181885 |
| terminator;485 | bile acid:sodium symporter family protein;HAD family hydrolase;hypothetical protein | chromosome:864910-866739(+) | chromosome:865227-865228(+):+0.212821[0.15384616,0.36666667] | | | | | | | | | -0.691914 to -0.295892 | +0.201414 to +2.63665 | -0.99713 to -0.0808777 |
| terminator;489 | hypothetical protein | chromosome:870736-871065(-) | | | | \|\|\|chromosome:870826-870827(+):-0.329114[0.32911393,0] | | | | | | +0.531642 to +0.531642 | +0.757656 to +0.757656 | -0.669755 to -0.669755 |
| terminator;496 | FAD-dependent monooxygenase | chromosome:881051-882175(+) | \|\|\|chromosome:881920-881921(-):-0.266129[0.75,0.48387095] | | | | | | | | | -0.366545 to -0.366545 | -0.916376 to -0.916376 | +0.183616 to +0.183616 |
| terminator;530 | GRP family sugar transporter | chromosome:926010-926873(-) | | | | \|\|\|chromosome:926788-926789(+):+0.224532[0.15384616,0.3783784] | | | | | | -0.311534 to -0.311534 | -1.28413 to -1.28413 | +0.965745 to +0.965745 |
| terminator;532 | hypothetical protein;DNA topoisomerase III | chromosome:928007-930263(+) | chromosome:928697-928698(+):+0.260526[0.039473683,0.3] | | | | | | | | | -0.446791 to +0.913634 | -0.522908 to -0.486848 | -0.294256 to -0.0548332 |

| | | | | | | | | | | | | | | |
|---|---|---|---|---|---|---|---|---|---|---|---|---|---|---|
| terminator;535 | protein S5;50S ribosomal protein L30;50S ribosomal protein L15;preprotein translocase subunit SecY;adenylate kinase;translation initiation factor IF-1;50S ribosomal protein L36;30S ribosomal protein S13;30S ribosomal protein S11;DNA-directed RNA polymerase subunit alpha;50S ribosomal protein L17;30S ribosomal protein S10;50S ribosomal protein L3;50S ribosomal protein L4;50S ribosomal protein L23;50S ribosomal protein | chromosome:932648-946563(+) | chromosome:938072-938073(+):-0.247863[0.6923077,0.44444445];chromosome:946523-946524(+):+0.262599[0.46153846,0.7241379] | | | | | | | | | -0.699887 to -0.176404 | -1.65079 to -0.662525 | -0.337348 to +0.839299 |
| terminator;554 | BCCT family transporter | chromosome:981591-983153(+) | | | | | chromosome:982137-982138(+):+0.207072[0.22857143,0.43564355] | | | | | -0.175042 to -0.175042 | +2.52132 to +2.52132 | -0.266315 to -0.266315 |

| | | | | | | | | | | | | | | |
|---|---|---|---|---|---|---|---|---|---|---|---|---|---|---|
| terminator;568 | 16S ribosomal RNA;23S ribosomal RNA;5S ribosomal RNA;tRNA-Asn;tRNA-Glu;tRNA-Val;tRNA-Tyr;tRNA-Gln;tRNA-Lys | chromosome:1006427-1011906(+) | 9653-1009654(+):+0.32266[0.03448276,0.35714287];chromosome:1010471-1010472(+):-0.2125[0.3125,0.1];chromosome:1010829-1010830(+):-0.270909[0.59090906,0.32];chromosome:1010925-1010926(+):+0.205742[0.6363636,0.84210527]\|\|\|chromosome:1006820-1006821(-):-0.256757[0.7567568,0.5];chromosome:1007467-1007468(-):+0.267692[0.04,0.30769232];chromosome:1007839-1007840(-):+0.326316[0.47368422,0.8];chromosome:1008726-1008727(-):- | | | \|\|\|chromosome:1007619-1007620(-):+0.270175[0.06315789,0.33333334];chromosome:1009206-1009207(-):-0.222222[0.33333334,0.11111111];chromosome:1010147-1010148(-):+0.310247[0.20588236,0.516129] | | | 7052-1007053(+):-0.211174[0.24242425,0.03125];chromosome:1010146-1010147(+):+0.211765[0.2,0.4117647];chromosome:1011736-1011737(+):+0.25317[0.23958333,0.49275362]\|\|\|chromosome:1007320-1007321(-):+0.207143[0.25,0.45714286];chromosome:1008820-1008821(-):-0.20303[0.36666667,0.16363636];chromosome:1010147-1010148(-):-0.24[0.74,0.5];chromosome:1010365-1010366(-):+0.329412[0.47058824,0.8];chromosome:1010465-1010466(- | chromosome:1007369-1007370(+):-0.35[0.85,0.5] | | -1.28537 to +0.450594 | -2.85863 to +1.03177 | -0.666824 to +0.176056 |
| terminator;588 | M20 family metallopeptidase;ATP-grasp domain-containing protein | chromosome:1052368-1054745(+) | | | | chromosome:1054086-1054087(+):+0.26455[0.14285715,0.4074074] | | | | | | -1.14754 to -1.01463 | +0.963309 to +1.04577 | -0.581932 to -0.460033 |
| terminator;590 | type II pantothenate kinase | chromosome:1055945-1056748(-) | | | | | | \|\|\|chromosome:1056247-1056248(+):+0.259259[0.074074075,0.33333334] | | | | +0.123692 to +0.123692 | -1.16029 to -1.16029 | -0.0629448 to -0.0629448 |
| terminator;603 | UDP-N-acetylglucosamine 1-carboxyvinyltransferase;3-hydroxyacyl-ACP dehydratase FabZ | chromosome:1084146-1085885(+) | chromosome:1085601-1085602(+):-0.274155[0.3611111,0.08695652] | | | | | chromosome:1084810-1084811(+):+0.210053[0.20930232,0.41935483] | | | | -0.280448 to -0.268536 | -0.608396 to -0.379973 | +0.14105 to +0.233642 |

| | | | | | | | | | | | | | | |
|---|---|---|---|---|---|---|---|---|---|---|---|---|---|---|
| terminator;606 | thiaminase II;bifunctional hydroxymethylpyrimidine kinase/phosphomethylpyrimidine kinase;hydroxyethylthiazole kinase;thiamine phosphate synthase;membrane protein insertase YidC | chromosome:1088443-1092331(+) | chromosome:1090778-1090779(+):+0.25[0.33333334,0.5833333] | | | \|\|\|chromosome:1091153-1091154(-):+0.35101[0.19444445,0.54545456] | | | | | | -0.315377 to +0.577698 | -1.0083 to +1.20165 | +0.21241 to +0.875506 |
| terminator;619 | tRNA-Leu;tRNA-Gly;16S ribosomal RNA;23S ribosomal RNA;5S ribosomal RNA | chromosome:1122440-1127702(+) | chromosome:1124281-1124282(+):-0.243961[0.5217391,0.2777778];chromosome:1126455-1126456(+):+0.234848[0.5833333,0.8181818];chromosome:1127204-1127205(+):+0.214286[0.2857143,0.5]\|\|\|chromosome:1125516-1125517(-):+0.233333[0.1,0.33333334];chromosome:1125986-1125987(-):-0.29[0.65,0.36];chromosome:1126457-1126458(-):-0.232143[0.60714287,0.375] | | | chromosome:1125994-1125995(+):+0.26046[0.0862069,0.34666666];chromosome:1127117-1127118(+):-0.274131[0.7027027,0.42857143];chromosome:1127138-1127139(+):+0.235993[0.29032257,0.5263158]\|\|\|chromosome:1123629-1123630(-):+0.3[0.5,0.8];chromosome:1124012-1124013(-):+0.211348[0.25531915,0.46666667];chromosome:1126675-1126676(-):-0.305556[0.5555556,0.25];chromosome:1126869-1126870(-):-0.256944[0.6944444,0.4375] | | | chromosome:1122493-1122494(+):+0.202857[0.15714286,0.36];chromosome:1123675-1123676(+):-0.341176[0.9,0.5588235];chromosome:1124211-1124212(+):-0.280702[0.28070176,0]\|\|\|chromosome:1122450-1122451(-):+0.261905[0.6666667,0.9285714];chromosome:1125986-1125987(-):-0.274945[0.36585367,0.09090909];chromosome:1126775-1126776(-):+0.246454[0.17021276,0.41666666] | | | -0.949407 to +3.9244 | -2.05999 to +1.3211 | -2.56639 to -0.0290437 |

| | | | | | | | | | | | | | | |
|---|---|---|---|---|---|---|---|---|---|---|---|---|---|---|
| terminator;620 | threonine ammonia-lyase IlvA;3-isopropylmalate dehydratase small subunit;3-isopropylmalate dehydratase large subunit;3-isopropylmalate dehydrogenase;2-isopropylmalate synthase;ketol-acid reductoisomerase;ACT domain-containing protein;biosynthetic-type acetolactate synthase large subunit;dihydroxy-acid dehydratase | chromosome:1127912-1138640(-) | chromosome:1129915-1129916(-):+0.279245[0.3207547,0.6] | | | | | | | | | -1.84435 to +0.494887 | +0.863103 to +1.99774 | -2.4026 to -1.16457 |
| terminator;622 | tRNA (adenosine(37)-N6)-threonylcarbamoyltransferase complex ATPase subunit type 1 TsaE;tRNA (adenosine(37)-N6)-threonylcarbamoyltransferase complex dimerization subunit type 1 TsaB;ribosomal protein S18-alanine N-acetyltransferase;tRNA (adenosine(37)-N6)-threonylcarbamoyltransferase complex transferase subunit TsaD | chromosome:1139151-1141710(+) | \|\|\|chromosome:1140885-1140886(-):-0.233333[0.5,0.26666668] | | | | | | | | | -0.142482 to +0.460071 | -1.17944 to -0.224708 | +0.0241902 to +0.580981 |

<table>
<tr><td>terminator;627</td><td>YeeE/YedE family protein;sulfurtransferase TusA family protein</td><td>chromosome:1146903-1148266(+)</td><td>chromosome:1147794-1147795(+):-0.335968[0.6086956,0.27272728]</td><td></td><td></td><td></td><td></td><td></td><td></td><td></td><td></td><td>-1.23569 to -1.15086</td><td>-0.785827 to -0.724355</td><td>-0.433239 to -0.305249</td></tr>
<tr><td>terminator;629</td><td>LacI family DNA-binding transcriptional regulator;sucrose-6-phosphate hydrolase</td><td>chromosome:1149907-1152490(+)</td><td>chromosome:1150714-1150715(+):-0.209302[0.20930232,0]</td><td></td><td></td><td></td><td></td><td></td><td></td><td></td><td></td><td>-0.772636 to -0.359056</td><td>-0.461882 to +0.388043</td><td>+0.202651 to +0.291748</td></tr>
<tr><td>terminator;635</td><td>SdrH family protein</td><td>chromosome:1159508-1160761(+)</td><td>chromosome:1160090-1160091(+):+0.227575[0.21428572,0.44186047]</td><td></td><td></td><td></td><td></td><td></td><td></td><td></td><td></td><td>+0.12742 to +0.12742</td><td>+0.670063 to +0.670063</td><td>-0.672592 to -0.672592</td></tr>
<tr><td>terminator;640</td><td>TrkH family potassium uptake protein</td><td>chromosome:1167337-1168644(-)</td><td>chromosome:1167757-1167758(-):-0.363821[0.5833333,0.2195122]</td><td></td><td></td><td></td><td></td><td></td><td></td><td></td><td></td><td>+0.0660154 to +0.0660154</td><td>-1.07504 to -1.07504</td><td>-0.594268 to -0.594268</td></tr>
<tr><td>terminator;649</td><td>helix-turn-helix transcriptional regulator;MW1434 family type I TA system toxin</td><td>chromosome:1180412-1180906(+)</td><td>chromosome:1180842-1180843(+):-0.240909[0.75,0.5090909];chromosome:1180845-1180846(+):+0.219958[0.13636364,0.35632184]|||chromosome:1180853-1180854(-):+0.221154[0.15384616,0.375]</td><td></td><td></td><td></td><td></td><td></td><td>|||chromosome:1180853-1180854(-):-0.281513[0.3529412,0.071428575]</td><td></td><td></td><td>+4.41839 to +4.49029</td><td>-0.740678 to -0.646053</td><td>-0.422134 to -0.369223</td></tr>
</table>

| | | | | | | | | | | | | | | |
|---|---|---|---|---|---|---|---|---|---|---|---|---|---|---|
| terminator;655 | hydrolase;single-stranded DNA-binding protein;DnaD domain protein;hypothetical protein;RusA family crossover junction endodeoxyribonuclease;SA1788 family PVL leukocidin-associated protein;phi PVL orf 51-like protein;acetyltransferase;DUF1024 family protein;hypothetical protein;dUTP diphosphatase;hypothetical protein;DUF1381 domain-containing protein;hypothetical protein;transcriptional | chromosome:1186865-1193166(+) | chromosome:1189167-1189168(+):-0.232278[0.4117647,0.17948718];chromosome:1192484-1192485(+):+0.291667[0.375,0.6666667] | | | chromosome:1192906-1192907(+):-0.248054[0.6097561,0.3617021] | | | | | | -0.212817 to +3.97564 | -0.506964 to +0.11136 | -0.725311 to +0.864757 |

| | | | | | | | | | | | | | | |
|---|---|---|---|---|---|---|---|---|---|---|---|---|---|---|
| terminator;657 | protein;terminase large subunit;phage portal protein;head maturation protease%2C ClpP-related;phage major capsid protein;hypothetical protein;phage head-tail adapter protein;head-tail adaptor protein;HK97 gp10 family phage protein;hypothetical protein;major tail protein;Ig-like domain-containing protein;hypothetical protein;hypothetical protein;phage tail tape measure protein;phage tail domain- | chromosome:1193827-1212848(+) | | | | chromosome:1205061-1205062(+):+0.237143[0.12,0.35714287] | | chromosome:1203016-1203017(+):-0.213636[0.23636363,0.022727273] | chromosome:1205681-1205682(+):+0.218402[0.13043478,0.3488372];chromosome:1207498-1207499(+):-0.209596[0.2777778,0.06818182] | | | +3.2242 to +4.12005 | -0.857005 to +0.263013 | -1.20077 to -0.0132812 |
| terminator;70 | 6-phospho-beta-glucosidase;glucose PTS transporter subunit IIA | chromosome:117414-119657(-) | | | | chromosome:118566-118567(-):+0.252174[0.4,0.65217394] | | | | | | +0.0103718 to +0.0717981 | +0.439825 to +1.06657 | -0.122156 to +0.0417711 |

| | | | | | | | | | | | | | | |
|---|---|---|---|---|---|---|---|---|---|---|---|---|---|---|
| terminator;721 | RNA;tRNA-Ile;tRNA-Ala;23S ribosomal RNA;23S ribosomal RNA;tRNA-Val;tRNA-Thr;tRNA-Lys;tRNA-Leu;tRNA-Gly;tRNA-Leu;tRNA-Arg;tRNA-Pro;tRNA-Ala;tRNA-Met;tRNA-Ile;tRNA-Ser;tRNA-Asp;tRNA-Ser;tRNA-Met;tRNA-Asp;tRNA-Phe;tRNA-Thr;tRNA-Tyr;tRNA-Trp;tRNA-His;tRNA-Gln;tRNA-Cys;tRNA-Gly;tRNA- | chromosome:1300732-1308312(+) | 1585-1301586(+):-0.210407[0.44117647,0.23076923];chromosome:1301794-1301795(+):-0.285714[0.42857143,0.14285715];chromosome:1301811-1301812(+):-0.203636[0.36363637,0.16];chromosome:1302276-1302277(+):-0.215517[0.46551725,0.25];chromosome:1302436-1302437(+):-0.207143[0.6,0.39285713];chromosome:1303686-1303687(+):+0.214286[0,0.21428572];chromosome:1305304-1305305(+):-0.212644[0.37931034,0.16666667 | chromosome:1303971-1303972(+):-0.212454[0.30769232,0.0952381];chromosome:1304849-1304850(+):-0.223077[0.3,0.07692308] | | 2435-1302436(+):-0.222222[0.22222222,0];chromosome:1302516-1302517(+):-0.244186[0.5,0.25581396];chromosome:1304231-1304232(+):+0.280702[0.05263158,0.33333334];chromosome:1305010-1305011(+):-0.337607[0.7222222,0.3846154];chromosome:1305323-1305324(+):-0.269231[0.5,0.23076923];chromosome:1306031-1306032(+):+0.537838[0.16216215,0.7];chromosome:1306741-1306742(+):-0.336364[0.7,0.36363637];chromosome:1307150-1307151(+):- | | | 2516-1302517(+):+0.222591[0.3488372,0.5714286];chromosome:1304617-1304618(+):+0.268421[0.6315789,0.9];chromosome:1304947-1304948(+):+0.411765[0,0.4117647];chromosome:1305010-1305011(+):-0.33913[0.73913044,0.4];chromosome:1305045-1305046(+):+0.253968[0.04761905,0.3015873];chromosome:1305318-1305319(+):+0.201299[0.72727275,0.9285714];chromosome:1305395-1305396(+):-0.209694[0.6712329,0.46153846]; | | | -1.39863 to +0.734174 | +0.130338 to +1.55063 | +0.254498 to +0.519571 |
| terminator;724 | ABC transporter permease subunit;amino acid ABC transporter ATP-binding protein | chromosome:1311578-1313750(+) | | | | \|\|\|chromosome:1313614-1313615(-):-0.201677[0.7111111,0.509434] | | | | | | -0.639222 to -0.639173 | -1.85929 to -1.49967 | -0.549562 to -0.256317 |
| terminator;730 | GAF domain-containing sensor histidine kinase;response regulator | chromosome:1320638-1322395(+) | chromosome:1321916-1321917(+):+0.200466[0.030303031,0.23076923]\|\|\|chromosome:1322167-1322168(-):+0.299663[0.18181819,0.4814815] | | chromosome:1321818-1321819(+):-0.213665[0.25714287,0.04347826] | | | chromosome:1321674-1321675(+):+0.225732[0.045454547,0.27118644];chromosome:1321818-1321819(+):+0.218102[0.27027026,0.4883721];chromosome:1322155-1322156(+):-0.20595[0.68421054,0.47826087] | | | | +0.350477 to +0.654943 | +0.142033 to +0.192729 | -0.291436 to -0.137615 |
| terminator;735 | peptidylprolyl isomerase | chromosome:1330884-1331846(-) | \|\|\|chromosome:1330897-1330898(+):+0.321839[0.3448276,0.6666667] | | | | | | | | | +0.245533 to +0.245533 | +3.04373 to +3.04373 | -0.205272 to -0.205272 |

| | | | | | | | | | | | | | | |
|---|---|---|---|---|---|---|---|---|---|---|---|---|---|---|
| terminator;743 | tRNA-Met;tRNA-Asp;tRNA-Phe;tRNA-His;tRNA-Gly;tRNA-Asn;tRNA-Glu;tRNA-Ser | chromosome:1342041-1342783(+) | chromosome:1342573-1342574(+):-0.238095[0.23809524,0] | | | | | | | | | +2.37461 to +2.51657 | -1.28193 to -0.972375 | -1.21378 to -1.206 |
| terminator;749 | DUF4888 domain-containing protein | chromosome:1358203-1358769(-) | | | | \|\|\|chromosome:1358283-1358284(+):+0.287477[0.31858408,0.6060606] | | | | | | +2.32194 to +2.32194 | -0.147971 to -0.147971 | -0.192278 to -0.192278 |
| terminator;762 | DUF4352 domain-containing protein;excalibur calcium-binding domain-containing protein | chromosome:1378702-1380428(-) | | chromosome:1379568-1379569(-):-0.227273[0.5,0.27272728] | | | | | | | | -0.804507 to -0.79915 | -0.128065 to -0.095854 | -0.227179 to -0.154682 |
| terminator;771 | aldo/keto reductase | chromosome:1390770-1391603(+) | \|\|\|chromosome:1390834-1390835(-):+0.276819[0.4651163,0.7419355] | | | | | | | | | -0.437747 to -0.437747 | +0.552265 to +0.552265 | +0.391843 to +0.391843 |
| terminator;784 | proline dehydrogenase family protein | chromosome:1408605-1409606(-) | \|\|\|chromosome:1409522-1409523(+):-0.219512[0.2195122,0] | | | chromosome:1409529-1409530(-):+0.256966[0.05882353,0.31578946] | | | | | | -0.950956 to -0.950956 | +0.527425 to +0.527425 | +0.634709 to +0.634709 |
| terminator;787 | class I SAM-dependent methyltransferase;TIGR01212 family radical SAM protein | chromosome:1412767-1414280(-) | | | | chromosome:1414158-1414159(-):+0.208064[0.05,0.2580645] | | | | | | -0.50648 to -0.430842 | +0.216931 to +0.762577 | +0.124736 to +0.404513 |
| terminator;788 | MDR family MFS transporter | chromosome:1414390-1415571(+) | | | | chromosome:1414820-1414821(+):+0.3[0.3,0.6] | | | \|\|\|chromosome:1414764-1414765(-):-0.230519[0.32142857,0.09090909] | | | -0.00213862 to -0.00213862 | -1.04297 to -1.04297 | +0.439513 to +0.439513 |
| terminator;807 | acetate--CoA ligase | chromosome:1452551-1454257(+) | | | | | | | \|\|\|chromosome:1453942-1453943(-):+0.203911[0.33846155,0.5423729];chromosome:1454124-1454125(-):+0.298693[0.32352942,0.62222224] | | | -0.376642 to -0.376642 | +1.18894 to +1.18894 | +0.615328 to +0.615328 |

| | | | | | | | | | | | | | | |
|---|---|---|---|---|---|---|---|---|---|---|---|---|---|---|
| terminator;808 | formate--tetrahydrofolate ligase | chromosome:1454697-1456364(+) | chromosome:1454892-1454893(+):-0.238095[0.5714286,0.33333334] | | | | | | | | | -0.568768 to -0.568768 | -0.638418 to -0.638418 | +0.05022 to +0.05022 |
| terminator;85 | acyl CoA:acetate/3-ketoacid CoA transferase;class I adenylate-forming enzyme family protein;acyl-CoA dehydrogenase family protein;3-hydroxyacyl-CoA dehydrogenase/enoyl-CoA hydratase family protein;acetyl-CoA C-acyltransferase | chromosome:153539-161631(+) | | | | chromosome:161591-161592(+):+0.324357[0.21052632,0.53488374] | | | \|\|\|chromosome:159266-159267(-):+0.288462[0.25,0.53846157] | | | +0.0206173 to +1.2322 | -0.260428 to +1.47364 | -1.59066 to -0.0977758 |
| terminator;851 | NUDIX domain-containing protein;hypothetical protein;hypothetical protein;trigger factor;ATP-dependent Clp protease ATP-binding subunit ClpX;ribosome biogenesis GTP-binding protein YihA/YsxC | chromosome:1524458-1529635(+) | \|\|\|chromosome:1528300-1528301(-):+0.289744[0.07692308,0.36666667] | | | | | | | | | -0.476173 to +0.11073 | -1.22587 to +0.0621545 | +0.0639271 to +0.385384 |

| | | | | | | | | | | | | | | |
|---|---|---|---|---|---|---|---|---|---|---|---|---|---|---|
| terminator;852 | glutamyl-tRNA reductase;cytochrome c biogenesis protein CcsA;hydroxymethylbilane synthase;uroporphyrinogen-III synthase;porphobilinogen synthase;glutamate-1-semialdehyde 2%2C1-aminomutase;hypothetical protein;AbrB family transcriptional regulator | chromosome:1529852-1537273(+) | chromosome:1530878-1530879(+):+0.270833[0.0625,0.33333334] | | | | | | | | | -0.33397 to +0.245317 | -1.11723 to +0.503021 | -0.0315608 to +0.565152 |
| terminator;855 | A24 family peptidase;DNA repair protein RadC | chromosome:1542643-1544033(+) | \|\|\|chromosome:1543880-1543881(-):-0.2174[0.3580247,0.140625] | | | | | | | | | +0.958683 to +2.19853 | +1.99663 to +2.09724 | -1.16471 to -0.0339513 |
| terminator;862 | single-stranded-DNA-specific exonuclease RecJ;adenine phosphoribosyltransferase | chromosome:1557186-1559999(+) | | | | | | | chromosome:1557528-1557529(+):+0.227848[0.1392405,0.36708862] | | | -0.0491189 to +0.0584542 | -1.30913 to -1.03821 | +0.0459915 to +0.122829 |
| terminator;863 | bifunctional (p)ppGpp synthetase/guanosine-3'%2C5'-bis(diphosphate) 3'-pyrophosphohydrolase;D-aminoacyl-tRNA deacylase;N-acetylmuramoyl-L-alanine amidase | chromosome:1560427-1563952(+) | chromosome:1561663-1561664(+):+0.215311[0.36363637,0.57894737] | | | | | | | | | -0.52057 to -0.0603255 | +0.165869 to +0.587217 | -0.328241 to -0.152662 |
| terminator;870 | cysteine desulfurase family protein;tRNA 2-thiouridine(34) synthase MnmA | chromosome:1572663-1574924(+) | \|\|\|chromosome:1572777-1572778(-):-0.402597[0.54545456,0.14285715] | | | | | | | | | -0.263523 to +0.309217 | -0.485645 to -0.131749 | +0.481485 to +0.603901 |

| | | | | | | | | | | | | | | |
|---|---|---|---|---|---|---|---|---|---|---|---|---|---|---|
| terminator;872 | alanine--tRNA ligase;IreB family regulatory phosphoprotein; Holliday junction resolvase RuvX;DUF1292 domain-containing protein | chromosome:1578876-1582584(+) | chromosome:1581419-1581420(+):-0.233987[0.4117647,0.17777778] | | | | | | | | | -0.527808 to -0.227251 | -0.251743 to +0.0782066 | -0.0818103 to +0.347647 |
| terminator;888 | DEAD/DEAH box helicase;deoxyribonuclease IV | chromosome:1634291-1636537(+) | | | | | | | \|\|\|chromosome:1635283-1635284(-):-0.240628[0.48387095,0.24324325] | | | -0.407994 to -0.392111 | -0.677187 to -0.670775 | +0.254574 to +0.41321 |
| terminator;891 | peptidoglycan D%2CD-transpeptidase FtsI family protein | chromosome:1639747-1641822(+) | | chromosome:1641621-1641622(+):-0.260613[0.5625,0.3018868] | | | | | | | | -0.177445 to -0.177445 | +0.585889 to +0.585889 | -0.358407 to -0.358407 |
| terminator;895 | hypothetical protein;shikimate kinase;glycine cleavage system aminomethyltransferase GcvT;aminomethyl-transferring glycine dehydrogenase subunit GcvPA;aminomethyl-transferring glycine dehydrogenase subunit GcvPB | chromosome:1650038-1654779(+) | | | | | | | \|\|\|chromosome:1654143-1654144(-):+0.200723[0.2278481,0.42857143] | | | -0.565923 to +0.440049 | +0.189276 to +1.55953 | -0.339362 to +1.22273 |

| | | | | | | | | | | | | | | |
|---|---|---|---|---|---|---|---|---|---|---|---|---|---|---|
| terminator;900 | acetyl-CoA carboxylase biotin carboxyl carrier protein;acetyl-CoA carboxylase biotin carboxylase subunit;Asp23/Gls24 family envelope stress response protein;transcription antitermination factor NusB;exodeoxyribonuclease VII large subunit;exodeoxyribonuclease VII small subunit;polyprenyl synthetase family protein | chromosome:1659618-1664699(+) | chromosome:1660851-1660852(+):+0.370924[0.28125,0.65217394] | \|\|\|chromosome:1661937-1661938(-):+0.21627[0.3392857,0.5555556] | | | | | | | | -0.520912 to -0.127282 | -0.776937 to +0.214976 | -0.314628 to +0.264361 |
| terminator;910 | pyrroline-5-carboxylate reductase | chromosome:1684555-1685370(-) | | | | \|\|\|chromosome:1684855-1684856(+):-0.47[0.72,0.25] | | | | | | -0.512315 to -0.512315 | +1.59056 to +1.59056 | -0.137004 to -0.137004 |
| terminator;914 | Fur family transcriptional regulator;site-specific tyrosine recombinase XerD | chromosome:1688384-1689769(+) | | | | | | | chromosome:1688625-1688626(+):-0.232637[0.57746476,0.3448276] | | | -0.232368 to +0.24569 | -1.12091 to -0.718151 | -0.78984 to -0.679892 |

| | | | | | | | | | | | | | | |
|---|---|---|---|---|---|---|---|---|---|---|---|---|---|---|
| terminator;935 | Ig domain-containing protein;hypothetical protein;hypothetical protein;phage tail tape measure protein;phage tail family protein;prophage endopeptidase tail family protein;hypothetical protein;hypothetical protein;BppU family phage baseplate upper protein;DUF2977 domain-containing protein;XkdX family protein;DUF2951 domain-containing protein | chromosome:1725559-1739825(+) | chromosome:1727702-1727703(+):+0.242529[0.033333335,0.27586207];chromosome:1731106-1731107(+):+0.226891[0.05882353,0.2857143] | | chromosome:1730477-1730478(+):-0.318253[0.46341464,0.14516129];chromosome:1732665-1732666(+):-0.20097[0.2820513,0.08108108];chromosome:1733575-1733576(+):+0.200397[0.17460318,0.375]\|\|\|chromosome:1731665-1731666(-):+0.20979[0.15384616,0.36363637] | \|\|\|chromosome:1731156-1731157(-):-0.21118[0.42857143,0.2173913];chromosome:1731497-1731498(-):-0.238095[0.32142857,0.083333336];chromosome:1731666-1731667(-):-0.235294[0.3529412,0.11764706] | | | | | | -0.052352 to +1.37448 | -1.75397 to +2.41122 | -1.60349 to +1.51548 |
| terminator;954 | chorismate synthase;3-dehydroquinate synthase;3-phosphoshikimate 1-carboxyvinyltransferase;tetratricopeptide repeat protein;YpiB family protein;DUF1405 domain-containing protein | chromosome:1765251-1771232(+) | \|\|\|chromosome:1765339-1765340(-):+0.212197[0.10526316,0.31746033] | | | | | | | | | -0.461702 to -0.257459 | +0.182579 to +0.492537 | +0.223073 to +0.606035 |
| terminator;957 | N-acetyl-alpha-D-glucosaminyl L-malate synthase BshA;CCA tRNA nucleotidyltransferase;biotin--[acetyl-CoA-carboxylase] ligase;ATP-dependent DNA helicase DinG | chromosome:1772879-1778903(+) | chromosome:1774558-1774559(+):+0.213439[0.09090909,0.3043478] | | | | | | | | | -0.451169 to -0.149075 | -0.329609 to -0.0940541 | +0.10085 to +0.369858 |

| | | | | | | | | | | | | | | |
|---|---|---|---|---|---|---|---|---|---|---|---|---|---|---|
| terminator;958 | asparagine--tRNA ligase | chromosome:1779225-1780517(+) | | | | \|\|\|chromosome:1779882-1779883(-):-0.217334[0.3392857,0.12195122] | | | | | | -0.326604 to -0.326604 | -0.322656 to -0.322656 | +0.117261 to +0.117261 |
| terminator;959 | DnaD domain-containing protein;endonuclease III;YpoC family protein | chromosome:1780845-1782526(+) | | | \|\|\|chromosome:1781135-1781136(-):-0.202369[0.7702703,0.56790125] | | | | | | | -0.522227 to -0.394863 | -0.793421 to -0.55116 | +0.30254 to +0.710494 |
| terminator;960 | transglycosylase domain-containing protein;Holliday junction resolvase RecU | chromosome:1783092-1785898(-) | chromosome:1785060-1785061(-):+0.275862[0,0.27586207] | | | | | | | | | -0.309815 to -0.1177 | +1.92901 to +2.15876 | +0.158938 to +0.329484 |
| terminator;964 | PepSY-associated TM helix domain-containing protein | chromosome:1790120-1791460(-) | chromosome:1790716-1790717(-):+0.225624[0.1632653,0.3888889] | | | | | | | | | +0.355168 to +0.355168 | -1.24167 to -1.24167 | +1.60591 to +1.60591 |
| terminator;967 | multidrug efflux MFS transporter NorB | chromosome:1800340-1801731(+) | | | | chromosome:1800785-1800786(+):+0.222222[0,0.22222222] | | | | | | +0.0327545 to +0.0327545 | -0.409981 to -0.409981 | -0.306965 to -0.306965 |
| terminator;968 | hyperosmolarity resistance protein Ebh | chromosome:1802129-1833394(+) | | | | \|\|\|chromosome:1808095-1808096(-):-0.22619[0.39285713,0.16666667] | | | | | | +0.250215 to +0.250215 | +0.0403769 to +0.0403769 | +0.141976 to +0.141976 |
| terminator;970 | hypothetical protein | chromosome:1834199-1834330(+) | chromosome:1834204-1834205(+):+0.315942[0.2173913,0.53333336] | | | | | | chromosome:1834206-1834207(+):-0.204503[0.5121951,0.30769232] | | | | | |
| terminator;981 | 2-oxoglutarate dehydrogenase E1 component;dihydrolipoyllysine-residue succinyltransferase | chromosome:1850231-1854311(+) | \|\|\|chromosome:1851244-1851245(-):-0.227273[0.5,0.27272728] | chromosome:1852942-1852943(+):-0.260163[0.33333334,0.07317073] | | | | | \|\|\|chromosome:1853539-1853540(-):+0.221269[0.41509435,0.6363636] | | | -0.610049 to -0.560094 | -0.079993 to +0.0445536 | -0.287981 to +0.263047 |
| terminator;983 | MoxR family ATPase;hypothetical protein | chromosome:1856113-1858804(+) | \|\|\|chromosome:1857086-1857087(-):-0.308482[0.5294118,0.22093023] | | | | | | | | | -0.591167 to -0.550428 | -0.458075 to -0.344843 | +0.433649 to +0.620302 |

| | | | | | | | | | | | | | | |
|---|---|---|---|---|---|---|---|---|---|---|---|---|---|---|
| terminator;992 | phosphate ABC transporter substrate-binding protein PstS;phosphate ABC transporter permease subunit PstC;phosphate ABC transporter permease PstA;phosphate ABC transporter ATP-binding protein PstB;phosphate signaling complex protein PhoU | chromosome:1877883-1882448(+) | | | | | | | chromosome:1877950-1877951(+):-0.200658[0.9375,0.7368421] | | | +1.00105 to +1.97702 | +1.34657 to +3.08954 | -0.936741 to +0.0120532 |
| terminator;995 | nickel ABC transporter permease;nickel transporter permease;ATP-binding cassette domain-containing protein;dipeptide/oligopeptide/nickel ABC transporter ATP-binding protein | chromosome:1885255-1888518(+) | chromosome:1886891-1886892(+):+0.289474[0.5,0.7894737]\|\|\|chromosome:1886897-1886898(-):-0.205128[0.20512821,0] | | | | | | | | | -0.10336 to +0.520769 | +0.775135 to +1.71068 | +0.241288 to +0.631201 |
| terminator;999 | tryptophan synthase subunit alpha;tryptophan synthase subunit beta;phosphoribosylanthranilate isomerase;indole-3-glycerol phosphate synthase TrpC;anthranilate phosphoribosyltransferase;aminodeoxychorismate/anthranilate synthase component II;anthranilate synthase component I | chromosome:1893481-1899798(-) | | | | | | | chromosome:1897587-1897588(-):+0.214912[0.36842105,0.5833333] | | | -0.228261 to +0.346296 | -0.569986 to +0.717822 | -1.19647 to -0.457384 |

<table>
<tr><td>upstream;1013</td><td>aconitate hydratase AcnA</td><td>chromosome:1923074-1923774(-)</td><td>|||chromosome:1923543-1923544(+):-0.278571[0.31428573,0.035714287]</td><td></td><td></td><td>|||chromosome:1923421-1923422(+):+0.237879[0.05,0.28787878]</td><td></td><td></td><td></td><td></td><td></td><td>-0.332158 to -0.332158</td><td>+0.352215 to +0.352215</td><td>-0.26488 to -0.26488</td></tr>
<tr><td>upstream;1047</td><td>hypothetical protein</td><td>chromosome:1965594-1966087(-)</td><td>chromosome:1965751-1965752(-):+0.219444[0.225,0.44444445]</td><td></td><td></td><td></td><td></td><td></td><td></td><td></td><td></td><td>-0.52214 to -0.52214</td><td>+0.897681 to +0.897681</td><td>-0.97493 to -0.97493</td></tr>
<tr><td>upstream;1058</td><td>hypothetical protein</td><td>chromosome:1989844-1990544(+)</td><td>chromosome:1990094-1990095(+):-0.236928[0.4722222,0.23529412]</td><td></td><td></td><td></td><td></td><td></td><td></td><td></td><td></td><td>-0.402202 to -0.402202</td><td>-0.175057 to -0.175057</td><td>-0.765249 to -0.765249</td></tr>
<tr><td>upstream;1083</td><td>RluA family pseudouridine synthase;signal peptidase II</td><td>chromosome:2098464-2099164(-)</td><td>chromosome:2098834-2098835(-):-0.396104[0.71428573,0.3181818]|||chromosome:2098826-2098827(+):-0.217949[0.8333333,0.61538464]</td><td></td><td></td><td>chromosome:2098878-2098879(-):+0.20597[0.6,0.80597013]</td><td></td><td></td><td></td><td></td><td></td><td>-0.507287 to -0.0498163</td><td>-1.12818 to -0.751374</td><td>+0.343275 to +0.417111</td></tr>
<tr><td>upstream;1086</td><td>cell division protein FtsZ;cell division protein FtsA;cell division protein FtsQ/DivIB;UDP-N-acetylmuramoyl-L-alanine--D-glutamate ligase;phospho-N-acetylmuramoyl-pentapeptide-transferase;penicillin-binding protein;cell division protein FtsL;16S rRNA (cytosine(1402)-N(4))-methyltransferase RsmH;division/cell wall cluster transcriptional repressor MraZ</td><td>chromosome:2118162-2118306(-)</td><td>|||chromosome:2118296-2118297(+):-0.209273[0.4473684,0.23809524]</td><td></td><td></td><td></td><td></td><td></td><td></td><td></td><td></td><td>-0.367098 to +0.498964</td><td>-0.19836 to +0.630972</td><td>-1.09948 to -0.0946943</td></tr>
</table>

| | | | | | | | | | | | | | | |
|---|---|---|---|---|---|---|---|---|---|---|---|---|---|---|
| upstream;1090 | beta-class phenol-soluble modulin;beta-class phenol-soluble modulin | chromosome:2121807-2122507(-) | | | | chromosome:2122221-2122222(-):-0.457143[0.6,0.14285715] | | | \|\|\|chromosome:2122232-2122233(+):+0.266667[0.53333336,0.8];chromosome:2122351-2122352(+):+0.272446[0.31578946,0.5882353] | | | -0.0235883 to +1.20067 | +1.24876 to +1.9684 | -3.11836 to -2.62291 |
| upstream;1155 | DUF5011 domain-containing protein | chromosome:2231924-2232624(+) | chromosome:2232283-2232284(+):+0.212308[0.48,0.6923077]\|\|\|chromosome:2232290-2232291(-):-0.290141[0.6,0.30985916] | | | | | | | | | +0.308692 to +0.308692 | +2.12195 to +2.12195 | -0.360905 to -0.360905 |
| upstream;1245 | pathogenicity island protein | chromosome:2397752-2397936(-) | | chromosome:2397864-2397865(-):-0.302953[0.47368422,0.17073171] | | | | | \|\|\|chromosome:2397866-2397867(+):-0.289263[0.82051283,0.53125] | | | +2.79209 to +2.79209 | +0.320948 to +0.320948 | +0.296099 to +0.296099 |
| upstream;1252 | MetQ/NlpA family ABC transporter substrate-binding protein;methionine ABC transporter permease;methionine ABC transporter ATP-binding protein | chromosome:2410903-2411153(-) | \|\|\|chromosome:2411060-2411061(+):+0.412698[0.30952382,0.7222222] | | | | | | chromosome:2411064-2411065(-):-0.231256[0.2682927,0.037037037] | | | -1.03419 to -1.00793 | +0.825532 to +1.17829 | -1.89223 to -0.913467 |
| upstream;126 | bifunctional UDP-sugar hydrolase/5'-nucleotidase;DNA-binding protein | chromosome:265457-266157(-) | chromosome:265972-265973(-):-0.245645[0.31707317,0.071428575] | | | | | | | | | +0.116022 to +2.03236 | +0.036202 to +0.224168 | -0.160551 to +0.0947152 |
| upstream;1287 | DNA-binding protein WhiA;YvcK family protein;RNase adapter RapZ | chromosome:2455159-2455859(-) | \|\|\|chromosome:2455573-2455574(+):-0.230303[0.36363637,0.13333334] | | | chromosome:2455624-2455625(-):-0.323529[0.5,0.1764706] | | | | | | -0.333848 to -0.114227 | -0.15307 to +0.732524 | +0.039034 to +0.141698 |

| | | | | | | | | | | | | | | |
|---|---|---|---|---|---|---|---|---|---|---|---|---|---|---|
| upstream;1288 | thioredoxin-disulfide reductase;tetratricopeptide repeat protein;DapH/DapD/GlmU-related protein;prolipoprotein diacylglyceryl transferase;HPr(Ser) kinase/phosphatase | chromosome:2460758-2461420(-) | \|\|\|chromosome:2461007-2461008(+):-0.319672[0.5,0.18032786] | | | | | | | | | -0.435729 to -0.193286 | -0.748642 to +0.496211 | -0.211857 to +1.12605 |
| upstream;1306 | siderophore ABC transporter substrate-binding protein;ABC transporter ATP-binding protein;iron chelate uptake ABC transporter family permease subunit;ABC transporter permease;hypothetical protein | chromosome:2492362-2492876(-) | | | | | | | \|\|\|chromosome:2492538-2492539(+):+0.310448[0.08955224,0.4];chromosome:2492708-2492709(+):+0.409091[0.09090909,0.5] | | | +0.0710806 to +1.86182 | -1.85978 to -0.662502 | +0.89835 to +2.02544 |
| upstream;1312 | 5'(3')-deoxyribonucleotidase | chromosome:2502817-2503168(-) | chromosome:2503055-2503056(-):-0.4[0.8,0.4] | | | chromosome:2503052-2503053(-):+0.268382[0.4375,0.7058824];chromosome:2503055-2503056(-):+0.232198[0.29411766,0.5263158];chromosome:2503056-2503057(-):+0.260606[0.46666667,0.72727275] | | | | | | +0.0691325 to +0.0691325 | -0.300113 to -0.300113 | -0.0662805 to -0.0662805 |
| upstream;1372 | metal ABC transporter ATP-binding protein;metal ABC transporter permease;metal ABC transporter substrate-binding protein | chromosome:2595494-2596194(+) | chromosome:2595690-2595691(+):+0.203209[0.09090909,0.29411766]\|\|\|chromosome:2595698-2595699(-):-0.203007[0.6315789,0.42857143] | | | | | | | | | -0.717349 to -0.468968 | -0.925436 to -0.698435 | -0.427184 to +0.197481 |

| | | | | | | | | | | | | | | |
|---|---|---|---|---|---|---|---|---|---|---|---|---|---|---|
| upstream;1378 | transposase;IS30 family transposase | chromosome:2617320-2617429(+) | \|\|\|chromosome:2617356-2617357(-):-0.208381[0.7307692,0.52238804] | | | | | | | | | -0.385561 to -0.324211 | +0.297596 to +0.483998 | -0.141847 to -0.0736932 |
| upstream;1385 | endonuclease III domain-containing protein | chromosome:2627824-2628212(-) | | | | \|\|\|chromosome:2627951-2627952(+):+0.331731[0.4375,0.7692308] | | | | | | +0.37189 to +0.37189 | +0.608773 to +0.608773 | -0.200594 to -0.200594 |
| upstream;1458 | LysR family transcriptional regulator | chromosome:2806950-2807650(+) | \|\|\|chromosome:2807322-2807323(-):+0.2125[0.1875,0.4] | | | | | | \|\|\|chromosome:2807316-2807317(-):-0.208333[0.20833333,0];chromosome:2807322-2807323(-):+0.20761[0.17948718,0.38709676] | | | +1.82224 to +1.82224 | +1.79492 to +1.79492 | -0.869766 to -0.869766 |
| upstream;233 | YceI family protein | chromosome:456973-457673(-) | | | | \|\|\|chromosome:457566-457567(+):-0.204678[0.31578946,0.11111111] | | | | | | -0.494641 to -0.494641 | -1.76324 to -1.76324 | +1.22213 to +1.22213 |
| upstream;25 | YeiH family protein | chromosome:35029-35729(+) | | | | chromosome:35409-35410(+):+0.303571[0.525,0.82857144] | | | chromosome:35464-35465(+):+0.216247[0.1521739,0.36842105] | | | -0.00612296 to -0.00612296 | +0.34111 to +0.34111 | -0.268437 to -0.268437 |
| upstream;292 | 3-methyl-2-oxobutanoate hydroxymethyltransferase;pantoate--beta-alanine ligase;aspartate 1-decarboxylase | chromosome:567586-568286(+) | | | | \|\|\|chromosome:568187-568188(-):+0.202797[0.18181819,0.3846154] | | | | | | +0.0642875 to +0.242038 | -0.050995 to +0.598832 | +0.240977 to +0.582526 |
| upstream;298 | quinone-dependent dihydroorotate dehydrogenase | chromosome:576150-576850(-) | chromosome:576647-576648(-):+0.301314[0.046511628,0.3478261] | | | | | \|\|\|chromosome:576580-576581(+):-0.212403[0.27906978,0.06666667] | | | | +0.604281 to +0.604281 | -0.445873 to -0.445873 | +0.941723 to +0.941723 |
| upstream;408 | GyrI-like domain-containing protein | chromosome:743049-743749(+) | | | | \|\|\|chromosome:743481-743482(-):-0.216346[0.5625,0.34615386] | | | | | | +0.622554 to +0.622554 | -0.533336 to -0.533336 | +0.0359044 to +0.0359044 |
| upstream;43 | hypothetical protein | chromosome:69201-69901(+) | | | | | | | chromosome:69775-69776(+):-0.218254[0.8055556,0.5873016] | | | +0.653203 to +0.653203 | +1.09037 to +1.09037 | -1.76922 to -1.76922 |

| | | | | | | | | | | | | | | |
|---|---|---|---|---|---|---|---|---|---|---|---|---|---|---|
| upstream;472 | MOSC domain-containing protein | chromosome:847483-848183(-) | | | | | | | |||chromosome:847965-847966(+):-0.219048[0.26666668,0.04761905] | | | -0.164514 to -0.164514 | -0.0610189 to -0.0610189 | +0.360017 to +0.360017 |
| upstream;480 | Na+/H+ antiporter NhaC family protein | chromosome:859079-859304(+) | | chromosome:859161-859162(+):-0.430769[0.7,0.26923078] | | | | | |||chromosome:859165-859166(-):+0.297297[0.27027026,0.5675676] | | | -0.198089 to -0.198089 | -0.51179 to -0.51179 | -0.0188675 to -0.0188675 |
| upstream;484 | hypothetical protein | chromosome:864642-865342(-) | |||chromosome:865227-865228(+):+0.212821[0.15384616,0.36666667] | | | | | | | | | +0.654338 to +0.654338 | +3.04712 to +3.04712 | -0.00364084 to -0.00364084 |
| upstream;490 | CPBP family intramembrane glutamic endopeptidase SdpB | chromosome:870414-871114(+) | | | | chromosome:870826-870827(+):-0.329114[0.32911393,0] | | | | | | +0.0821259 to +0.0821259 | -0.579765 to -0.579765 | -0.687332 to -0.687332 |
| upstream;520 | HTH-type transcriptional regulator SarV | chromosome:914743-915434(+) | chromosome:914901-914902(+):+0.208238[0.73913044,0.94736844] | | | |||chromosome:914897-914898(-):-0.204082[0.20408164,0] | | | |||chromosome:915011-915012(-):-0.27451[0.9411765,0.6666667] | | | -0.53921 to -0.53921 | -1.42733 to -1.42733 | -0.255953 to -0.255953 |
| upstream;601 | hypothetical protein | chromosome:1083240-1083578(+) | |||chromosome:1083476-1083477(-):-0.204348[0.4,0.19565217] | | | | | | | | | +1.18592 to +1.18592 | +0.122267 to +0.122267 | -0.755793 to -0.755793 |
| upstream;604 | YwpF-like family protein | chromosome:1086384-1087084(-) | | | |||chromosome:1086790-1086791(+):+0.206774[0.15686275,0.36363637] | | | | | | | -0.0362646 to -0.0362646 | -0.021591 to -0.021591 | -0.91358 to -0.91358 |
| upstream;605 | lytic transglycosylase SceD | chromosome:1086662-1087362(+) | | | chromosome:1086790-1086791(+):+0.206774[0.15686275,0.36363637] | | | | | | | +1.13779 to +1.13779 | +0.750479 to +0.750479 | +1.22534 to +1.22534 |

| | | | | | | | | | | | | | | |
|---|---|---|---|---|---|---|---|---|---|---|---|---|---|---|
| upstream;648 | hypothetical protein;exonuclease domain-containing protein;XRE family transcriptional regulator | chromosome:1180278-1180895(-) | chromosome:1180853-1180854(-):+0.221154[0.15384616,0.375]\|\|\|chromosome:1180842-1180843(+):-0.240909[0.75,0.5090909];chromosome:1180845-1180846(+):+0.219958[0.13636364,0.35632184] | | | | | | chromosome:1180853-1180854(-):-0.281513[0.3529412,0.071428575] | | | +0.865604 to +1.30609 | +0.246961 to +0.724232 | -0.726602 to -0.141567 |
| upstream;657 | protein;terminase large subunit;phage portal protein;head maturation protease%2C ClpP-related;phage major capsid protein;hypothetical protein;phage head-tail adapter protein;head-tail adaptor protein;HK97 gp10 family phage protein;hypothetical protein;major tail protein;Ig-like domain-containing protein;hypothetical protein;hypothetical protein;phage tail tape measure protein;phage tail domain- | chromosome:1193697-1193827(+) | \|\|\|chromosome:1193750-1193751(-):+0.31677[0.35714287,0.67391306];chromosome:1193751-1193752(-):+0.216184[0.30555555,0.5217391] | | | | | | \|\|\|chromosome:1193750-1193751(-):+0.459649[0.47368422,0.93333334] | | | +3.2242 to +4.12005 | -0.857005 to +0.263013 | -1.20077 to -0.0132812 |
| upstream;673 | C45 family autoproteolytic acyltransferase/hydolase | chromosome:1231450-1232150(-) | chromosome:1231752-1231753(-):+0.22979[0.15116279,0.3809524] | | | | | | | | | +0.294846 to +0.294846 | +0.809211 to +0.809211 | -0.358874 to -0.358874 |

<table>
<tr><td></td><td></td><td></td><td></td><td></td><td></td><td></td><td></td><td></td><td></td><td></td><td></td><td></td><td></td><td></td></tr>
<tr><td>upstream;674</td><td>hypothetical protein</td><td>chromosome:1231201-1231901(+)</td><td>|||chromosome:1231752-1231753(-):+0.22979[0.15116279,0.3809524]</td><td></td><td></td><td></td><td></td><td></td><td></td><td></td><td></td><td>-0.308327 to -0.308327</td><td>+0.113998 to +0.113998</td><td>-0.000997561 to -0.000997561</td></tr>
<tr><td>upstream;692</td><td>diacylglycerol kinase</td><td>chromosome:1262652-1263235(+)</td><td>chromosome:1263081-1263082(+):-0.251196[0.84210527,0.59090906]</td><td></td><td></td><td></td><td></td><td></td><td></td><td></td><td></td><td>-0.339949 to -0.339949</td><td>-0.0402432 to -0.0402432</td><td>+0.0813331 to +0.0813331</td></tr>
<tr><td>upstream;701</td><td>aromatic acid exporter family protein</td><td>chromosome:1273604-1274304(-)</td><td>chromosome:1273883-1273884(-):+0.373016[0.071428575,0.44444445]</td><td></td><td></td><td>chromosome:1273882-1273883(-):+0.204545[0.54545456,0.75];chromosome:1273883-1273884(-):-0.270588[0.47058824,0.2]</td><td></td><td></td><td>chromosome:1273882-1273883(-):-0.236264[0.8076923,0.5714286]</td><td></td><td></td><td>+0.374092 to +0.374092</td><td>+0.321459 to +0.321459</td><td>-0.263036 to -0.263036</td></tr>
<tr><td>upstream;702</td><td>type I methionyl aminopeptidase</td><td>chromosome:1273432-1274132(+)</td><td>|||chromosome:1273883-1273884(-):+0.373016[0.071428575,0.44444445]</td><td></td><td></td><td>|||chromosome:1273882-1273883(-):+0.204545[0.54545456,0.75];chromosome:1273883-1273884(-):-0.270588[0.47058824,0.2]</td><td></td><td></td><td>|||chromosome:1273882-1273883(-):-0.236264[0.8076923,0.5714286]</td><td></td><td></td><td>-0.480099 to -0.480099</td><td>-0.510743 to -0.510743</td><td>+0.0674126 to +0.0674126</td></tr>
<tr><td>upstream;731</td><td>helix-turn-helix transcriptional regulator</td><td>chromosome:1322395-1322751(+)</td><td></td><td></td><td>chromosome:1322473-1322474(+):-0.308036[0.59375,0.2857143]</td><td></td><td></td><td></td><td></td><td></td><td></td><td>+0.989949 to +0.989949</td><td>-1.33973 to -1.33973</td><td>-0.420171 to -0.420171</td></tr>
<tr><td>upstream;756</td><td>type I toxin-antitoxin system Fst family toxin PepA1</td><td>chromosome:1374286-1374986(-)</td><td>chromosome:1374414-1374415(-):-0.204299[0.24528302,0.040983606]</td><td></td><td></td><td></td><td></td><td></td><td></td><td></td><td></td><td>+0.799098 to +0.799098</td><td>-0.644078 to -0.644078</td><td>-0.442574 to -0.442574</td></tr>
<tr><td>upstream;757</td><td>transposase</td><td>chromosome:1373789-1374489(+)</td><td>|||chromosome:1374414-1374415(-):-0.204299[0.24528302,0.040983606]</td><td></td><td></td><td></td><td></td><td></td><td></td><td></td><td></td><td>-0.283869 to -0.283869</td><td>+0.523982 to +0.523982</td><td>-0.316062 to -0.316062</td></tr>
<tr><td>upstream;767</td><td>IS200/IS605 family transposase</td><td>chromosome:1847022-1847722(-)</td><td></td><td>chromosome:1847121-1847122(-):-0.24539[0.26666668,0.021276595]</td><td></td><td></td><td></td><td></td><td></td><td></td><td></td><td>+0.307815 to +0.307815</td><td>-1.48515 to -1.48515</td><td>+0.280622 to +0.280622</td></tr>
</table>

| | | | | | | | | | | | | | | |
|---|---|---|---|---|---|---|---|---|---|---|---|---|---|---|
| upstream;768 | phosphoenolpyruvate carboxykinase (ATP) | chromosome:1387961-1388661(-) | | | | chromosome:1388151-1388152(-):+0.225694[0.21875,0.44444445] | | | | | | -0.709471 to -0.709471 | -0.660969 to -0.660969 | +0.490619 to +0.490619 |
| upstream;769 | methionine adenosyltransferase | chromosome:1387632-1388332(+) | | | | \|\|\|chromosome:1388151-1388152(-):+0.225694[0.21875,0.44444445] | | | | | | -0.415291 to -0.415291 | -0.792243 to -0.792243 | -0.287849 to -0.287849 |
| upstream;785 | hypothetical protein;alpha/beta hydrolase | chromosome:1409018-1409718(+) | chromosome:1409522-1409523(+):-0.219512[0.2195122,0] | | | \|\|\|chromosome:1409529-1409530(-):+0.256966[0.05882353,0.31578946] | | | | | | -0.292211 to +0.0835537 | -0.801728 to +0.0133029 | -0.615414 to +0.496183 |
| upstream;787 | class I SAM-dependent methyltransferase;TIGR01212 family radical SAM protein | chromosome:1414280-1414980(-) | | | | \|\|\|chromosome:1414820-1414821(+):+0.3[0.3,0.6] | | | chromosome:1414764-1414765(-):-0.230519[0.32142857,0.09090909] | | | -0.50648 to -0.430842 | +0.216931 to +0.762577 | +0.124736 to +0.404513 |
| upstream;788 | MDR family MFS transporter | chromosome:1413690-1414390(+) | | | | \|\|\|chromosome:1414158-1414159(-):+0.208064[0.05,0.2580645] | | | | | | -0.00213862 to -0.00213862 | -1.04297 to -1.04297 | +0.439513 to +0.439513 |
| upstream;793 | Mn(2+)-dependent dipeptidase Sapep;D-amino-acid transaminase | chromosome:1430129-1430829(+) | \|\|\|chromosome:1430572-1430573(-):+0.4[0.3,0.7] | | | \|\|\|chromosome:1430157-1430158(-):+0.266667[0.6,0.8666667] | | | chromosome:1430569-1430570(+):+0.352564[0.23076923,0.5833333];chromosome:1430577-1430578(+):+0.254945[0.51428574,0.7692308]\|\|\|chromosome:1430571-1430572(-):+0.206767[0.57894737,0.78571427];chromosome:1430573-1430574(-):-0.234756[0.6097561,0.375] | | | -0.66712 to -0.520942 | +0.349521 to +0.463577 | -0.499333 to -0.313378 |
| upstream;836 | pApA hydrolase Pde2;DNA polymerase III subunit alpha | chromosome:1487688-1487949(+) | chromosome:1487741-1487742(+):+0.208333[0.125,0.33333334] | | | | | | | | | -0.490123 to -0.48341 | -0.148647 to +0.0469507 | -0.0764602 to -0.0456154 |

| | | | | | | | | | | | | | | |
|---|---|---|---|---|---|---|---|---|---|---|---|---|---|---|
| upstream;848 | threonine--tRNA ligase | chromosome:1518874-1519072(+) | | | | chromosome:1518890-1518891(+):+0.226144[0.10280374,0.32894737] | | | | | | -0.56537 to -0.56537 | -0.36733 to -0.36733 | +0.463234 to +0.463234 |
| upstream;869 | LLM class flavin-dependent oxidoreductase | chromosome:1572351-1573051(-) | chromosome:1572777-1572778(-):-0.402597[0.54545456,0.14285715] | | | | | | | | | -0.679537 to -0.679537 | -2.54147 to -2.54147 | +1.89307 to +1.89307 |
| upstream;885 | glycine--tRNA ligase | chromosome:1627289-1627989(-) | | | | | | | chromosome:1627322-1627323(-):+0.220196[0.2580645,0.47826087] | | | -0.446343 to -0.446343 | +0.110584 to +0.110584 | +0.17359 to +0.17359 |
| upstream;886 | helix-turn-helix transcriptional regulator;pyruvate%2C water dikinase regulatory protein;DNA primase;RNA polymerase sigma factor RpoD | chromosome:1626924-1627624(+) | | | | | | | \|\|\|chromosome:1627322-1627323(-):+0.220196[0.2580645,0.47826087] | | | -0.443859 to +0.0941541 | -0.218755 to +0.618995 | -0.220279 to +0.229226 |
| upstream;911 | SDR family NAD(P)-dependent oxidoreductase;hypothetical protein | chromosome:1684812-1685512(+) | | | | chromosome:1684855-1684856(+):-0.47[0.72,0.25] | | | | | | +0.224446 to +0.33544 | +0.764847 to +1.1186 | -0.351664 to -0.0956352 |
| upstream;933 | P27 family phage terminase small subunit;terminase TerL endonuclease subunit;phage portal protein;head maturation protease%2C ClpP-related;phage major capsid protein | chromosome:1718018-1718145(+) | | | | \|\|\|chromosome:1718089-1718090(-):+0.229167[0.33333334,0.5625] | | | | | | +0.45656 to +1.26587 | +0.214863 to +1.52277 | -1.94031 to +0.498661 |
| upstream;969 | ribonuclease HI family protein | chromosome:1833854-1834395(-) | \|\|\|chromosome:1834204-1834205(+):+0.315942[0.2173913,0.53333336] | | | | | | \|\|\|chromosome:1834206-1834207(+):-0.204503[0.5121951,0.30769232] | | | -0.230523 to -0.230523 | +0.0406014 to +0.0406014 | -0.477134 to -0.477134 |

| | | | | | | | | | | | | | | |
|---|---|---|---|---|---|---|---|---|---|---|---|---|---|---|
| upstream;978 | hypothetical protein | chromosome:1846566-1847266(+) | | \|\|\|chromosome:1847121-1847122(-):-0.24539[0.26666668,0.021276595] | | | | | | | | +0.177745 to +0.177745 | +0.232164 to +0.232164 | +1.91293 to +1.91293 |
| upstream;992 | phosphate ABC transporter substrate-binding protein PstS;phosphate ABC transporter permease subunit PstC;phosphate ABC transporter permease PstA;phosphate ABC transporter ATP-binding protein PstB;phosphate signaling complex protein PhoU | chromosome:1877183-1877883(+) | \|\|\|chromosome:1877389-1877390(-):-0.366071[0.9285714,0.5625] | | | chromosome:1877418-1877419(+):+0.219231[0.05,0.26923078] | | | | | | +1.00105 to +1.97702 | +1.34657 to +3.08954 | -0.936741 to +0.0120532 |
