## Supplemental Figs 1 and 2 for "Stress-induced DNA methylome plasticity and transcriptional re-programming in *Staphylococcus aureus*"

**Supplementary Data**

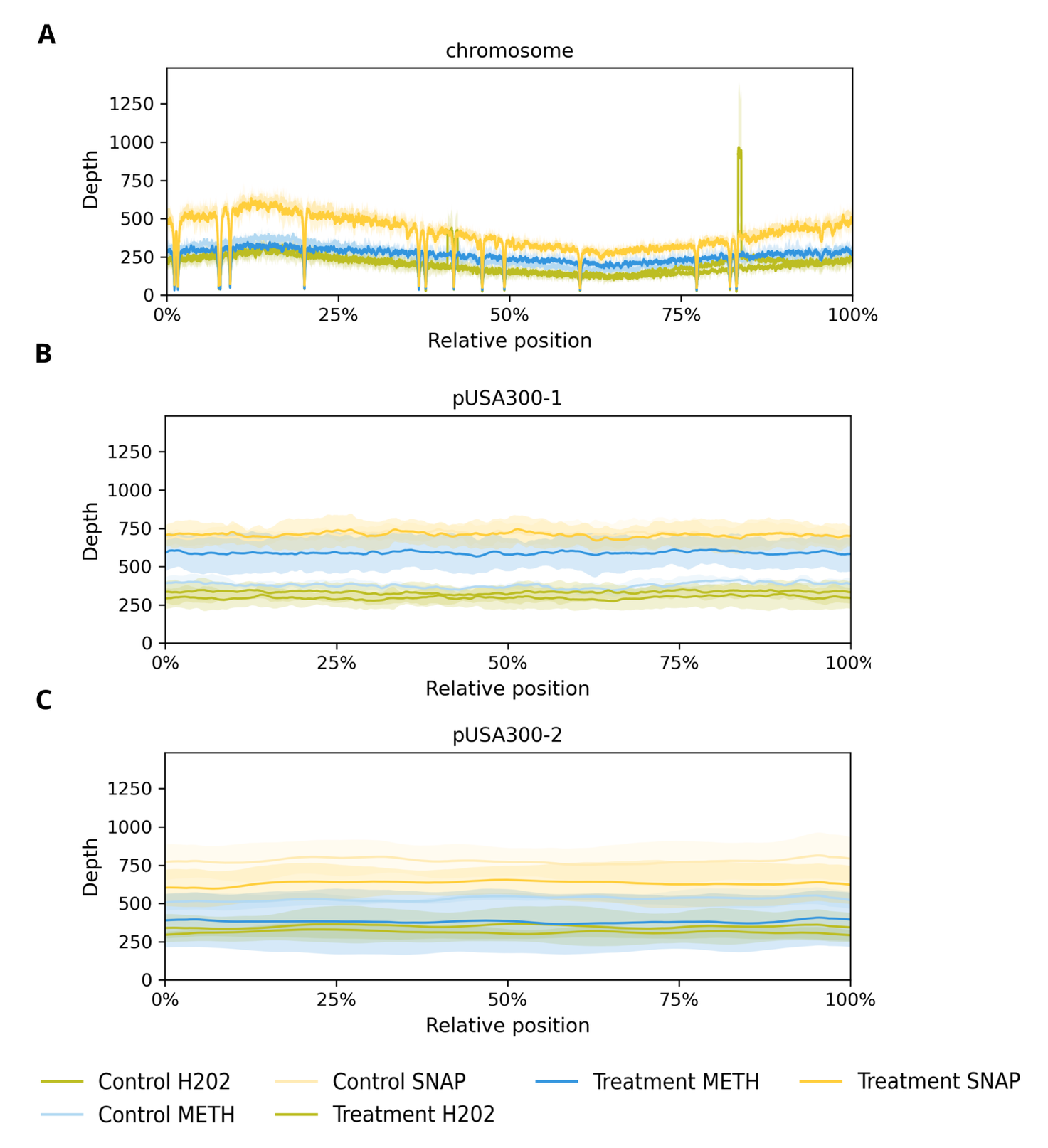
**Supplementary Figure 1. Assembly coverage map of reads used in base level methylation calling. Sample coverage data are plotted *per* contig: chromosome and plasmids pUSA300-1 and pUSA300-2.**

**Explanation for superficial differences between the volcano plots (Figure 1 in main text) and the GSEA results (Figure 2 in main text)**

**The superficial differences between the volcano plots and the GSEA results arise because the two analyses measure different things. Volcano plots show individual genes that pass predefined significance and fold-change thresholds, whereas GSEA uses the complete list of genes ranked by log₂FC and tests whether members of a pathway (assigned by GO tags) tend to occur toward one end of that ranking. A pathway can therefore show significant negative enrichment when many of its genes undergo small, coordinated decreases, even if few individual genes are significantly downregulated.** This explains the H₂O₂ response. Although only three genes were significantly downregulated in the volcano plot, KEGG orthology-assigned genes were over-represented among genes with negative log₂FC values (R = 1.18) and under-represented among genes with positive log₂FC values (R = 0.78) (Supplementary Figure 2). Thus, many functionally annotated genes showed modest negative shifts that collectively produced predominantly negative pathway enrichment, while a smaller number of genes showed strong, individually significant induction. Following SNAP treatment, KEGG orthology-assigned genes showed the opposite overall directional bias, being under-represented among genes with negative log₂FC values (R = 0.86) and over-represented among genes with positive log₂FC values (R = 1.15). Nevertheless, more negatively enriched pathway terms were detected because many of the same negatively shifted genes belonged to several overlapping metabolic pathways and therefore contributed to multiple enriched terms. Positive shifts were concentrated within fewer pathways. Accordingly, neither the number of enriched pathways nor bubble size should be interpreted as the number of significantly upregulated or downregulated genes. The volcano plots describe significant changes in individual genes, whereas the GSEA plots describe coordinated shifts in pathway-associated gene rankings; the two figures are therefore complementary rather than contradictory.

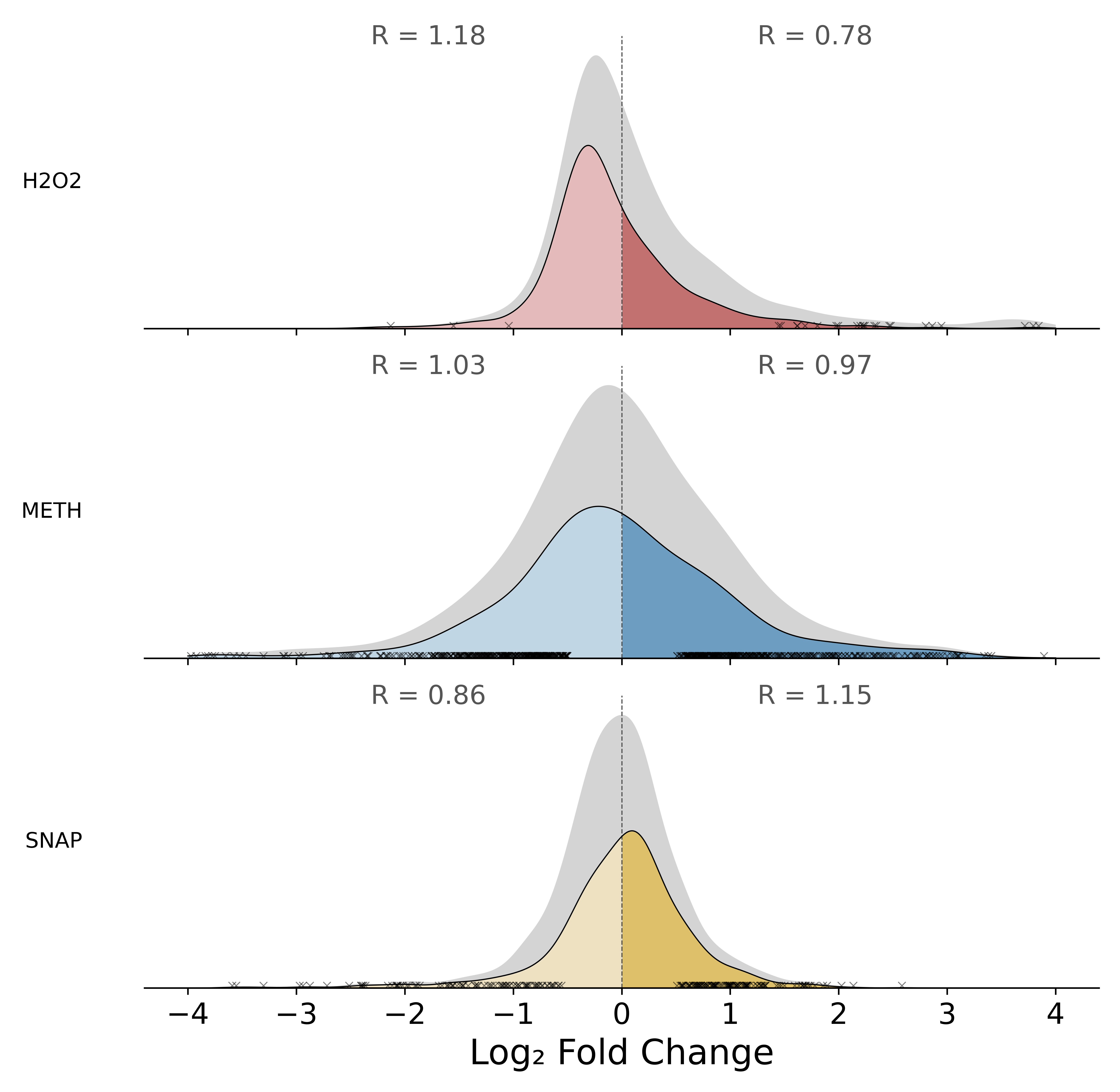
**Supplementary Figure 2. Distribution of GO annotated genes relative to expression change. Ridge density plots of log₂FC distributions for genes with** **KEGG orthology assignments, under each treatment. Coloured areas represent** **KEGG orthology-assigned genes; grey background areas represent all genes. Vertical dashed line marks log₂FC = 0. Black ‘x’ markers indicate KEGG orthology-assigned genes that passed significance thresholds (|log₂FC| > 0.5 and p < 0.05). R values report the directional enrichment ratio, calculated as R = (k / K) ÷ (g / G) (where: k = number of upregulated/downregulated KO-assigned genes; K = total number of KO-assigned genes; g = number of genes being upregulated/downregulated; G = total number of genes), such that R > 1 indicates enrichment and R < 1 indicates depletion of KEGG orthology genes on that side (upregulated or downregulated). Each subplot uses its own y-axis scale (area ∝ gene count), allowing the absolute heights of the grey (all genes) and coloured (KEGG orthology genes) ridges to be visually distinguished.**

**Supplementary Table 1. Genome-wide mapping of base-level significant differential methylation events across treatments and annotated genomic regions. This table summarises all significant methylation sites and their overlap with annotated genomic features, integrating positional information, strand orientation, and changes in methylation within a single tabular format. Each row represents one annotated genomic region (gene, upstream, or terminator; shown in column one). All rows included contain at least one significant methylation change event. Methylation events are categorised by treatment and methylation type (hydrogen peroxide = H₂O₂_4mC/5mC, H₂O₂_6mA; methicillin = METH_4mC/5mC, METH_6mA; SNAP = SNAP_4mC/5mC, SNAP_6mA). Within each treatment-specific cell, sites listed before ‘|||’ are located on the coding strand relative to the genomic feature, whereas those listed after ‘|||’ occur on the opposing strand. Sites are reported in the format ‘contig:start coordinate–end coordinate (strand): change in methylation’ (e.g. “chromosome:2223–2224(+):−0.45”, indicating a chromosomal methylation decrease of 45%). Subsequent columns report log₂ fold change (log₂FC) values associated with these regions. For gene coding regions, the log₂FC shown in the three treatment columns corresponds to gene expression changes. For upstream and terminator regions, the log₂FC value or range provided for each treatment corresponds to the genes encoded within the operon to which the associated genomic feature belongs.**
